# Defective heart chamber growth and myofibrillogenesis after knockout of *adprhl1* gene function by targeted disruption of the ancestral catalytic active site

**DOI:** 10.1101/2020.02.13.947424

**Authors:** Stuart J. Smith, Norma Towers, Kim Demetriou, Timothy J. Mohun

## Abstract

ADP-ribosylhydrolase-like 1 (Adprhl1) is a pseudoenzyme expressed in the developing heart myocardium of all vertebrates. In the amphibian *Xenopus laevis*, knockdown of the two cardiac Adprhl1 protein species (40 and 23 kDa) causes failure of chamber outgrowth but this has only been demonstrated using antisense morpholinos that interfere with RNA-splicing. Transgenic production of 40 kDa Adprhl1 provides only part rescue of these defects. CRISPR/Cas9 technology now enables targeted mutation of the *adprhl1* gene in G0-generation embryos with routine cleavage of all alleles. Testing multiple gRNAs distributed across the locus reveals exonic locations that encode critical amino acids for Adprhl1 function. The gRNA recording the highest frequency of a specific ventricle outgrowth phenotype directs Cas9 cleavage of an exon 6 sequence, where microhomology mediated end-joining biases subsequent DNA repairs towards three small in-frame deletions. Mutant alleles encode discrete loss of 1, 3 or 4 amino acids from a di-arginine (Arg271-Arg272) containing peptide loop at the centre of the ancestral ADP-ribosylhydrolase site. Thus despite lacking catalytic activity, it is the modified (adenosine-ribose) substrate binding cleft of Adprhl1 that fulfils an essential role during heart formation. Mutation results in striking loss of myofibril assembly in ventricle cardiomyocytes. The defects suggest Adprhl1 participation from the earliest stage of cardiac myofibrillogenesis and are consistent with previous MO results and Adprhl1 protein localization to actin filament Z-disc boundaries. A single nucleotide change to the gRNA sequence renders it inactive. Mice lacking *Adprhl1* exons 3-4 are normal but production of the smaller ADPRHL1 species is unaffected, providing further evidence that cardiac activity is concentrated at the C-terminal protein portion.

**Highlights:** Comparison of *adprhl1* morpholinos. Knockdown of the two *Xenopus* cardiac Adprhl1 protein species (40 and 23 kDa) causes failure of ventricle outgrowth.

CRISPR/Cas9 targeted gene mutation of *adprhl1* with multiple gRNAs reveals exonic locations that encode critical amino acids for Adprhl1 function.

Repair of DSBs at exon 6 yields small in-frame deletions that cause specific ventricle myofibril assembly defects.

The deletions disturb a conserved di-arginine containing peptide loop at the centre of the ancestral substrate binding cleft/ADP-ribosylhydrolase site of this pseudoenzyme.

Mice lacking *Adprhl1* exons 3-4 are normal but production of the smaller ADPRHL1 species is unaffected, providing further evidence that cardiac activity is concentrated at the C-terminal protein portion.

## Introduction

The embryonic heart forms as a simple, linear muscle tube that subsequently loops and is transformed by growth of the cardiac chambers that balloon out from regions of the tube’s outer curvature (Reviewed Boogerd et al., 2009; Desgrange et al., 2018). At the level of the cardiomyocytes, chamber outgrowth occurs via an increase in cell size and coordinated changes in cell shape and directionality (Lin et al., 2012). This is augmented by a localized resumption of cell proliferation and the continued addition of newly differentiated tissue to the ends of the tube (the heart poles) (Kelly et al., 2014; Uribe et al., 2018). Directed hypertrophy determines the overall shape of a chamber while cell division is more evident during ballooning stages in mammalian hearts that undergo rapid increases in size (Günthel et al., 2018). Outgrowth occurs as the linear heart has already commenced peristaltic beating and zebrafish studies have shown that blood fluid forces can influence final chamber morphology by affecting cytoskeletal protein localization, cardiomyocyte maturation and also endocardial proliferation (Bornhorst et al., 2019; Fukuda et al., 2019; Reviewed Sidhwani and Yelon, 2019). Underpinning the directed growth is myofibrillogenesis, the process of assembling the contractile protein machinery within the muscle cells.

Aquatic vertebrate species have two-chambered (fish) or three-chambered (amphibia) hearts featuring a single ventricle. Growth of the large ventricle within *Xenopus* tadpoles is sustained over a two month larval period, although the initial acquisition of form occurs within four days. It commences as a group of cardiomyocytes elongate within the heart tube and transiently align in a rosette pattern (Smith et al., 2016). This rearrangement is biased to the left side, breaks the left-right symmetry and defines the position of the chamber apex. The chamber cells assemble myofibrils whose predominant trajectory will extend across the ventricle width. Their direction of muscle filament extension is essentially perpendicular with respect to circumference-axes imagined running from inner to outer curvature (or perpendicular to concentric lines encircling the primitive tube). As the ventricle shape develops, that alignment becomes perpendicular with regard base to chamber apex axes. For cardiomyocytes positioned further away from the apex, the angle of myofibril production is shifted progressively towards a parallel to circumference axes direction. As a consequence of the directed growth, the anterior wall of the ventricle has a left-sided origin, the posterior wall right-sided (Ramsdell et al., 2006), plus the inflow and outflow poles are brought closer together to produce the classic heart-shape common to terrestrial animals. Inside the ventricle, ridges of trabeculae muscle form on the lumenal surface of anterior and posterior walls. Trabecular cardiomyocytes orient their myofibrils along the ventricle length (parallel to axes). The result is a layered chamber wall structure with a ‘cross-grained’ configuration of myofibrils.

Myofibrils initially form near the surface of muscle cells and hence, rearrangement of the actin cytoskeleton in the cell cortex is a crucial early step. Whether these actin stress fibres act as temporary templates, or actually physically transition, during the production of muscle-type α-actin filaments with their uniform (sub-sarcomeric) length and opposed polarity remains an active topic of research (Fenix et al., 2018 and refs therein). The starting points for assembly are aggregations of α-actin and α-actinin 2 (termed Z-bodies) that associate with the cell membrane at integrin adhesion sites (proto-costameres) (Reviewed Sparrow and Schöck, 2009). For recruitment of the motor proteins, intermediate steps involve incorporation of non-muscle myosin type II into filaments before it is then replaced by muscle myosin II protein to establish the correct sarcomere spacing (Du et al., 2008; Fenix et al., 2018). There is a mutual dependency between proper formation of actin and myosin filaments in muscle, although recent experiments have observed part assembled components of each in the absence of the other structure (Fenix et al., 2018; Rui et al., 2010). Chaperones and co-chaperones such as Unc45b that facilitate folding of the myofilament proteins have been identified (Reviewed Carlisle et al., 2017) and the number will likely increase, given the size and complexity of sarcomere architecture (Reviewed Gautel and Djinović-Carugo, 2016). Models that describe these essential steps during myofibrillogenesis don’t, however, address the timing and spatial requirements that are fundamental to cardiac chamber growth in the embryo. What determines the precise locations within the cardiomyocyte that initiate filament precursor association, when is this triggered in each cell and in which direction does myofibril assembly then proceed? It should be emphasized that the ‘where, when and in which direction’ questions surrounding cardiac myofibrils are equally pertinent to the prospects for stimulating repair of damaged adult (human) hearts as they are for studying normal embryonic chamber development.

One gene that exerts a profound effect on myofibril formation during the early stages of heart chamber growth is *adprhl1*, which encodes the protein ADP-ribosylhydrolase-like 1. By sequence similarity, Adprhl1 belongs to a small group of enzymes found in vertebrates that catalyse hydrolysis cleavage reactions involving ADP-ribosylated substrates (Mashimo et al., 2014; Moss et al., 1992). Nevertheless, the familial active site differs in Adprhl1 and lacks the necessary amino acids required to support catalytic activity (see Discussion) (Oka et al., 2006; Rack et al., 2018). Pseudoenzymes such as this can be a challenge to study, but there is gathering evidence that Adprhl1 may be an important factor in cardiogenesis.

In a series of experiments using *Xenopus* embryos, we first showed that heart-specific expression of *adprhl1* mRNA was biased towards actively growing, chamber myocardium (Smith et al., 2016). Expression is conserved in mouse embryo hearts and *ADPRHL1* is also among the first mRNAs induced as human embryonic stem cells are differentiated *in vitro* towards a cardiac fate (Beqqali et al., 2006). When *adprhl1* activity was knocked down in *Xenopus* using antisense morpholino oligonucleotides (MO) that inhibit RNA-splicing, a consistent, severe heart defect occurred (Smith et al., 2016). Hearts formed with small ventricles that could not beat. The number of cardiomyocytes present as ventricle formation commenced was not affected by the MO, nor was propagation of the electrical calcium signal across the inert ventricle and the expression of myofibrillar subunit genes was unaltered. Nonetheless, a sequence of myofibrillogenesis defects was observed, beginning with delayed cortical actin rearrangement and loss of cell elongation at the presumptive ventricle apex. The few myofibrils that did eventually form in the ventricle remained short and disarrayed, with no consistent trajectory of growth, and the overall result was a wholesale failure to the cellular architecture of the chamber.

In normal tadpoles, our Adprhl1-specific antibody identified two cardiac proteins of 40 and 23 kDa size. The larger protein matched the expected translation product from *adprhl1* exons 1 to 7, but the precise composition of the smaller species was unknown, despite its abundance increasing as chamber development proceeded. Significantly, both proteins were lost from heart extracts of MO injected embryos. Stimulating 40 kDa Adprhl1 production using cardiac-specific transgenes was potent and triggered severe myofibril structural abnormalities as if excessive formation of Z-disc precursors had occurred (Smith et al., 2016). Recombinant 40 kDa Adprhl1 that included an N-terminal epitope tag additionally showed a direct association with myofibrils that the antibody could not detect. In some linear myofibrils, this Adprhl1 localized to two clear stripes on either side of the Z-disc and also a diffuse stripe at the H-zone of the sarcomere, potentially marking the boundaries of actin filaments. Overall, the *Xenopus* experiments describe a gene that is fundamentally required for proper cardiac myofibrillogenesis and ventricle growth, possibly through a physical interaction of Adprhl1 with nascent actin filaments. Nonetheless, the picture is complicated by the presence of two distinct sized Adprhl1 proteins in the heart, with the variant that most resembles the ADP-ribosyl-acceptor hydrolase family limited by strict constraints on its synthesis and exerting an adverse effect on Z-disc structure (see Results).

In four subsequent years, there has been just a single report that linked a missense sequence variation in the human *ADPRHL1* locus to a specific clinical defect of the left ventricle through a genome-wide association study (GWAS) (Norland et al., 2019). No other gene knockdown or knockout in other experimental model animal species has yet validated *adprhl1* function during heart development. Because our understanding of *adprhl1* is over-reliant on the results of MO experiments, we have turned to CRISPR/Cas9 technology in order to induce mutations across the *Xenopus adprhl1* gene. Co-injection of synthetic guide-RNA (gRNA) and Cas9 endonuclease into one-cell stage *X. laevis* embryos cuts double-strand DNA breaks (DSB) at the targeted sequence with 100% efficiency. It is the nature of the attempted repairs made to the DSB that determine whether alleles of the gene retain function or are inactivated. In G0-generation animals, there is also mosaic cellular distribution of the resulting alleles. Hence, a search for defective embryonic phenotypes by testing different gRNAs distributed throughout *adprhl1* exons is actually a screen for essential positions where sequence variations including in-frame mutations are not tolerated.

We identify a sequence within exon 6 of *adprhl1* where targeted mutation causes small in-frame deletions and produces tadpoles with dysfunctional cardiac ventricles. At the level of myofibrillogenesis, the malformations are remarkably consistent with previous MO work. Crucially, this region of exon 6 encodes a di-arginine motif unique to Adprhl1 that is situated within the ancestral active site cleft of the ADP-ribosylhydrolase family. We discuss the relationship between Adprhl1, ADP-ribosylation pathways and the further link to actin polymerization in order to consider the role of Adprhl1 in the heart. Finally, in mice carrying a partial deletion of the *Adprhl1* gene, we examine cardiac ADPRHL1 protein production to show that for mammals at least, selective loss of the 40 kDa form has no adverse effect on cardiogenesis. Results from each species complement each other and point to where critical residues for Adprhl1 action are located.

## Materials and Methods

### Tyrosinase gRNA sequences

Two gRNAs were previously designed against both *X. laevis tyrosinase* homeologous alleles (Wang et al., 2015). Their gene-specific sequences correspond to the *tyr* DNA listed below.

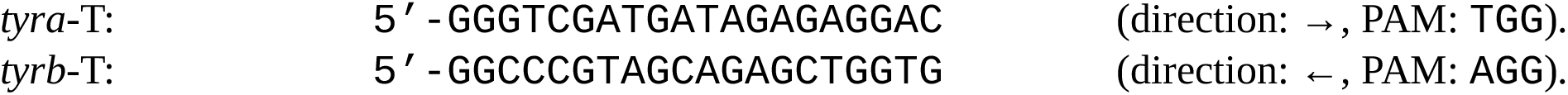

### Adprhl1 gRNA sequences

The *adprhl1* sequences used to design the principal gRNAs used in the study are listed below. Mismatched bases of control gRNAs are coloured red.

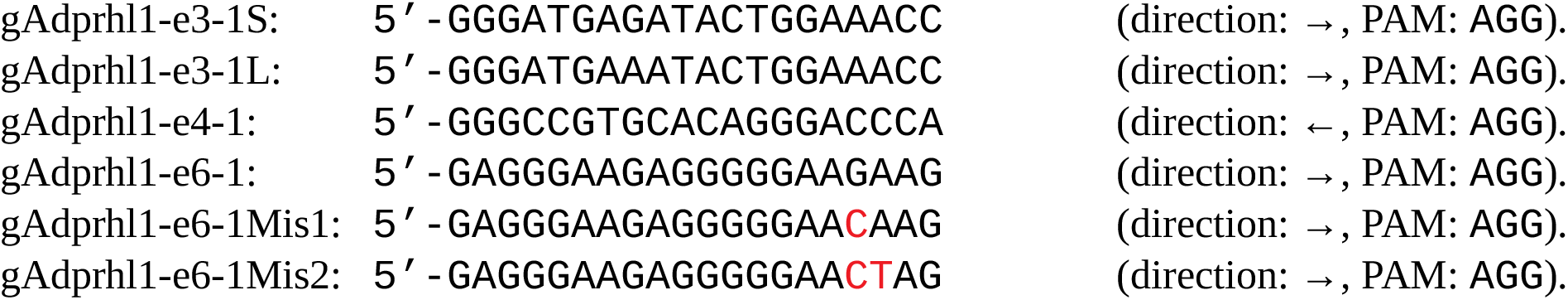

### Additional adprhl1

gRNAs were also tested for activity, some with longer 21-22 nucleotides of gene-specific sequence. Mismatched 5’-bases added to enable efficient T7-transcription are coloured blue.

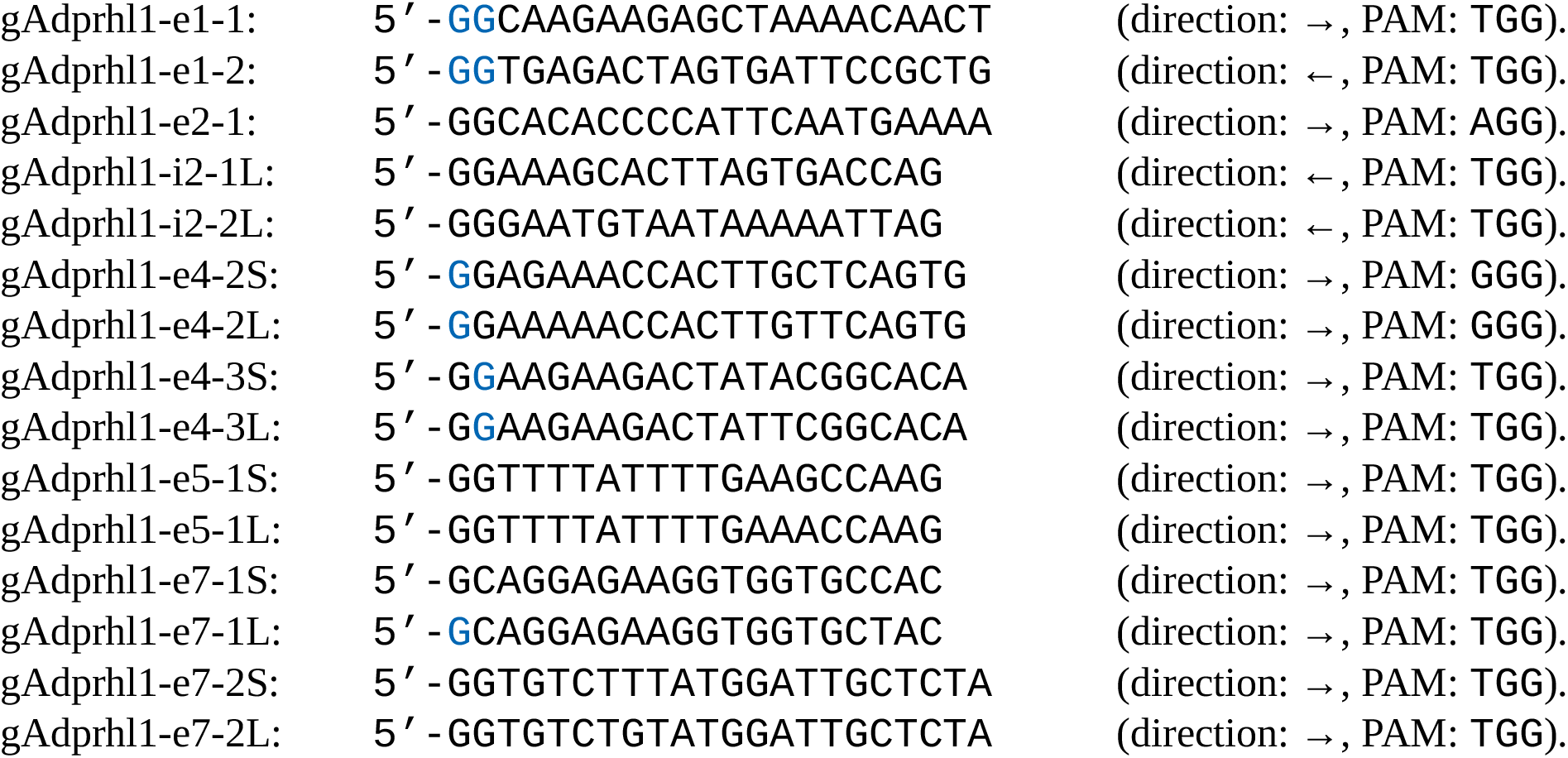

Note -e5-1 (S and L) fits the optimal gRNA consensus but gave consistently poor synthesis yields due to an adverse T-repeat sequence and could not be assessed by injection into embryos.

Current computational tools used to predict off-target activity of gRNAs have difficulty advancing beyond sequence alignment complementarity towards identifying those potential off-targets that are likely to be cleaved *in vivo* (Reviewed Wilson et al., 2018). Here, all gRNA sequences were assessed for hybridization at other positions using the JGI *X. laevis* genome v9.1 and Cas-Offinder (Bae et al., 2014), which considered the potential for both sequence mismatches and bulges. Results for potential off-targeting by the most active gAdprhl1-e6-1 gRNA to other gene coding sequences are listed in the Supplementary Methods. None of the additional genes would be anticipated to contribute to a heart phenotype if mutated and none of the potential interactions align with both of the homeologous alleles present in *X. laevis*.

### Cas9 and gRNA injection into Xenopus laevis embryos

Preliminary experiments tested the activity of different Cas9 RNAs by co-injecting them with the two gRNAs that target *tyrosinase*. A mixture containing a Cas9 RNA (125 pg/nl) and the two gRNAs (both at 125 pg/nl) was prepared and 4 nl injected into one-cell stage embryos, giving a final 500 pg mass of each reagent. Injections were directed towards the animal pole of the embryo (uppermost third). Injection of embryos continued from 35 until 60 minutes post-fertilization. They were incubated at room temperature (22 °C) until 90 mpf, then transferred to 17 °C. Culture media used for injection and first 24 hours incubation was 0.1xNAM, 0.5 % Ficoll^®^-400, 20 µg/ml gentamycin. Thereafter, 0.1xNAM was used.

Greater efficiency of *tyrosinase* and *adprhl1* gene knockout was achieved using a commercial Cas9 protein preparation, EnGen^®^ Spy Cas9 NLS (NEB: M0646M). In this case, gRNAs were preloaded onto Cas9 protein using a mixture assembled in the following order: 1 µl 1.3 M KCl (302 mM final), 2 µl gRNA (<1 µg of a single gRNA, so <233 pg/nl final) and 1.3 µl EnGen Cas9 (26 pmol). The mixture was incubated at 37 °C for 10 minutes, immediately prior to injection. Injection of 4 nl mixture and subsequent embryo culture was as before. The method was adapted from Burger *et al* (2016). For *tyrosinase* knockout, the phenotype classes used to define the extent of pigmentation-loss are described in Supplementary Figure S6 and follow those of Guo *et al* (2014).

For *adprhl1* knockout, embryos that gastrulated normally were allowed to develop to tadpole stage 44. Their external morphology was recorded each day and the appearance of their heart closely monitored. Tadpoles were assigned to one of four distinct phenotype classes, which differed slightly from those used to assess morpholino injection (see Supplementary Methods):

Heart defect 1 - inert ventricle. As per MO study.

Heart defect 2 - thin wall ventricle. In tadpoles showing a cardiac oedema that produced a beating heart, the ventricle was frequently thin-walled and became increasingly dilated by stage 44. This was especially true for embryos that received the exon 6 -e6-1 gRNA. Other malformations. Any non-cardiac developmental defect visible externally by stage 44, however subtle.

Normal (heart) morphology. Perfect development through to stage 44.

### Sanger sequence analysis of mosaic adprhl1 exon 3, 4 and 6 mutations in G0-generation X. laevis

DNA was extracted from individual tadpoles. Frozen tissue was mixed with 200 µl 50 mM Tris pH8.8, 1 mM EDTA, 0.5% Tween20 containing freshly added 600 µg/ml proteinase K and incubated at 55 °C for 20 hours. PCR amplification of *adprhl1* genomic sequences used Platinum SuperFi DNA polymerase (Invitrogen) and 1 µl tadpole extract per 30 µl reaction. Plasmid clone isolates of amplicon DNA were prepared using a Zero Blunt™ TOPO™ PCR cloning kit (K287520, Invitrogen) and their inserts were sequenced. Sanger sequencing allowed study of 2 kbp amplicons so that the presence of larger deletions and insertions could be detected. The PCR primers used for genotyping are listed in Supplementary Methods.

Each mutated sequence was assigned a genotype score (a number code) according to the size of the amino acid lesion that it encoded. This classification of mosaic mutations found within an individual tadpole was then compared against its cardiac morphology. Some assumptions were made for sequence deletions that disrupted exon splice junctions. Where a splice acceptor site was lost, it was assumed the exon was skipped. Where a splice donor sequence was removed, it was assumed the following intron was inappropriately retained.

Genotype score codes.

01: Inactive mutant. Frame-shift-stop or nonsense mutation.

02: In-frame mutant causing more than 20 amino acid changes.

03: In-frame mutant causing between 11-20 amino acid changes.

04: In-frame mutant causing 6-10 amino acid changes.

05: In-frame mutant causing 1-5 amino acid changes.

06: Normal amino acid sequence.

### Amplicon-EZ (NGS) sequence analysis of mosaic adprhl1 exon 6 mutations

The Amplicon-EZ next generation sequencing service (Genewiz) was used as a cost-effective method to obtain deeper coverage of the mosaic exon 6 mutations from individual tadpoles, providing 50,000 reads per sample. Genomic PCRs obtained with primers p2560+p2648 were sequenced directly using Illumina^®^ technology. The Galaxy web platform (www.usegalaxy.org) was used to assemble sequence reads and combine duplicates (coalesce identical aligned reads) (Afgan et al., 2018). Reads were aligned to small 100 bp reference sequences of the *adprhl1* S- and L-alleles, covering 50 bp either side of the -e6-1 gRNA PAM. SeqMan Pro (DNASTAR) was also used to sort the sequences according to the variants found at the gRNA site and count the number of wild-type reads that occurred for each tadpole.

### Western blot detection of Adprhl1 protein

Adprhl1 protein was detected using a rabbit antibody raised against an ADPRHL1 peptide (mouse ^248^DNYDAEERDKTYKKWSSE^265^, encoded by exons 5-6) (Smith et al., 2016). The antibody is active against the *Xenopus*, mouse and human species orthologs. For protein extraction from embryonic *Xenopus* hearts, typically 100 hearts were dissected, pooled, snap frozen, homogenized in 120 µl RIPA buffer and boiled with an equal volume of 2x reducing protein sample buffer. Individual adult mouse hearts were homogenized in 400 µl RIPA buffer, the resulting slurry mixed with 400 µl sample buffer, then aliquots diluted a further three-fold with RIPA/sample buffer before SDS-PAGE.

### Immunocytochemistry of Xenopus hearts

Immunocytochemistry was performed on whole tadpoles and subsequently the hearts were dissected, mounted in 12 µl CyGEL Sustain (biostatus) and viewed using Zeiss LSM5-Pascal or LSM710 confocal microscopes. Confocal images of whole hearts captured 2 µm deep optical sections. Both the ventricle myocardial wall and also deeper trabecular layers were assessed by scanning different depths. Images of cardiomyocytes and myofibrils were 1 µm optical sections, at a depth 1-2 µm below the outer (apical) myocardial surface. Antibodies used were Adprhl1, Myosin A4.1025 (DSHB) and phospho-Histone H3(Ser10) 3H10 (Sigma), along with fluorescent dye-conjugated secondary antibodies (Jackson ImmunoResearch). Atto 633-conjugated phalloidin (Sigma) stained actin filaments. Dying cells were visualized with the ApopTag^®^ Red In Situ Apoptosis Detection kit (Sigma).

## Results

### Current understanding of Adprhl1 function relies on RNA-splice interfering morpholinos

Our previous experiments explored the role of Adprhl1 during heart chamber formation in *Xenopus* embryos. Expression of *adprhl1* mRNA is restricted to the heart myocardium and select ocular muscles, in contrast to the multiple tissues that stain for the founding member of the ADP-ribosylhydrolase family, *adprh* (Fig 1A, B). We knocked down *adprhl1* activity using morpholino oligonucleotides that target RNA-splicing. The Adprhl1-e2i2MO caused retention of intron 2 while -i2e3MO induced exon 3 skipping (Smith et al., 2016). For both MOs, this led to a wholesale loss of *adprhl1* mRNA abundance as measured by wholemount *in situ* hybridization (Fig 1D, F). Our Adprhl1-specific antibody identified two cardiac proteins of 40 and 23 kDa size and both were lost from heart extracts of MO injected embryos (Smith et al., 2016). Thus RNA-splice interfering MOs have defined activities and cause a well-documented defect in embryonic cardiogenesis by stage 40. The ventricle remains small and inert (Fig 1C, E), with oedema developing beyond stage 41. In reviewing earlier work, it should also be noted that the antibody, when used for immunocytochemistry, could not detect Adprhl1 protein in the heart above background levels of signal. This reflected the low endogenous production of cardiac Adprhl1 that occurs *in situ*. Vertebrate gene models of *adprhl1* now also include an alternative transcript containing an additional 3’-exon 8. This long exon was not expressed in the forming heart, although suitably large 100 kDa Adprhl1 protein species have been detected in tadpole gut tissue (data not shown).

**Fig 1.**
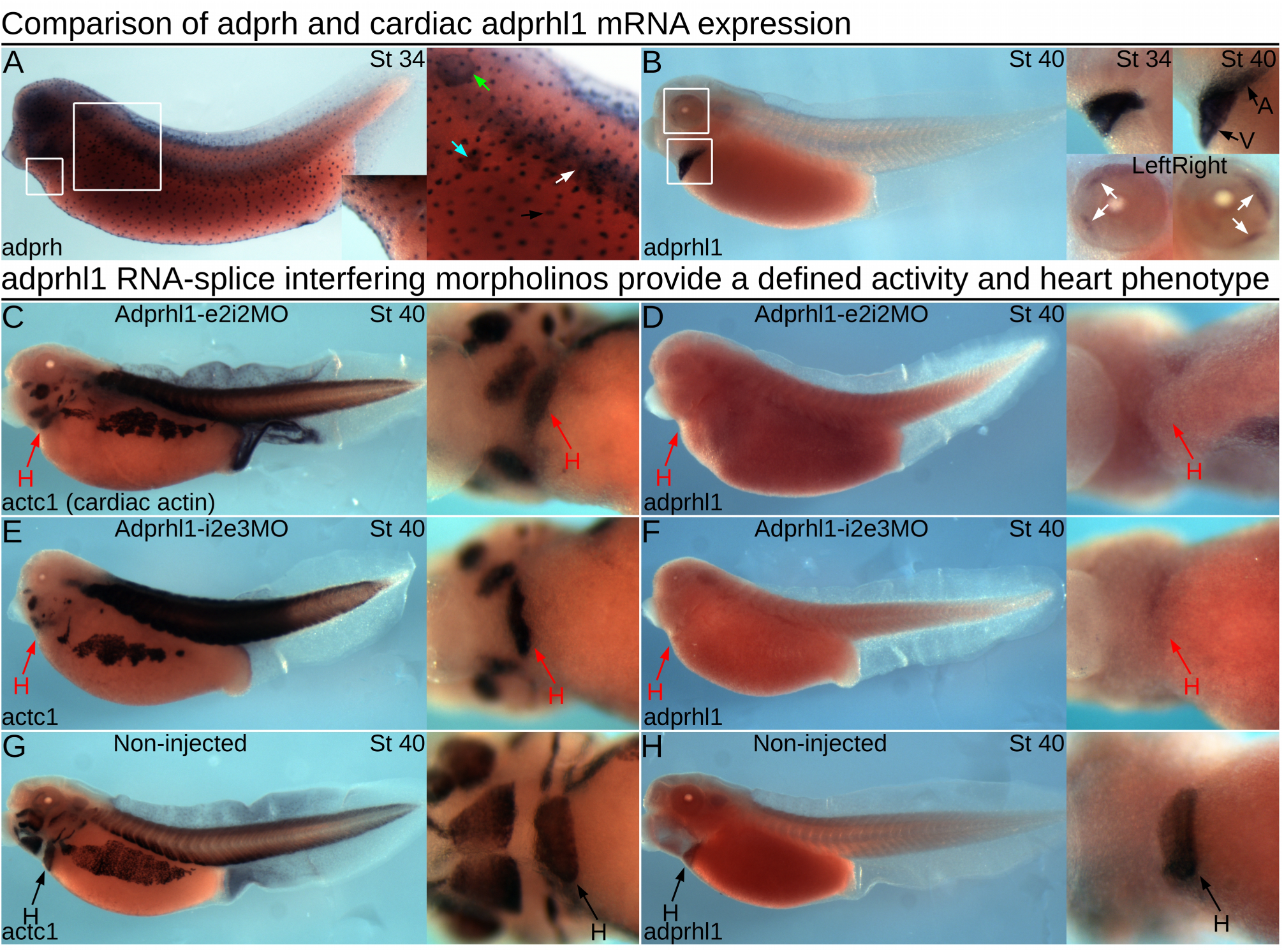
Cardiac adprhl1 expression and morpholino knockdown in Xenopus embryos. **A, B:** Comparison of *adprh* and *adprhl1* mRNA expression. A stage 34 tadpole (left-lateral view, A) shows principal sites of *adprh* expression, with the position of detail images marked by white squares. Mucus producing small secretory epidermal cells (black arrow) contain *adprh* from stage 28 (see Dubaissi et al., 2014), detection in somites (white arrow) resolves towards hypaxial (ventral) muscle groups by stage 38, transient expression occurs in nephrostomes of the pronephros (cyan arrow), plus otic vesicle (green arrow), pharyngeal arches and the brain. A stage 40 tadpole (B, plus a stage 34 detail image) shows strong *adprhl1* mRNA expression in the heart myocardium and also in the eyes within two forming muscle blocks, located medially (anterior) and at superior (upper) and inferior (lower) positions (white arrows). **C-H:** *adprhl1* RNA-splice interfering MOs provide a defined activity and inert heart phenotype. **C, D:** Expression of *actc1* (heart and skeletal muscle, C) and *adprhl1* (D) mRNAs in stage 40 tadpoles after injection of 32 ng Adprhl1-e2i2MO at the one-cell stage. Impaired heart ventricle growth and a loss of *adprhl1* mRNA signal is observed. Left-lateral view of tadpole and detail ventral view of heart region presented. **E, F:** Identical heart phenotype caused by injection of the distinct Adprhl1-i2e3MO morpholino. **G, H:** Normal ventricle size and *adprhl1* signal in non-injected sibling tadpoles. Red arrows denote aberrant morphology. H, heart; A, atrium; V, ventricle.

### Combining morpholino knockdown with transgenic 40 kDa over-expression experiments reveals a complexity to Adprhl1 action

Using a cardiac-Gal4/UAS system, we also produced a series of transgenes to drive over-expression from *adprhl1* cDNAs. They actually revealed strict translational control operating in the heart that must act to restrict the synthesis of the endogenous 40 kDa *Xenopus* Adprhl1 protein (Smith et al., 2016). In fact, the only way to achieve additional Adprhl1 production was to engineer changes to the transgene 5’-cDNA sequence adjacent to the translation initiating ATG. One transgene exchanged the 5’-most 156 bp for the corresponding coding sequence from human-species *ADPRHL1* cDNA (*Tg[UAS:human*^*1-52*^*-Xenopus*^*53-354*^ *adprhl1])*. It escaped the regulation and encodes a human-*Xenopus* hybrid Adprhl1 containing 21 amino acid substitutions compared to the native *Xenopus* protein. A second transgene incorporated silent nucleotide changes (synonymous substitutions) within the *Xenopus* coding sequence (*Tg[UAS:Xenopus adprhl1(silent 1-282bp)])*. This only partially evaded the endogenous control such that recombinant Adprhl1 accumulated transiently in a fraction of the cardiomyocytes. Nonetheless, it does offer the advantage that its translated protein remains identical to natural *Xenopus* Adprhl1.

The activity of both transgenes was unaffected by the RNA-splice interfering MOs. We examined whether recombinant Adprhl1 proteins could rescue the MO defects in tadpoles, in particular the assembly of myofibrils and morphology of the resulting ventricle chamber. These experiments are presented in Supplementary Figure S1, with a concise version showing the key panels reproduced as Figure 2. Only a limited recovery of cardiac myofibril assembly is possible when using recombinant 40 kDa Adprhl1 protein.

**Fig 2.**
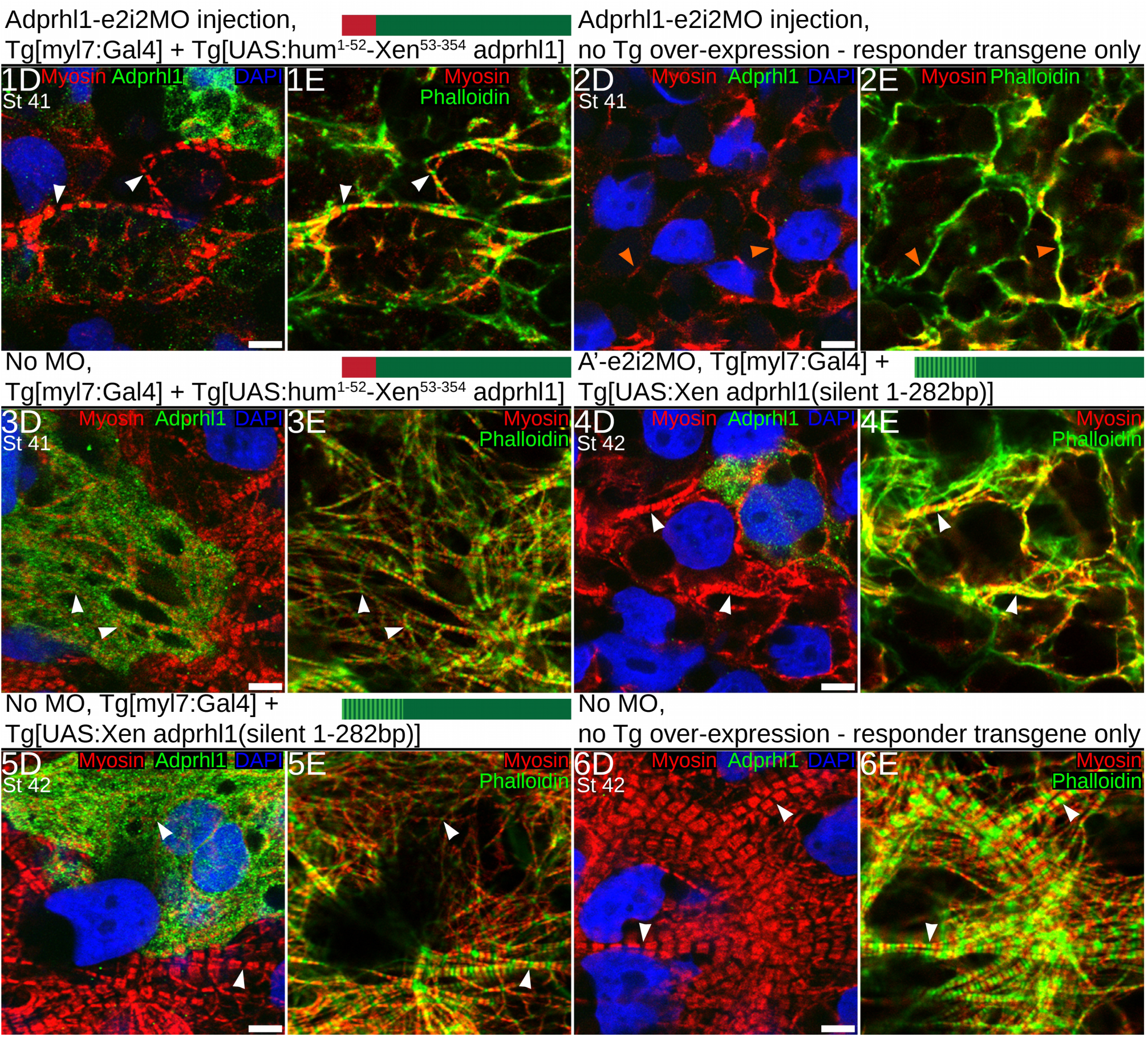
Limited recovery of cardiac myofibril assembly in adprhl1 morpholino injected embryos by transgenic synthesis of recombinant 40 kDa Adprhl1 proteins. Experiments that combine *adprhl1* MO knockdown with two distinct transgenes engineered to achieve *adprhl1* over-expression. This is the concise version of Supplementary S1. For brevity, Fig 2 presents only the high magnification (D and E) images that reveal ventricle wall myofibril patterns within the experimental hearts. The extended figure additionally includes the morphology of each ventricle, the extent of Adprhl1 protein production within, plus squares to locate the position of the myofibril images within each ventricle. **1D, E:** Cardiomyocytes within the heart ventricle wall (anterior surface) of a stage 41 tadpole that was injected with the RNA-splice interfering MO, Adprhl1-e2i2MO, into dorsal (D-2/4) blastomeres. Additionally, it carried binary transgenes to over-express recombinant Adprhl1 protein, consisting of *Tg[myl7:Gal4]* driver and the *Tg[UAS:human*^*1-52*^*-Xenopus*^*53-354*^ *adprhl1]* responder. Scale bar = 5 µm (for D, and E). Fluorescence image (D) shows anti-Adprhl1 immunocytochemistry (green), anti-myosin (red) and DAPI-stained nuclei (blue). The second panel (E) displays a merge of myosin and phalloidin actin stain, with the phalloidin coloured green to evaluate signal overlap. **2D, E:** Ventricle cardiomyocytes from a sibling tadpole that received the same Adprhl1-e2i2MO injection but carried only the UAS-responder transgene and hence did not produce excess recombinant Adprhl1. **3D, E:** A double transgenic sibling that synthesized recombinant human-*Xenopus* hybrid Adprhl1 but was not injected with the MO. **4D, E:** Ventricle cardiomyocytes from a second experiment, a stage 42 tadpole that was injected with Adprhl1-e2i2MO and carried the *Tg[myl7:Gal4]* driver but a different *Tg[UAS:Xenopus adprhl1(silent 1-282bp)]* responder transgene. This incorporates silent nucleotide changes (synonymous substitutions) to the cDNA sequence in order to partially evade endogenous translational regulation. **5D, E:** Ventricle cardiomyocytes of a double transgenic, silent mutation, sibling tadpole that synthesized recombinant *Xenopus* Adprhl1 but was not injected with the MO. **6D, E:** A non-injected sibling control harbouring only the silent mutation responder transgene that did not produce excess recombinant Adprhl1. Paired white arrowheads indicate Z-disc sarcomere positions, orange arrowheads denote non-striated filaments. V, ventricle; OT, outflow tract.

After injection of Adprhl1-e2i2MO into dorsal blastomeres, the resulting small ventricle revealed severely disrupted myofibrillogenesis, with actin fixed at the cell cortex and no sarcomere striations evident in either myosin or actin filaments (Supplementary S1#2A-E, or Fig 2#2D, E, orange arrowheads). Separately, in the heart expressing the human-*Xenopus* hybrid, clusters of cardiomyocytes with strong Adprhl1 signals occurred adjacent to regions with more modest production (Supplementary S1#3A-E, or Fig 2#3D, E). Fewer bright cells contributed to the ventricle apex region (Supplementary S1#3B). Over-expression of hybrid Adprhl1 altered myofibril patterns even in the absence of the MO. There was greater disarray to the myofibril orientation and thinner actin filaments were observed (Fig 2#3D, E, arrowheads). By combining the two interventions, the uncontrolled synthesis of 40 kDa hybrid Adprhl1 in hearts where the MO had removed endogenous Adprhl1 production did not rescue normal cardiac development and the small hearts remained inert (Supplementary S1#1A-E, Fig 2#1D, E). Indeed, the first signs of oedema could be detected at stage 41 in tadpoles that received the MO (Supplementary S1#1A, 2A). Nonetheless, myofibril assembly did actually occur within cells where lower levels of recombinant Adprhl1 were detected (Fig 2#1D, E). Myofibrils extended in the perpendicular direction that is necessary to support proper ventricle chamber growth (Smith et al., 2016). Sarcomeres were evident in both myosin and actin filaments suggesting a functional maturity of the myofibrils (Fig 2#1D, E, arrowheads).

With the transgene containing silent mutations, scarce ventricular cells containing excess 40 kDa Adprhl1 had profoundly affected myofibrillogenesis (Supplementary S1#5A-E, Fig 2#5D, E). Their dense network of thin, striated myofibrils linked together by branch points at every Z-disc suggested excessive formation of Z-disc precursor structures (Fig 2#5D, E). They contrasted with the mature myofibrils found within adjacent control cardiomyocytes (Fig 2#5D, E, also Fig 2#6D, E). Once again, a functional rescue of the MO phenotype was not achieved by this transgene as it yielded too few cells synthesizing recombinant *Xenopus* Adprhl1 (Supplementary S1#4A-E, Fig 2#4D, E). Yet despite the widespread disarray in the heart presented, sarcomeres in myofibrils formed emanating from (or adjacent to) the sole ventricular cardiomyocyte containing detectable Adprhl1 protein (Fig 2#4D, E, arrowheads).

In summary, loss of Adprhl1 halts cardiogenesis while 40 kDa over-production is potent and triggers severe myofibril structural abnormalities. A full rescue of the MO defect is not achieved with the 40 kDa protein, although cortical actin turnover and some myofibril assembly can be restored. The natural control of Adprhl1 synthesis is far more sophisticated than current transgene technology. Thus, a clean rescue would require further understanding of Adprhl1 primary sequence and function, the relative contributions of both the 40 kDa and 23 kDa forms of Adprhl1, plus the mechanism for translational regulation to be determined. Nonetheless it is clear that altering Adprhl1 levels has serious consequences for myofibrillogenesis during heart formation.

Other features of the *adprhl1* transgenes are summarized in Supplementary S2 and S3. The cardiomyocytes producing recombinant *Xenopus* Adprhl1 were viable cells. They were not unusually mitotic and nor were they dying (Supplementary S3). Moreover, for the hybrid Adprhl1 at least, translation of 40 kDa Adprhl1 did not lead to a consequent increase in 23 kDa Adprhl1 abundance, suggesting the latter is not a processed fragment of the former (Supplementary S2).

**Fig 3.**
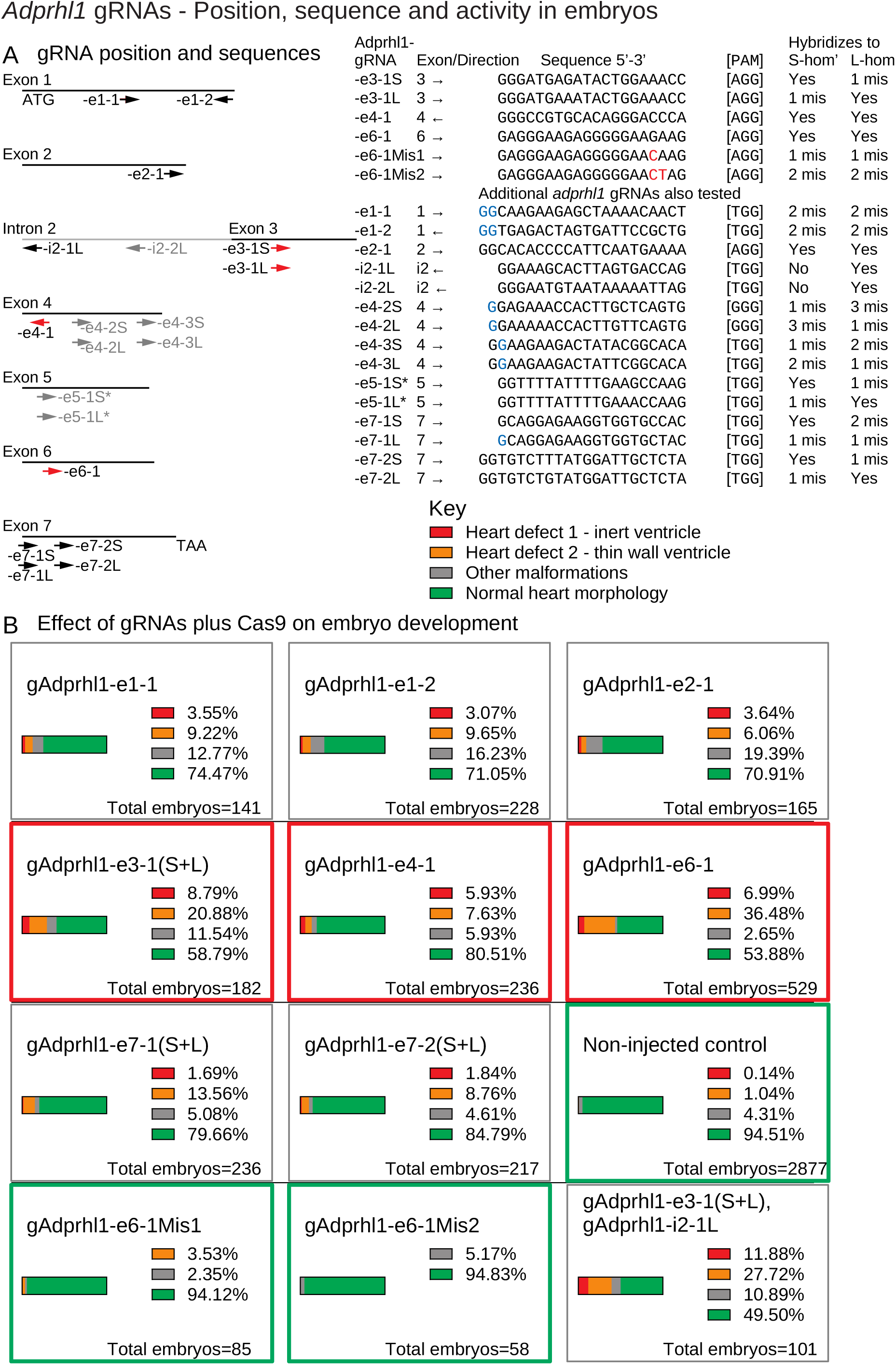
Adprhl1 gRNAs - Position, sequence and activity in embryos. **A:** Diagram showing the hybridization position of gRNAs in relation to exons of the *X. laevis adprhl1* locus. Separate gRNAs for S- and L-homeologous alleles were prepared if sequence differences existed between the two at the selected location. Arrows indicate the 5’-3’ direction of each gRNA. Red arrows denote gRNAs whose activities were further examined by sequencing mutant allele DNA. Black arrow gRNAs have their effects on embryo development presented in this figure while grey gRNAs feature in Supplementary S7. The table lists the *adprhl1*-specific sequence of each gRNA along with its protospacer-associated motif (PAM). Mismatched bases of control gRNAs are coloured red. Mismatched 5’-bases added to enable gRNA transcription from plasmid template DNA are coloured blue. Note -e5-1 (S and L) targeting exon 5 gave poor synthesis yields (Materials and Methods). **B:** Effect of *adprhl1* gRNAs plus Cas9 on embryo development. Parts-of-whole charts showing the frequency of stage 44 tadpole phenotypes that occurred after injection of gRNA along with Cas9 protein into one-cell stage embryos. Red rectangles surround charts for the principal gRNAs whose activities were also examined by DNA sequencing. Green rectangles denote control gRNAs and also a chart presenting the cumulative total for all non-injected sibling tadpoles from the experiments. The lower right chart shows the consequence of injecting a mixture of gRNAs to target adjacent intron 2 - exon 3 regions of the *adprhl1* gene (additional combinatorial gRNA experiments are shown in Supplementary S7). Heart defects were detected at higher frequency using the gAdprhl1-e3-1 and in particular the -e6-1 gRNA. Images showing representative tadpoles after gAdprhl1-e6-1 mutation are shown in Fig 6.

### Limitations of adprhl1 morpholinos - Contrast between RNA-splicing versus translation inhibition

One way to determine the role of the 40 kDa Adprhl1 would be to utilize MOs that specifically inhibit translation initiation of this protein form. As a pseudo-tetraploid species, *Xenopus laevis* has separate loci for *adprhl1* arranged on distinct S- and L-homeologous copies of chromosome 2 (Karimi et al., 2018; Session et al., 2016). However, each of three overlapping MOs covering both S- and L-allele sequences surrounding the 5’-most AUG of *adprhl1* mRNA caused unforeseen defective tail growth observed from stage 32 (Supplementary S4 and S5F, G). Cardiac ventricle growth was clearly impaired in these tadpoles (Supplementary S5F, G) and heart tissue showed loss of 40 kDa Adprhl1 (data not shown), but this was overshadowed by the other prominent malformations. The potential of using MOs to identify a translation initiation site for the 23 kDa Adprhl1 protein was also explored, but did not provide convincing answers (see Supplementary S4 and S5H-O). It is not clear whether the translation inhibition MOs simply have poor specificity for *adprhl1* or if additional early effects on maternally-derived transcripts occur. For now, the previously published RNA-splice interfering MOs remain the only antisense reagents that achieve a clean cardiac *adprhl1* knockdown with an efficiency close to a null phenotype.

**Fig 4.**
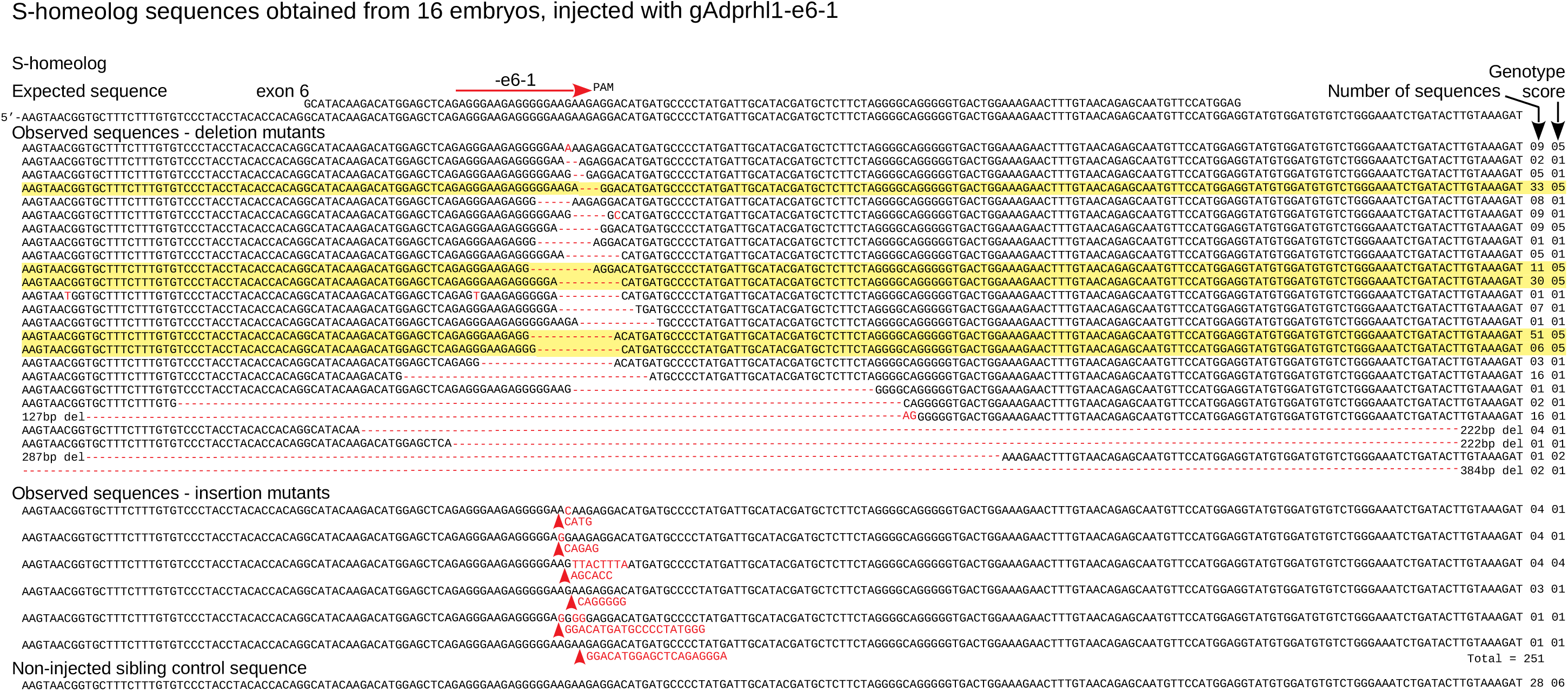
Targeting adprhl1 exon 6 causes near complete mutation and in-frame repair bias - S- homeolog DNA sequences. Sanger cloned DNA sequences of *adprhl1* S-homeologous locus exon 6 after mutation by the gAdprhl1-e6-1 gRNA plus Cas9. Mutated sequences from the L-locus exon 6 are presented in Supplementary S13. The gRNA position is shown by the red arrow placed above the expected sequence (top 2 rows, exon and genomic). Alignment of 251 (S-) DNA clones obtained from 16 tadpoles, with every sequence carrying a lesion at the gRNA binding site. Mutant nucleotide sequences are coloured red. A missense mutation is listed first, followed by deletions (red hyphens) and then insertions (red arrowheads). Frequently occurring sequences containing in-frame 3, 9 or 12 bp deletions are highlighted. The number of instances of each sequence is to the right, alongside its genotype score.

**Fig 5.**
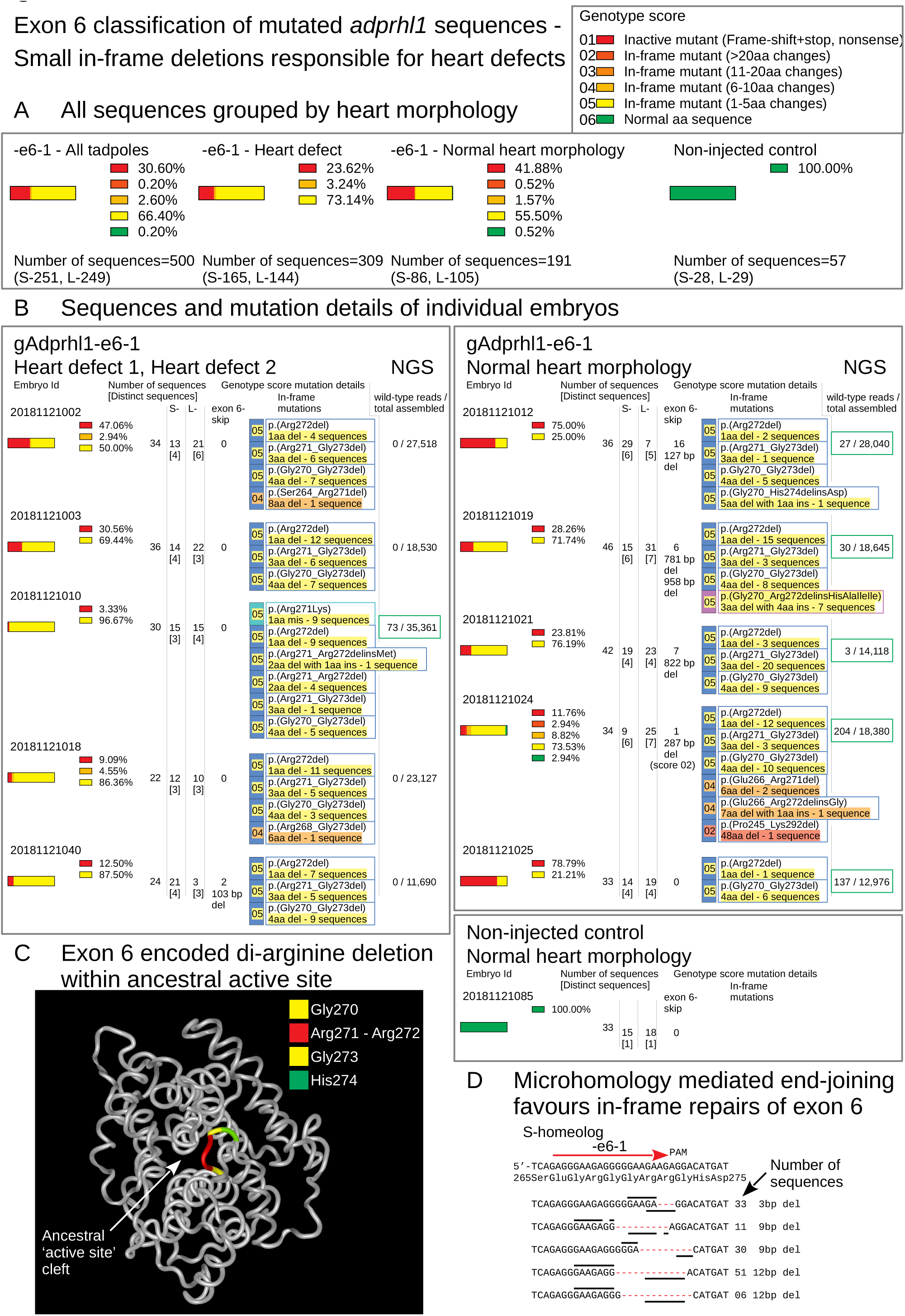
Exon 6 classification of mutated adprhl1 sequences - Small in-frame deletions responsible for heart defects. **A:** All cloned Sanger sequences grouped by heart morphology. Parts-of-whole charts record the frequency of exon 6 sequence genotype scores tallied for all embryos that received Cas9 and the gAdprhl1-e6-1 gRNA (far-left chart), or divided into two groups representing tadpoles with heart defects (centre-left chart) versus those whose hearts developed normally (centre-right chart). Sequences from non-injected control embryos were also compared (far-right chart). The total number of S- and L-locus sequences analysed is listed below each chart. The key to interpret the genotype score is also included. There was just one wild-type sequence detected among 500 examined and for this region of exon 6, there was a disproportionately high frequency of in-frame deletion mutations. **B:** Sequences and mutation details of individual embryos. Separate charts for 11 of the embryos, including 5 tadpoles with heart defects, 5 with normal heart morphology and one non-injected sibling. Columns list the number of Sanger sequences and the number of distinct sequences (in square brackets) for each embryo, plus the presence of larger deletions (and their size) that would skip exon 6. Most importantly, the precise amino acid changes are given for all of the in-frame lesions. A standard nomenclature for protein sequence variations is used to describe the changes, with a simplified summary (highlighted) underneath stating whether the mutation caused a missense, deletion or aa insertion modification. Highlight colour matches the genotype score of the sequence. The rectangle border colour denotes a missense (teal), net deletion (blue) or net insertion (violet). Three specific amino acid deletions were common to many of the embryos: a single aa loss of Arg272, loss of 3 aa Arg271 to Gly273, or loss of 4 aa Gly270 to Gly273. Deletion of these amino acids must eliminate Adprhl1 protein function and thus be responsible for the cardiac malformations caused by the -e6-1 gRNA. The far-right column shows next generation sequence data (NGS, Materials and Methods) employed specifically to search for the presence of wild-type reads at the gRNA site. For the majority of injected embryos with a heart defect, increasing the depth of mutation analysis did not reveal any wild-type alleles. In contrast, wild-type alleles were found in each of the tadpoles with a normal heart. **C:** Structure model of *X. laevis* Adprhl1 protein backbone, oriented with the ancestral active site foremost (Smith et al., 2016). The di-arginine (Arg271-Arg272 coloured red) sequence frequently deleted by the -e6-1 gRNA resides in a loop at the heart of the active site (Gly270, Gly273 yellow, His274 green). **D:** S-locus sequence at the -e6-1 gRNA site. Five rows represent common in-frame deletions of 3, 9, 9, 12 and 12 bp. Lines above and below the rows mark short direct repeats of nucleotide sequence utilized by the microhomology mediated end-joining DSB repair pathway.

Yet the questions posed by *adprhl1* require more precise dissection than can be achieved by gene knockdown experiments alone. The MO work could never reveal sub-domains within Adprhl1 that are necessary for its function. It was for this reason we turned to CRISPR/Cas9 technology to generate mutations across the *Xenopus adprhl1* locus. By studying the consequences of mutation at defined points within the gene, it was hoped critical amino acids would be identified in Adprhl1 that support ordered myofibril assembly in the heart.

### Optimizing CRISPR/Cas9 gene mutation for phenotype discovery in X. laevis embryos using tyrosinase knockout

Harnessing the activity of the bacterial adaptive immunity system of CRISPR and Cas genes has revolutionised efforts to introduce precise, targeted changes to the genomes of cells and experimental model animals (Reviewed Naert and Vleminckx, 2018; Tandon et al., 2017). The key to its success is the simplicity by which the Cas9 endonuclease can be programmed to cut a specific DNA sequence using a synthetic guide-RNA (Jinek et al., 2012). Endogenous genome repair mechanisms will rejoin any double-strand break (DSB) that occurs within the cell nucleus. If no intact copy of the targeted DNA is available to direct an accurate repair, non-homologous end-joining will result in a sequence lesion at the site of Cas9 cleavage, such as base pair deletion or insertion (Ceccaldi et al., 2015; Seol et al., 2018). If a modified DNA template is provided, then precise gene editing can even be achieved, say to replicate a defined mutation that is observed in human disease.

In order to optimize a method for CRISPR experiments in *Xenopus*, like many studies (Guo et al., 2014; Wang et al., 2015), we used knockout of the *tyrosinase* embryo pigmentation gene to test several Cas9 RNAs along with a commercial Cas9 protein preparation in *X. laevis* (Supplementary S6, Supplementary Methods). Cas9 RNA injection achieved 46% of tadpoles that were complete albinos or contained just a few pigmented cells (Supplementary S6A-C). Significantly, by switching to EnGen^®^ *Spy* Cas9 protein and preloading the *tyr* gRNAs, the proportion of completely albino tadpoles was increased beyond 80%, confirming the enhanced effectiveness of protein injection compared to RNA for one-cell stage delivery of the nuclease activity (Supplementary S6A, B). This rate of *tyr* albinism far exceeded what had been previously recorded for G0-generation tadpoles.

### Identification of adprhl1 gRNAs that cause defective heart development

For knockout phenotype discovery in G0-generation animals, the position of the gRNA within the targeted gene is paramount. The *tyr* gRNAs, for example, hybridize near the start of the coding sequence. At this site, frame-shift mutations produce non-functional alleles while subtler in-frame mutations disturb the N-terminal signal sequence of the tyrosinase enzyme. With the *adprhl1* gene, it was unclear where good positions to site gRNAs might be located. The existence of two protein species cast doubt on targeting the consensus 5’-coding region while its current status as a pseudoenzyme undermined the potential of gRNA hybridization to sequences that encode the ancestral site for ADP-ribosylhydrolase activity. A naïve approach was taken with gRNAs designed to each exon of the *adprhl1* gene, while two were chosen specifically to probe the importance of the active site cleft (-e6-1 and -e7-1). The effect of *adprhl1* gRNA plus Cas9 injection on embryo and cardiac morphology is summarized in Fig 3 (and Supplementary S7).

**Fig 6.**
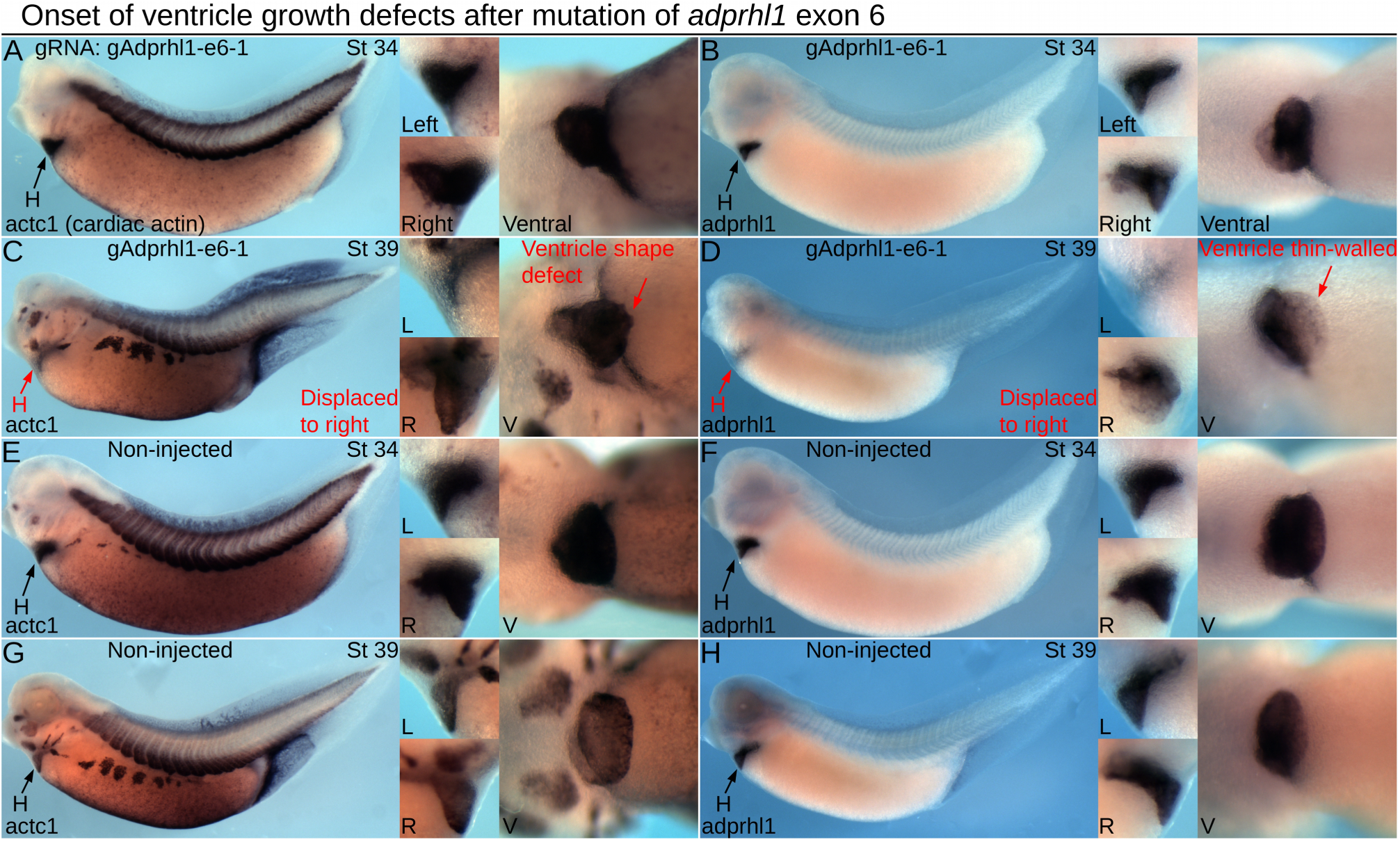
Onset of ventricle growth defects after mutation of adprhl1 exon 6. **A, B:** Expression of *actc1* (heart and skeletal muscle, A) and *adprhl1* (B) mRNAs in stage 34 tadpoles after *adprhl1* exon 6 mutation using the gAdprhl1-e6-1 gRNA plus Cas9. Left-lateral view of tadpole and detail left, right and ventral views of heart region presented. **C, D:** Older stage 39 tadpoles that received the same exon 6 mutation. **E, F:** Sibling non-injected stage 34 tadpoles. **G, H:** Sibling non-injected stage 39 tadpoles. By stage 39, exon 6 mutated tadpoles show ventricle defects and early signs of cardiac oedema. One ventricle (C) is displaced towards the right side of the tadpole and has a malformed apex region. The second ventricle (D) is also displaced and has a faint *adprhl1* signal, suggesting a thinner myocardial wall. Red arrows denote aberrant morphology. H, heart; L, R, V, left, right and ventral views.

**Fig 7.**
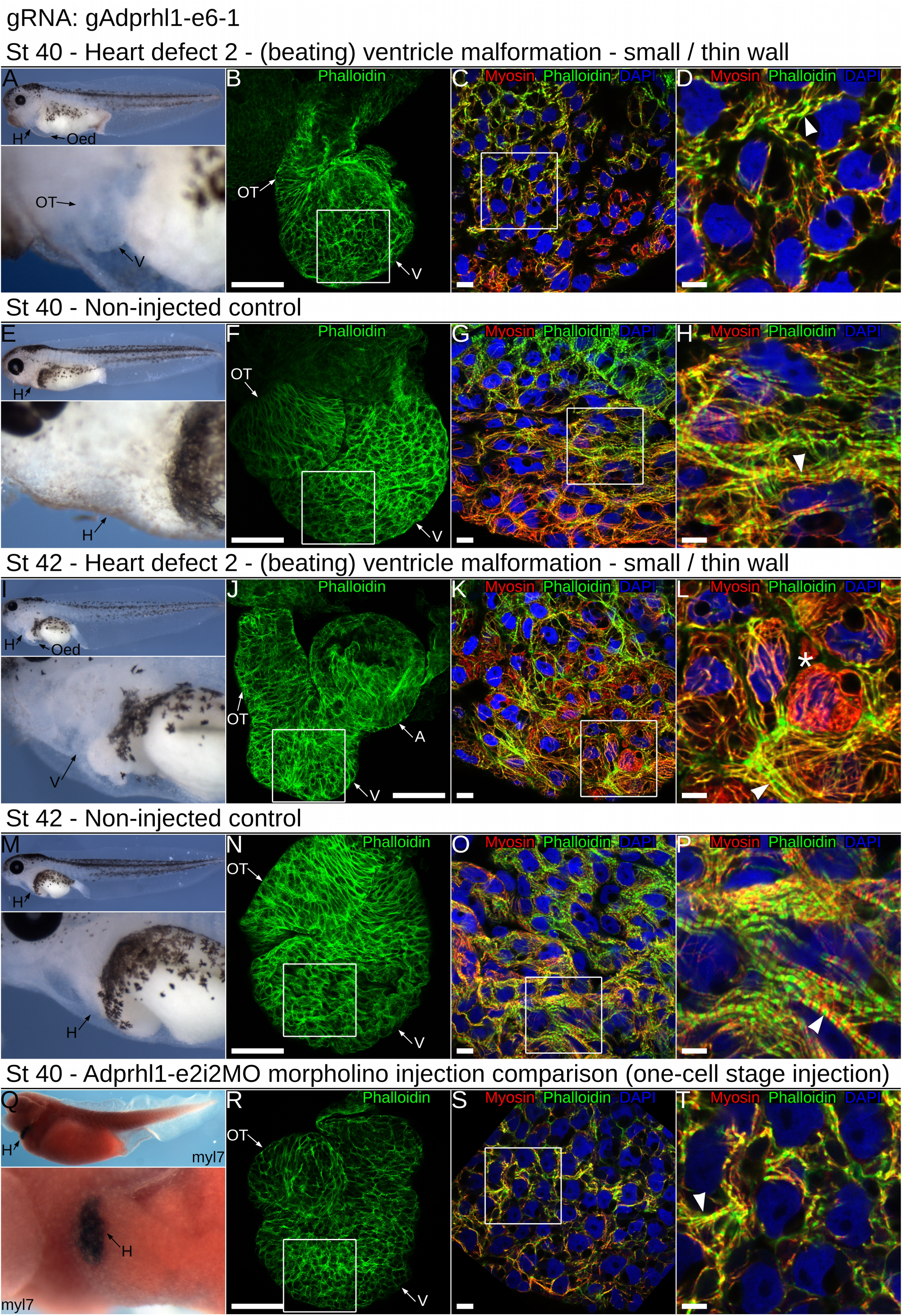
Impaired ventricle myofibril assembly caused by mutation of adprhl1 exon 6. **A:** Developing cardiac oedema typical of a stage 40 tadpole after injection of the gAdprhl1-e6-1 gRNA plus Cas9. Left-lateral view of tadpole and detail of heart region presented. The oedema increased embryo transparency so that the small ventricle became visible at an earlier stage compared to controls. **B-D:** Fluorescence images of the dissected heart ventricle placed with the anterior surface uppermost (B) and displaying merged signals (C, D) of phalloidin actin filaments (green), anti-myosin filaments (red) and DAPI nuclei (blue). The white square (B) denotes the position of the ventricular cardiomyocytes (C) and the white square (C) in turn marks the further magnified image (D). The ventricle is small compared to controls. Cardiomyocytes either have few assembled muscle filaments or contain disarrayed myofibrils with poorly defined sarcomeres (arrowhead, D). Scale bars = 100 µm (B), = 10 µm (C) and = 5 µm (D). **E-H:** Non-injected sibling control stage 40 tadpole and dissected cardiac ventricle. The cardiomyocytes of the ventricle wall assemble myofibrils that extend in a perpendicular to chamber direction (horizontal in the image, H). Discrete sarcomeres are visible (arrowhead, H). **I:** A typical stage 42 tadpole mutated with the - e6-1 gRNA and Cas9. The cardiac oedema and small ventricle are the only overt malformations. **J-L:** The dissected heart has the anterior ventricle surface uppermost, while its aberrant shape positions the outflow tract to the left of the atria after mounting (J)(in controls, the outflow is folded in front of the atria). There is mosaicism amongst the ventricular cardiomyocyte population (K, L), with round non-functional cells (asterisk *, L) and also elongated cells containing disarrayed myofibrils (arrowhead, L). **M-P:** Non-injected sibling control stage 42 tadpole and dissected ventricle. The tadpole epidermis is now transparent allowing simple assessment of cardiac morphology (M). Myofibrils are packed together (O) and Z-disc stripes are prominent (arrowhead, P). **Q-T:** For comparison, a stage 40 tadpole and a dissected heart ventricle obtained after injection of the RNA-splice interfering Adprhl1-e2i2MO morpholino at the one-cell stage. This tadpole was probed for myocardial *myl7* mRNA and its epidermal pigment removed by bleaching. Left-lateral view and ventral detail (Q). The heart dissected from another MO injected tadpole is small (R), with myofibril disarray (S, arrowhead, T) that is comparable to the CRISPR targeted animals. Note, injection of the MO at the one-cell stage to match the CRISPR experiments yielded a slightly milder cardiac phenotype compared to the previous four-cell stage dorsal blastomere injections due to lower MO concentration apportioned to heart forming tissue (compare Fig 2#2D, E). Oed, oedema; H, heart; V, ventricle; OT, outflow tract; A, atria.

There was wide variation in the ability of the different gRNAs to disturb heart formation. Many gRNAs had modest effects, with only a few tadpoles in an injected cohort (<15%) developing cardiac oedema. Nonetheless, two gRNAs did induce oedema and heart ventricle malformations. The gAdprhl1-e3-1 gRNA produced a 30% rate of tadpole heart defects while the most active, gAdprhl1-e6-1, caused a cardiac malformation in over 43% of tadpoles (Fig 3B). The type of abnormality was also consistent amongst affected tadpoles. Hearts produced after -e6-1 gRNA injection typically could beat but developed a small, thin-walled ventricle (see following section) that became increasingly dilated as the tadpole grew. The observed phenotype frequency was lower for *adprhl1* compared to that obtained for *tyrosinase* gene knockout, but did approach the activity of the *adprhl1* RNA-splice interfering MOs when they were injected at the one-cell stage. Moreover, aside from the heart, no other morphological defects were observed consistently in tadpole cohorts. The apparent activity of gAdprhl1-e6-1 was exciting, since this gRNA was one of the two designed to probe the ancestral active site. It hybridized to an exon 6 sequence that encodes a loop between two helices in the substrate binding cleft (see Discussion).

To determine why some gRNAs were more effective than others, we examined the *adprhl1* genomic sequence surrounding gRNA targeted sites. We selected the -e3-1, -e4-1 and -e6-1 gRNAs for further analysis. Each of these gRNAs contained the optimal 20 nucleotides of gene-specific sequence and an identical adjacent PAM (Fig 3A). S- and L-homeolog *adprhl1* DNA was PCR amplified from individual tadpoles that represented all the heart defect and normal morphology phenotypic groups. Sanger DNA sequencing of cloned isolates was then analysed for each gRNA experiment. Sequence results for the three targeted exons are presented in order. Moreover, direct next generation sequencing of a small PCR amplicon was also utilized for analysis of the most penetrant exon 6 mutation.

### Exon 3 mutation reveals Cas9 endonuclease operating with 100% efficiency and cellular mosaicism of resulting alleles

We compared 342 sequences across exon 3 obtained from 23 tadpoles that had received the -e3-1(S+L) gRNA. Alignments revealed that every sequence contained a lesion at the gRNA hybridization position whereas all sequences from a non-injected control tadpole were wild type (Supplementary S8 and S9A). Small nucleotide deletions were most common, with some insertions and a missense mutation also observed. At this site, a mixture of two gRNA molecules prepared to both S- and L-loci had been used, resulting in both homeologs being effectively mutated (Supplementary S9A). Thus injection of CRISPR reagents at the early one-cell stage showed the Cas9 endonuclease operating reproducibly with 100% efficiency in *Xenopus* embryos. Within each tadpole, the number of distinct sequences provided an estimate of the rate of Cas9 action. The highest number detected was 6 different S-locus sequences, although four or fewer was more common (Supplementary S9B). This level of sequence complexity indicated DSB formation and subsequent repair continued beyond S-phase at the one-cell stage and occasionally occurred in separate blastomeres at the two-cell stage. Each injected tadpole therefore contained a mixture of differently mutated sequences and some mosaicism of allele distribution must have occurred across individual cells.

**Fig 8.**
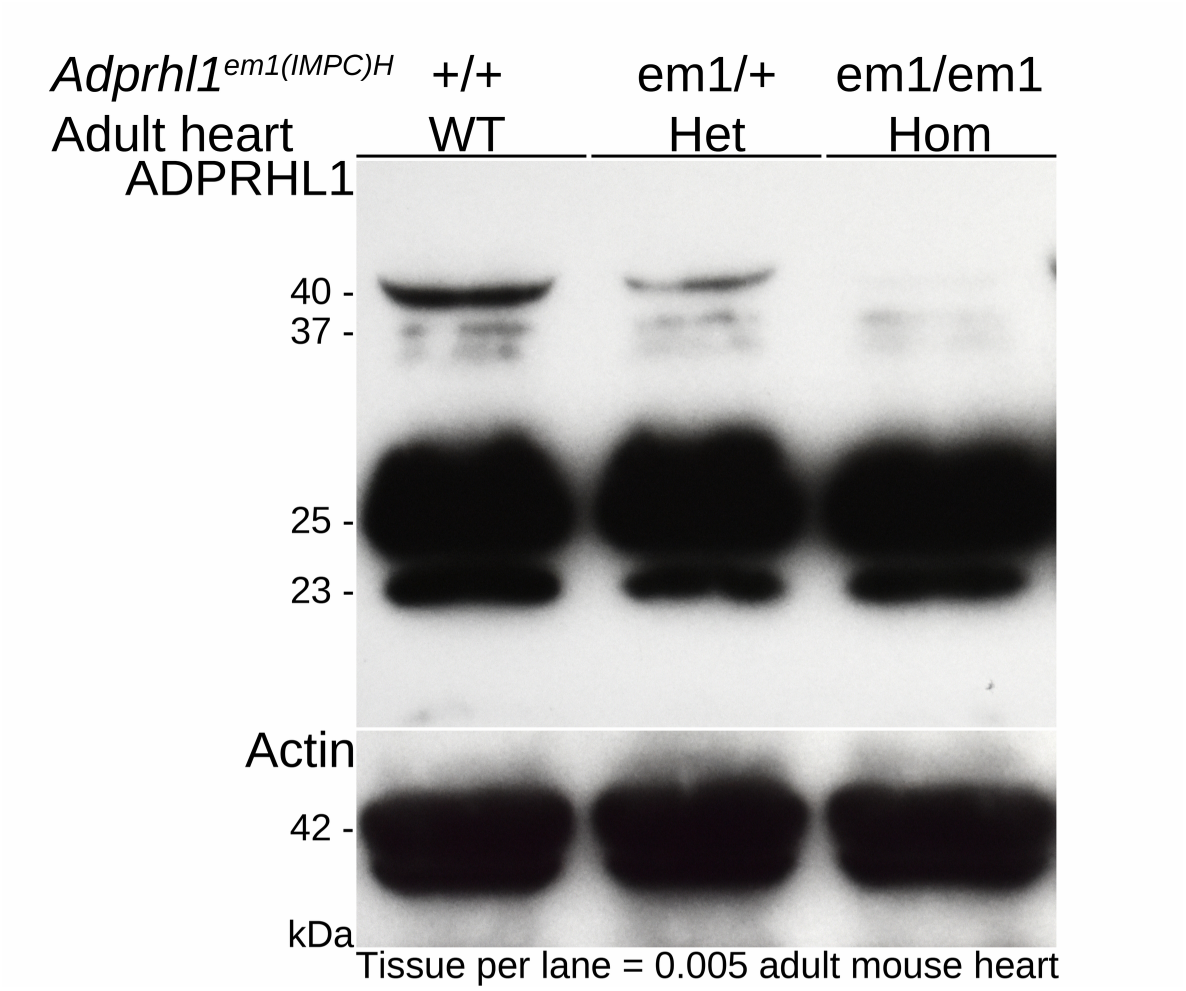
Mice lacking Adprhl1 exons 3 and 4 are normal - They still produce 25 and 23 kDa ADPRHL1 proteins. Western blot detection of ADPRHL1 protein from individual adult mouse hearts carrying the *Adprhl1*^*em1(IMPC)H*^ allele (*em1*). The 40 kDa ADPRHL1 protein was clearly lost from the *em1* homozygote heart and no new species appeared in its place. Significantly though, two other major cardiac ADPRHL1 species, 25 and 23 kDa, were unaffected by the *em1* deletion. Actin detection was used to normalize the samples. WT, wild type; Het, heterozygote; Hom, homozygote. Note that additional, fainter ADPRHL1 protein species were detected around 37 kDa (and 70 kDa - not shown), although their relative abundance was inconsistent in different western blot experiments.

**Fig 9.**
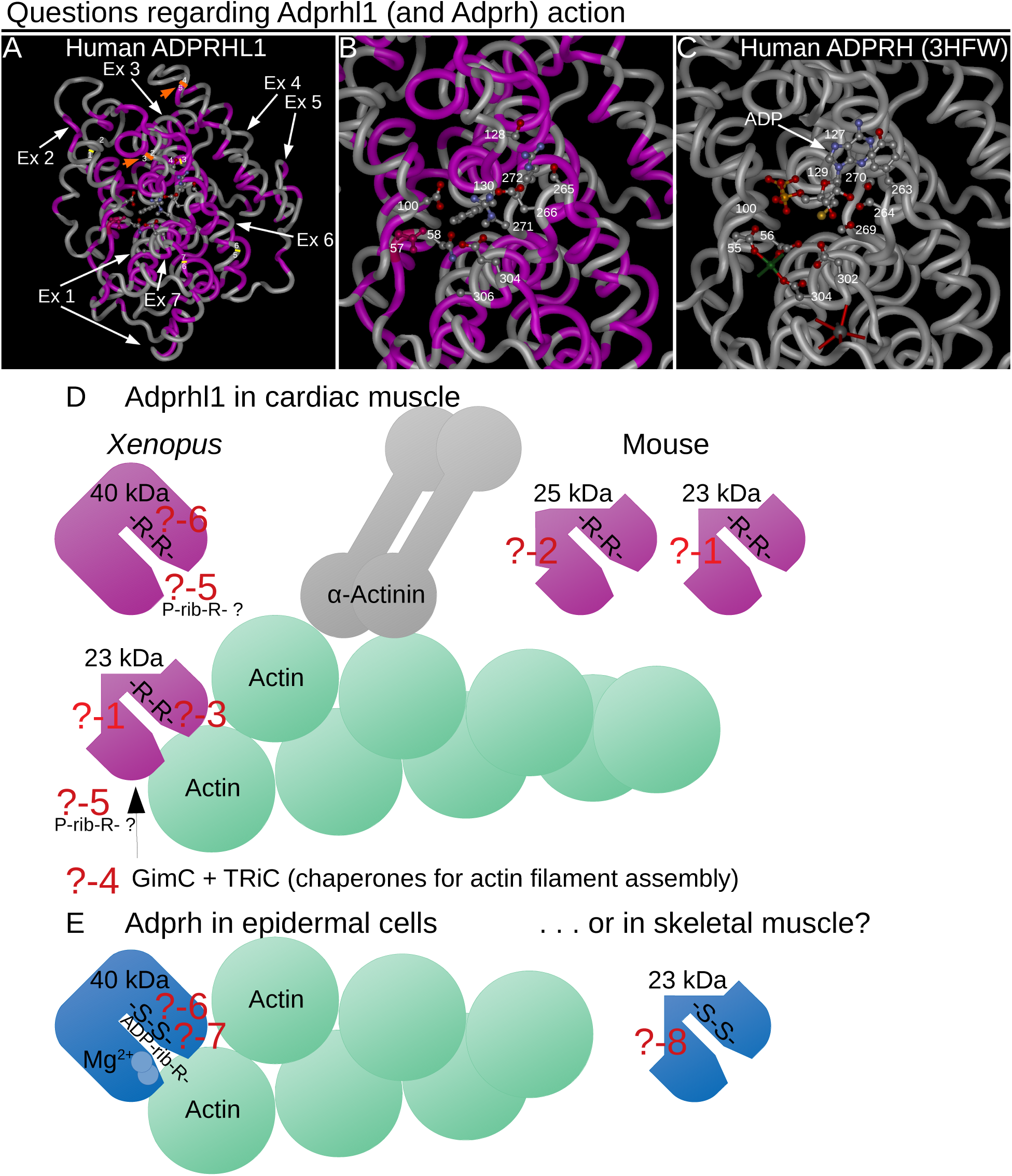
Questions regarding Adprhl1 (and Adprh) action. **A-C:** Structural model of human ADPRHL1 with protein backbone drawn as a tube (A) and magnified active site region (B) (Smith et al., 2016) that is based on the solved crystal structure of human ADPRH (3HFW) (C-active site shown only) (Kernstock et al., 2009). Amino acids that are common to both proteins are coloured magenta (A, B). White arrows indicate the contribution made by each *ADPRHL1* exon to the sequence, orange arrows highlight the positions encoded by the exon-2-3 and exon-4-5 boundaries, with yellow lines marking the remaining exon borders (A). The translated exon-2-3 and 4-5 boundaries reside close to each other and have parallel alignment. Thus the smaller 23 kDa ADPRHL1 form that lacks aa sequence from exons 3-4 could conceivably retain a similar protein fold. Select aa side chains within the active sites are shown as ball and sticks models. For active enzyme ADPRH, aspartates-55, 56, 302, 304 (Mg^2+^ coordination and catalysis), Cys129, Tyr263, serines-264, 269, 270 (substrate binding cleft) and the location of glycines-100, 127 are shown (C). A molecule of ADP occupies the active site, along with one of the two Mg^2+^ cations (green sphere) and a K^+^ ion (grey sphere) (C). For cardiac ADPRHL1, the corresponding active site residues are mostly changed. Asp57 (conserved and coloured red), Asn58, Glu302, Ala304 (no cation coordination), Phe130, Ser265, Glu266, arginines-271, 272 (changed substrate cleft), plus Asp100, Ser128 are shown (B). With this altered active site, ADP cannot be forcibly docked into the ADPRHL1 model. **D, E:** A deeper understanding of Adprhl1 action is required in order to describe how myofibrillogenesis is linked to chamber outgrowth in the embryo. This diagram illustrates several questions that will need to be addressed: **?-1:** What is the precise composition of the smaller 23 kDa Adprhl1 protein? The mouse *em1* allele has already provided some information. It does not contain exons 3-4 encoded sequence but will include exons 5-6 sequence recognised by the peptide antibody. **?-2:** What is the composition of the additional 25 kDa mouse Adprhl1 protein? It is likely to be related to the 23 kDa species. **?-3:** Which sequences localize Adprhl1 to Z-disc/actin filament barbed end boundaries and what myofibril components does it associate directly with? The localization is observed when using an N-terminal epitope tag but not with the exons 5-6-specific antibody. The epitope for the peptide antibody is possibly obscured when Adprhl1 associates with myofibrils. **?-4:** Might Adprhl1 cooperate with the chaperones and co-chaperones that fold and assemble actin filaments? GimC and TRiC are among factors known to contribute to actin dynamics. There are many additional components of the functional sarcomere unit of myofibrils that Adprhl1 could interact with. **?-5:** Does Adprhl1 retain binding activity for a post-translational modification that is related to ADP-ribose? For example, an ADP-ribose modification can be partially degraded to a smaller phospho-ribose group by a pyrophosphatase or phosphodiesterase reaction. **?-6:** Could the substrate binding clefts of Adprhl1 and Adprh also be targets for ADP-ribosylation? Comparison of the di-arginine versus di-serine residues present in the active sites of the two proteins. Di-arginine is common among verified sites for ADP-ribosylation on arginine side chains. Moreover, serine is another acceptor site for ADP-ribosylation that can be hydrolysed by the action of Adprhl2. **?-7:** Does the active enzyme Adprh contain sequences that provide specificity for particular target proteins, in addition to the binding and hydrolysis of ADP-ribosylated arginine? In the illustration, Adprh is depicted acting on ADP-ribosylated actin, one of a number of known (arginine acceptor) targets of bacterial toxin ADP-ribosyltransferases. Could domains adjacent to the active site help stabilize the interaction with particular modified target proteins. Might this be the activity that is common to both Adprh and Adprhl1. **?-8:** Could smaller protein forms of Adprh also exist, for example in skeletal muscle? There could yet be analogy between 23 kDa Adprhl1 action in cardiac muscle and Adprh in skeletal muscle. However, evidence against this comparison would be that exon structure is not conserved between the two family members.

Cas9 cut the targeted DNA completely. Whether a tadpole would reveal a developmental defect due to *adprhl1* gene inactivation depended on the specific details of the DNA strand repairs that occurred. To simplify presentation of the sequence data, a numerical code was assigned that graded each mutation according to the type and size of amino acid sequence modification it encoded (See Materials and Methods). This genotype score scale ranged from 01 for the most severe frame-shift and nonsense mutations, 02 to 05 for in-frame amino acid changes of decreasing size, with score 06 reserved for the natural *Xenopus* Adprhl1 primary sequence. After mutation by -e3-1 gRNA, over 85% of the sequences belonged to the 01 category that would truncate the full length protein product. These frame-shift mutations accounted for the majority of sequences found in tadpoles with heart defects, suggesting an intact exon 3 is an essential component of *adprhl1* mRNA in *Xenopus* (Supplementary S9A, B).

More subtle, in-frame mutations were present at low frequency in most embryos. It was the nature of these mutations that actually correlated with the cardiac phenotype. For example, whenever the p.(Lys145Ile) missense substitution occurred, the tadpole had preserved a normal heart morphology (Supplementary S9B-see tadpoles#20170406004, #20170406024). The substitution must not affect Adprhl1 activity and therefore its presence in a (mosaic) proportion of the mutated cardiac progenitor cells enabled heart formation to proceed. We were less certain of the consequence of small amino acid deletions or insertions at this position. Triplet deletions may have preserved heart function whereas the presence of insertions was biased towards animals with defective hearts (see Supplementary S9B).

Overall, targeting *adprhl1* exon 3 caused complete mutation and a 30% rate of developmental heart abnormalities. The presence of substitutions, in-frame deletions or insertions in nearly all embryos prevented a uniform defective knockout phenotype from prevailing at the G0-generation because some of these mutations retained activity. We did experiment with injecting additional gRNAs that hybridized close to -e3-1 to induce larger deletions between two neighbouring DSB positions (Fig 3A). This combinatorial gRNA approach was partially successful and did increase heart defect frequency (Fig 3B, Supplementary S7), but only targeted the L-locus and never eliminated the genotype score 04-05 alleles (Supplementary S9A).

### Exon 4 mutation is incomplete so rarely causes heart defects

We initially compared 93 sequences from 5 tadpoles that had received the -e4-1 gRNA. This time, specific lesions at the gRNA position were detected in only 82% of sequences and normal alleles persisted in each of the tadpoles (Supplementary S10 and S12). Combinations of -e4-1 with adjacent -e4-2(S+L) or the more distant -e4-3(S+L) gRNAs were then tested (Fig 3A, Supplementary S7). Alignment of a further 171 sequences showed the -e4-1 plus -e4-2 mixture induced deletions between the two DSB sites but again, a few wild-type sequences remained (Supplementary S11 and S12A). With two gRNAs, the frequency of heart abnormalities increased but never matched the level observed for exon 3 (Supplementary S7).

### Exon 6 mutation is biased towards in-frame repairs yet still causes the highest rate of heart defects

For exon 6, 500 DNA clones were examined from 16 tadpoles that had received the -e6-1 gRNA. Alignments showed 499 had defects at the gRNA site and just one retained a wild-type sequence (Fig 4, Supplementary S13). The number of distinct sequences present within each tadpole was similar to that observed for exon 3 mutation (Fig 5B).

The most striking feature of targeting exon 6 was the disproportionately high, 69% frequency of in-frame mutations (Fig 5A). Sequences with 3, 9, or 12 bp deletions constituted over half of the in-frame mutated alleles and were found in all animals (Fig 4, Supplementary S13). A 1 bp missense substitution, 6, 18 and 24 bp deletions, plus some 3 and 6 bp insertions were also identified. The common mutations corresponded to specific deletions of the Adprhl1 protein sequence (264SSEGRGGRRGH274): a single amino acid loss p.(Arg272del), loss of 3 aa p. (Arg271_Gly273del), or loss of 4 aa p.(Gly270_Gly273del) (Fig 5B). The stereotyped deletions resulted from the pattern of short direct repeats of nucleotide sequence found at this site that were utilized for cellular DSB repair by the microhomology mediated end-joining pathway (MMEJ or Alt-NHEJ) (Fig 5D) (Seol et al., 2018). Each of the three common deletion mutations must have eliminated Adprhl1 activity and together, were responsible for the cardiac malformations caused by the -e6-1 gRNA. Even a conserved p.(Arg271Lys) substitution was obtained from a tadpole with a typically malformed ventricle, suggesting it too lacked function (Fig 5B-tadpole#20181121010).

The same in-frame deletions occurred in tadpoles with oedema and heart defects, but also in examples where the heart had apparently formed normally. One wild-type Sanger sequence was additionally detected in an animal with a normal heart (Fig 5B-tadpole#20181121024). We suspected that increasing the number of sequences at the gRNA site would reveal more wild-type alleles and explain the presence of the normal tadpoles in the experiment. Next generation sequencing (NGS) was employed to increase the depth of mutation analysis using a small PCR product for both the S- and L-loci (Materials and Methods). NGS reads contained the same prevalent mutations in the tadpoles that had been identified by Sanger sequencing, albeit with a higher level of sequence variant complexity. Additionally, of the 16 mutated tadpoles screened, wild-type alleles were detected in all 5 animals that had preserved a normal heart morphology but in only 3 of the 11 tadpoles with heart defects (Fig 5B). Of those three, given the mosaic distribution of mutations within the animals, if wild-type sequences persisted in ventral blastomere-derived tissues, they would not be able to contribute to cardiac development due to the ordered cell lineage hierarchy in *Xenopus* embryos (Sive et al., 2000).

There is a marked contrast between the results obtained for exon 6 mutation versus exon 3. At exon 3, the presence of in-frame mutations at low frequency can help protect heart formation. Whereas for exon 6, small in-frame deletions predominate and are actually responsible for the cardiac phenotype observed in 43% of the tadpoles. A model of Adprhl1 protein structure illustrates the position the exon 6 deleted amino acids should occupy within a loop of peptide backbone that lies at the centre of the ancestral ADP-ribosylhydrolase site (Fig 5C) (Smith et al., 2016). Within this sub-domain, an interesting comparison can be made between different members of the (pfam03747) protein family. Where the active enzyme ADP-ribosylhydrolase (Adprh) has di-serine residues that interact with the adenosine-ribose moiety of a substrate ADP-ribosylated protein, in Adprhl1 these amino acids are instead changed to di-arginine (Adprh SYSGWGG<u>SS</u>GH, Adprhl1 264SSEGRGG<u>RR</u>GH274). The unique loop sequences of active enzyme versus cardiac pseudoenzyme are absolutely conserved across vertebrate species from frog to man (see Discussion). Thus the -e6-1 gRNA is effective because the targeted exon 6 sequence translates into a critical Adprhl1 di-arginine motif where amino acid changes are not tolerated. Crucially, it reveals that despite lacking catalytic activity, it is the active site of Adprhl1 that performs an essential role during embryonic cardiogenesis.

### Adprhl1 protein is detected within exon 6 mutated hearts

Because of the in-frame deletions observed at exon 6, it was likely that tadpoles with defective hearts were still competent to produce a near full length Adprhl1 protein. Indeed by western blot, cardiac proteins extracted from the malformed hearts of -e6-1 gRNA injected animals contained both 40 and 23 kDa Adprhl1 similar to control hearts (Supplementary S14).

### A single nucleotide change to the exon 6 gRNA sequence abolishes its activity

Hybridization between the Cas9-gRNA complex and the target DNA site is essential for double-strand endonuclease digestion. Annealing of the gene-specific element of the gRNA (the spacer region) to its target, the opposite DNA strand to the PAM-containing strand, proceeds in a 3’ to 5’-direction (away from the PAM sequence). A perfect match of the 8-10 bases at the 3’-portion of the gRNA spacer (known as the seed sequence) is absolutely required for successful cleavage and the DSB occurs 3-4 bp upstream of the PAM (Jiang et al., 2015; Wu et al., 2014). In order to demonstrate the specificity of the interaction required for *adprhl1* mutation, we synthesized two control exon 6 gRNAs that contained a single or double nucleotide mismatch located at the putative DSB site (Fig 3A). These control gRNAs injected under identical conditions exerted no effect on heart formation and the resulting tadpoles developed completely normally (Fig 3B).

### Myofibril assembly defects observed in adprhl1 exon 6 mutated hearts

After identifying the mutations produced within *adprhl1* exon 6, we next examined the aberrant heart phenotypes of the affected tadpoles in finer detail. Heart formation was monitored on five consecutive days from stage 34 through to 44. This developmental period covers looping morphogenesis of the early heart tube in addition to outgrowth and maturation of the ventricle chamber. At stage 34, there was little to distinguish the cohort of mutated heart tubes from controls. The size of the differentiated cardiac muscle tissue shown by *actc1* expression, the detection of *adprhl1* mRNA, plus the extent of looping all appeared normal (Fig 6A, B, E, F). The following day at stage 39, some differences could be detected. Among the exon 6 mutated tadpoles, the ventricle chamber was frequently smaller than controls, displaced from the ventral midline and oriented incorrectly with regard to the outflow tract position (Fig 6C, D, G, H). In some examples, the *adprhl1* mRNA signal in the ventricle also appeared weaker.

A central process during cardiac ventricle growth is the assembly of muscle motor proteins within cardiomyocytes into functional myofibrils that are aligned with distinctive perpendicular or parallel to chamber orientations (see Introduction). Immunological detection of myosin and actin filaments assessed these features of ventricle morphology and linked myofibrillogenesis within tadpoles carrying exon 6 mutations (Fig 7, Supplementary S15). Each animal harbored a subtly distinct complement of *adprhl1* mutations and this was reflected in the range of aberrant chamber phenotypes observed. The severity of myofibril defects even differed between adjacent cardiomyocytes within a single heart indicating a functional consequence to the cellular mosaicism. Figure 7 presents typical examples of tadpoles of an intermediate phenotype severity, developing with small ventricles that retained some heart beat function. Once the final phenotype classes were assigned at stage 44, Supplementary S15 shows tadpoles representing extremes from the range of hearts considered abnormal, from the most severely affected inert ventricle to the mildest malformation observed in the beating ventricle group.

By stage 40, cardiac oedema formation indicated which of the mutated tadpoles had impaired circulation (Fig 7A). Upon dissection of these hearts, the ventricles were small compared to controls (Fig 7B). Their cardiomyocytes either had few assembled muscle filaments or contained short, disarrayed myofibrils with poorly defined sarcomeres (Fig 7C, D). Non-injected sibling control ventricles contained long, perpendicular myofibrils with discrete sarcomere structure at this stage (Fig 7E-H).

Within individual stage 42 mutated hearts, the mosaicism amongst the ventricular cardiomyocyte population became more pronounced (Fig 7I-L). Many cells remained round, lacked myofibrils and instead contained a dense mesh of muscle myosin protein stain (asterisk *, Fig 7L). Other cardiomyocytes that retained some competency to assemble myofibrils extended filaments that enveloped the round cells. However, as a consequence of the disruption, there was no clear order to the direction of filament extension (Fig 7K). Viewed at high magnification, there was considerable disarray, with branched myofibrils and poor alignment of the myosin and actin filaments into ordered sarcomeres (arrowhead, Fig 7L). Within control stage 42 ventricles (Fig 7M-P), increased packing together of the perpendicular chamber myofibrils had occurred (Fig 7O). The prominent green stripe of actin at the Z-discs provided evidence of the maturing sarcomere structure (arrowhead, Fig 7P).

At stage 44, tadpoles classed as having the strongest heart defect with an inert ventricle had large oedemas but no other discernible malformations (Supplementary S15A). Ventricle growth had failed to the extent that the heart still resembled a primitive tube that also lacked trabeculae ridges (Supplementary S15B, C). Round cardiomyocytes and the myofibril defects persisted as described before (Supplementary S15D-H). Even mutated tadpoles exhibiting the mildest form of heart defects were noteworthy (Supplementary S15I-P). Their cardiomyocytes did contain extensive myofibril networks but they had an unusual appearance (Supplementary S15L-O). The periodicity of actin filaments was equally spaced rather than having the characteristic striated pattern. Moreover, at the resolution available, the actin signal never showed a concentrated stripe to mark the existence of Z-discs (Supplementary S15P). By contrast, all control hearts examined had a consistent morphology (Supplementary S15Q-X). In particular, all the cardiomyocytes within the ventricular myocardial wall assembled myofibrils at an equal rate and no round cells, or developmentally delayed cells were detected (Supplementary S15T-X).

The defects to chamber myofibrillogenesis observed after *adprhl1* exon 6 mutation were compared to that previously reported for *adprhl1* gene knockdown using the Adprhl1-e2i2MO morpholino reagent (Smith et al., 2016). There is a remarkable consistency to the results. Small, inert hearts formed after MO injection at the one-cell stage. When assayed at stage 40, ventricle growth was impaired and their cardiomyocytes produced only small numbers of myofibrils, which were characteristically short, malformed (branched) and with no chamber-type alignment pattern (Fig 7Q-T).

### Mice lacking Adprhl1 exons 3 and 4 are normal - They still produce 25 and 23 kDa ADPRHL1 proteins

Thus far, *Xenopus* is the sole vertebrate model species to have revealed an essential role of *adprhl1* in heart development, despite the clear evolutionary conservation of gene sequence and cardiac gene expression. In mouse, a definitive *Adprhl1* gene knockout has not been reported. An *Adprhl1*^*em1(IMPC)H*^ allele (*em1*) has been produced as part of the international mouse phenotyping consortium (www.mousephenotype.org) (Dickinson et al., 2016). These homozygote *em1* mouse embryos develop normally and the detailed phenotyping pipeline did not reveal any significant adult deleterious traits. The *em1* allele is not however a complete deletion of the *Adprhl1* gene. Rather *em1* harbours a 1087 bp deletion that removes *Adprhl1* exons 3 and 4. Both exons encode an intact number of codons (126 and 141 bp) and 89 amino acids are theoretically lost from the full length protein.

To clarify the nature of this allele, we obtained *em1* mice and examined ADPRHL1 protein production within their hearts (Fig 8). The 40 kDa ADPRHL1 did indeed show a dose-dependent loss in heterozygote and homozygote *em1* adult hearts but there was no linked appearance of a new species equivalent to an exons 3-4 deleted form. There were nonetheless two other abundant protein species identified by the ADPRHL1 antibody at 25 and 23 kDa. Significantly, these smaller ADPRHL1 proteins were unaffected by the *em1* deletion. It should be noted that the two smaller proteins had also been observed previously in E11.5 embryonic mouse hearts (Smith et al., 2016). Thus the *em1* data proves the full length ADPRHL1 protein is not actually required for heart formation in mammals but also shows that the *em1* deletion is probably not a null mutant allele. It highlights the potential importance of the smaller ADPRHL1 protein forms and provides valuable information on their likely composition. The 25 and 23 kDa species do not contain exons 3-4 encoded sequence but will include the exons 5-6 sequence recognised by the antibody.

Given the results reported here for *Xenopus adprhl1* mutation, we anticipate a true mouse *Adprhl1* gene knockout would also show comparable severe defects in embryonic cardiogenesis. By identifying the di-arginine containing peptide loop that is conserved across species whose exon 6 sequence is amenable to targeting by CRISPR/Cas9, our *Xenopus* experiments can help inform revised attempts to document *Adprhl1* gene inactivation in mammalian systems.

## Discussion

### CRISPR/Cas9 adprhl1 gene knockout in G0-generation embryos

We have used CRISPR/Cas9 technology to induce mutations across the *Xenopus adprhl1* gene in order to build on a previous study of morpholino-mediated *adprhl1* expression knockdown. Growth of the heart ventricle in embryos and particularly the assembly of cardiac myofibrils was dependent on *adprhl1* but how it acted was uncertain and was complicated by the existence of two Adprhl1 proteins. MO reagents that interfered with *adprhl1* RNA-splicing depleted both the expected 40 kDa protein and also the smaller 23 kDa species (Smith et al., 2016). Using expression of transgenes, we now show the 40 kDa protein only part rescues MO myofibril defects. Meanwhile, other MOs designed to selectively inhibit translation initiation of the 40 kDa Adprhl1 cause earlier developmental malformations that overshadow any heart abnormalities. Their poor specificity presents a barrier to further progress using antisense experiments.

CRISPR/Cas9 is a transformative technology that allows the genomes of cells and experimental model animals to be precisely targeted, altered and scrutinised (Jinek et al., 2012). It is a powerful tool for developing applications of gene editing (Reviewed Naert and Vleminckx, 2018; Tandon et al., 2017). In a simple strategy for targeted gene mutation, we introduced DSBs at different exonic locations of the *Xenopus laevis adprhl1* gene by injecting guide-RNAs and the Cas9 endonuclease into newly fertilized embryos. Subsequent genome repair by non-homologous end-joining results in a DNA sequence lesion at the site of Cas9 cleavage. Repair occurs simultaneously with ongoing DNA synthesis and cell division, so variant alleles can have a mosaic distribution among cells as the animals grow. Rapid discovery of a uniform *adprhl1* knockout phenotype in G0-generation embryos thus requires complete DSB cutting of every gene copy present at the one-cell stage and that all resulting variant alleles lack activity. Our optimized method routinely achieves 100% DSB efficiency but it is inevitable that in-frame nucleotide deletions, insertions or substitutions within the *adprhl1* coding sequence will always occur along with mutations that cause catastrophic frame shifts. Therefore, testing many gRNAs distributed across the *adprhl1* locus for their ability to impair heart formation is, in effect, a screen for DSB positions where all favoured repairs yield non-functional alleles and where even the most subtly repaired mutation that preserves an intact reading frame still encodes an inactive protein. Such a sensitive gRNA target site would likely identify a sub-domain of Adprhl1 that is essential for its participation in cardiac myofibrillogenesis.

Many studies of heart formation have shown the embryo has significant regulatory potential in order to produce a functional organ in the face of potentially damaging insults (Drenckhahn et al., 2008). For example, a cardiac progenitor tissue field where only a portion of cells were compromised and unable to contribute to the developing chambers could reconfigure and still produce a normal heart. This consideration is critical for understanding the difference between G0-experiments that target an essential cardiogenic gene versus say monitoring pigment-loss after *tyrosinase* mutation. Yet despite all these potential limitations, we find that a gRNA that promotes *adprhl1* mutation at exon 6 does yield consistent heart myofibril defects.

### Is there a link between Adprhl1, ADP-ribosylation on arginine and actin filament dynamics?

Little was known regarding Adprhl1 action before it was linked to the heart. Sequence similarity defined a small family of vertebrate ADP-ribosylhydrolases (pfam03747), comprising the active enzyme Adprh (sometimes named ARH1, or ADP-ribosyl-acceptor hydrolase), Adprh-like 1 (Adprhl1, or ARH2) and also an enzyme Adprh-like 2 (Adprhl2, or ARH3) (Mashimo et al., 2014). Comparing human homologs, the 357 amino acid ADPRH and 354 aa ADPRHL1 share 46% sequence identity, while ADPRH and ADPRHL2 are 22% identical. Moreover, the evolutionary conservation of Adprhl1 sequence extends to frogs, with *Xenopus* Adprhl1 being 75% identical to human ADPRHL1 and 47% identical to the *Xenopus* species Adprh.

The founding member Adprh was first identified during the study of mono-ADP-ribosylation, a post-translational modification of proteins in which arginine side chains are a frequently used attachment site (Reviewed Laing et al., 2011). Adprh is a cytosolic enzyme that can reverse the modification by cleaving the ADP-ribose linkage and restoring unmodified arginine residues (Moss et al., 1992). Its discovery supported a proposal that cycles of ADP-ribosylation and removal might occur within (animal) cells to regulate target protein function. Protein structures and the reaction mechanism reveal that active ADP-ribosylhydrolases rely on a pair of divalent cations, magnesium for ADPRH and ADPRHL2, that are coordinated by two pairs of aspartate residues located in N- and C-terminal portions of the enzyme (Kernstock et al., 2009; Mueller-Dieckmann et al., 2006). ADPRHL2 contains a distinct flexible substrate binding cleft that enables hydrolysis of ADP-ribosylated serine side chains, as well as poly(ADP-ribose) and O-acetylated-ADP-ribose degradation (Pourfarjam et al., 2018; Rack et al., 2018). However, within the familial active site of Adprhl1, it is the amino acids necessary for catalysis that are changed. Three of the four critical aspartates have been lost in mammalian ADPRHL1, along with key tyrosine and serine residues required to stabilize the adenine and ribose substrate groups (Fig 9A-C, residues of human ADPRH lost are Asp56, Asp302, Asp304, Tyr263, Ser264, Ser269, Ser270, only Asp55 is retained) (Smith et al., 2016). The sequence changes suggest binding of ADP-ribosylated protein substrates and cation-mediated catalysis are both abolished in Adprhl1 and biochemical assays have confirmed the lack of any comparable enzymatic activity (Oka et al., 2006). The term ‘pseudoenzyme’ has been coined to raise awareness of such members of a protein family that have lost essential catalytic residues (Reviewed Ribeiro et al., 2019). As an example, without a functioning active site, pseudokinases have evolved to redeploy a substrate binding activity for a new purpose, or confer allosteric control upon a relative kinase that retains enzymatic activity. So did cardiac Adprhl1 evolve to bind proteins that can also be targets of the active hydrolase? The result of CRISPR-mediated cleavage by the gAdprhl1-e6-1 gRNA and the small in-frame deletions that predominate at this site suggests this is likely. Severe ventricle myofibril defects resulting from disturbed Adprhl1 function occur because of the specific omission of between one and four amino acids from a loop of peptide backbone at the centre of the ancestral ADP-ribosylhydrolase site. The critical Adprhl1 deletion covers the exact structural position where in the active enzyme Adprh, two adjacent serines that support adenosine-ribose substrate binding are located. So while ADP-ribose may not be the correct molecular group, it is the substrate binding cleft of Adprhl1 that fulfils an essential role during heart chamber growth.

Although indirect, two pieces of evidence point towards actin being a shared protein target of both Adprhl1 and Adprh. Previously, an N-terminal epitope tag revealed a direct Adprhl1 association with myofibrils in *Xenopus*, localizing to barbed end boundaries of actin filaments (Smith et al., 2016). We now also know that *Xenopus adprh* mRNA has expression within epidermal cells that function to protect the embryo from external pathogens (Fig 1A). This is significant as it potentially changes the context of Adprh enzyme action in aquatic vertebrates. An epidermal cell activity could counter the invasive ADP-ribosyltransferase (ART) components of toxins from pathogenic bacteria (Reviewed Simon et al., 2014). Arginine-specific toxin ARTs have been studied extensively and the host cell cytoskeleton is a major focus. At present, there are nine distinct toxins identified that selectively ADP-ribosylate actin at a single arginine-177 (equates to Arg179 in muscle α-actins) and this modification triggers the collapse of actin filaments within the stricken cell (Reviewed Aktories et al., 2011). They attack the monomeric form of actin, which then behaves like a capping protein at the barbed end of filaments (Wegner and Aktories, 1988). Arginine-177 sits at the interaction site between opposing subunits of the actin filament when modelled as a two-stranded helix (Holmes et al., 1990; Margarit et al., 2006). The affinity of ADP-ribosylated actin for the barbed end appears ten-fold higher than unmodified actin, but the protruding modification will then prevent further additions to the adjacent strand, while continued dissociation from the pointed end ultimately causes filament depolymerization.

So Adprhl1 likely affects myofibrils through direct binding of actin and if Adprh may liberate actin from toxin mediated ADP-ribosylation, then we could speculate that the filament interaction surface of actin that surrounds Arg177 might be the link that unites the two. Finding unbiased ways to test the binding specificities of the ADP-ribosylhydrolase family in their correct cellular environment will be key to understanding their true evolutionary relationship. The illustration in Fig 9(D, E) summarises outstanding questions regarding Adprhl1 action that need to be addressed.

### Mapping essential regions of the Adprhl1 protein

Separate gRNAs that hybridized to exon 3 and exon 6 sequences of the *Xenopus adprhl1* gene produced developmental heart defects in 30% and over 43% of injected embryos respectively. After exon 6 mutation, myofibril phenotypes in the forming ventricle could be compared to the previous RNA-splicing MO results (Smith et al., 2016). Disruption of *adprhl1* at the one-cell stage did cause mosaic effects likely due to the heterogeneity of gene repairs. Common features included cardiomyocytes that were severely malformed, remaining round shaped and devoid of myofibrils. Other cells that presumably retained residual Adprhl1 function extended low numbers of muscle filaments with defective sarcomere structure. The MO myofibril abnormalities were generally uniform in all cardiomyocytes, reflecting an initially complete Adprhl1 loss that gradually became less effective as the tadpole grew. Both gene knockout and knockdown experiments revealed that absence of Adprhl1 is associated with a fundamental myofibril assembly deficit.

At exon 3, the range of mutations detected was as expected. Without an intact template to direct accurate repair, the majority of DSBs were rejoined with the reading frame shifted so as to interrupt the coding sequence. The presence of in-frame mutations at low frequency in nearly all embryos determined the severity of the cardiac phenotype, with a milder missense p.(Lys145Ile) substitution being found in tadpoles that preserved a normal heart morphology. Cataloguing the mutations at exon 6 nicely illustrated the consequence of short, direct DNA sequence repeats occurring near the gRNA target site. An emergency repair process exists termed microhomology mediated end-joining, where exonuclease digestion of single DNA strands at the DSB liberates the two repetitive sequences to act as a (erronious) strand annealing position for religation of the lesion (Ceccaldi et al., 2015; Seol et al., 2018). The repeats adjacent to the -e6-1 gRNA ensured that repairs were biased in favour of three small in-frame deletions of 3, 9 or 12 bp. Thus exon 6 mutation provided opportune new information, since it linked disturbed Adprhl1 function and ventricle myofibrillogenesis defects to precise amino acid deletions from within the ancestral substrate binding region of the active site cleft.

If Adprhl1 has repurposed a binding affinity from ADP-ribosylhydrolases, then it stands that not all of the familial amino acid sequence would necessarily be required in order to fulfil its new role. The smaller protein identified by the Adprhl1 antibody is conserved in vertebrates; 23 kDa in *Xenopus* while 25 and 23 kDa species exist in mouse hearts. Study of the mouse *Adprhl1 em1* allele has now turned our focus towards these smaller proteins. Homozygous *em1* adult mice are completely normal despite this deletion having removed *Adprhl1* exons 3 and 4, which would ordinarily contribute a whole 89 amino acids to the full length 40 kDa protein. The hearts of *em1* mice did lack 40 kDa ADPRHL1 but the 25 and 23 kDa variants were abundant and unaffected by the deletion. It seems murine 40 kDa ADPRHL1 is a relatively minor protein species that is not actually required for heart formation in mammals. The exact composition of the 25 and 23 kDa ADPRHL1 proteins is not yet known. They do not contain exons 3-4 encoded sequence but will include the exons 5-6 sequence recognised by the antibody.

The conclusions drawn from *Xenopus* and mouse experiments are not identical but there is considerable alignment between the two. The main difference concerns *adprhl1* exon 3, whose size and sequence is conserved across species. Nonetheless, exon 3 frameshift mutations in *Xenopus* were implicated in cardiac defects whereas it is clearly dispensable in the mouse heart. One way to explain the disparity would be if *Xenopus* hearts require both the 40 and 23 kDa Adprhl1 proteins to assemble myofibrils. A true mouse *Adprhl1* gene knockout has yet to be achieved but we anticipate it would show a comparable embryonic cardiac phenotype that depends on the presence or absence of the smaller 25 and 23 kDa ADPRHL1 proteins. Both species point towards the 3’-most exons of the *adprhl1* locus as the necessary focus for further study and *Xenopus* has highlighted the peptide loop encoded within exon 6. Conservation of this loop sequence means it is also amenable to targeting by CRISPR/Cas9 in mouse and in human cells. Species-specific gRNAs can be designed that match the same exonic PAM location and a repeat sequence favouring a 3 bp deletion repair is even present in the mouse. It would be an excellent starting point for attempts at inactivating the *Adprhl1* gene in mammals.

A recent human GWAS of the Icelander population has discovered a link between *ADPRHL1* and heart ventricle function (Norland et al., 2019). They identified 190 sequence variations from a pool of 32.5 million Icelander SNPs and indels that associated with changes in a detailed analysis of electrocardiogram QRS parameters measured in 81 thousand individuals. One encoded a missense (p.Leu294Arg) substitution in ADPRHL1 that additionally associated with genome-wide significance to an automated ECG diagnosis of a left anterior fascicular block (LAFB) conduction defect. LAFB reflects a disturbance to the normal spatial conduction/contraction sequence of the left ventricle. It is striking that this low frequency *ADPRHL1* variant is located within exon 6. The leucine altered is absolutely conserved across vertebrate species and structure models place it in an α-helix immediately underlying the essential loop of the active site. Where the loop of the enzyme Adprh contains di-serine residues, Adprhl1 has instead di-arginine. It is too early to comment on the functional significance of the change, except to mention that auto-regulation is a common feature of ADP-ribosylation enzymes, that both serine and arginine can be acceptors of such modification and that di-arginine and prolyl-arginine are prominent among verified acceptor target sequences (Laing et al., 2011; Matic et al., 2012).

### CRISPR/Cas9 experiments and Xenopus embryos

Amphibian embryos have consistently provided major insights across all scales of biological research. CRISPR/Cas9 technology now allows genomes to be interrogated with such speed and precision that new experimental strategies are possible that complement the traditional strengths of the *Xenopus* model. There is tremendous scope for gene editing in the diploid *X. tropicalis* species to replicate specific gene mutations observed in human disease (Reviewed Naert and Vleminckx, 2018). Stable frog lines and the large embryo spawn size can even enable screens for interventions that might ameliorate mutant defects. Other studies require an instant readout of results from a cohort of Cas9 and gRNA injected embryos, say to assess the consequence of specific genome changes at multiple locations. For G0-generation experiments, *X. tropicalis* and *X. laevis* offer different attributes (Reviewed Tandon et al., 2017). *X. laevis* does have four alleles of a gene to consider but this provides extra protection against potential off-target activity and the slower development gives more time for Cas9 action at the one-cell stage. Its large embryo size is particularly suited to subsequent analysis of gene edited tadpoles using proteomic methods (size of *X. laevis*=3x>*X. tropicalis*=2x>*D. rerio*). For our study, the power of CRISPR/Cas9 over previous strategies for targeted mutation lies in the speed that many gRNAs can be designed, synthesized and tested for activity.

How myofibril assembly underpins ventricle construction is a three-dimensional challenge. The *Xenopus* embryo offers cardiac structural features relevant to complex mammalian heart chambers, a conservation of *adprhl1* gene sequence, external development and manipulation by gene editing. Moving forwards, *Xenopus* can contribute to Adprhl1 biology through systematic addition of in-frame epitope tags to each coding exon. Tagging the endogenously encoded Adprhl1 will map the composition of the different protein forms within the heart and provide the necessary handles to explore its interaction partners. This previously overlooked but potentially pivotal facilitator of cardiac myofibrillogenesis can now be explored in far greater detail.

## Acknowledgements

This work was supported by The Francis Crick Institute, which receives its core funding from Cancer Research UK, the UK Medical Research Council (grant U117562103) and the Wellcome Trust. We thank the Aquatics STP of The Francis Crick Institute for supply of *Xenopus* embryos.

