## Supplementary Figures and Methods for "Defective heart chamber growth and myofibrillogenesis after knockout of *adprhl1* gene function by targeted disruption of the ancestral catalytic active site"

Stuart J. Smith[1], Norma Towers[1], Kim Demetriou[2] and Timothy J. Mohun[1]*.

1. Heart Formation in Vertebrates Laboratory, The Francis Crick Institute, 1 Midland Road, London NW1 1AT, UK.
2. Aquatics STP, The Francis Crick Institute, 1 Midland Road, London NW1 1AT, UK.

* Corresponding author, Tel: +44 20.

### Supplementary Figure Legends.

**Supplementary S1.** *Limited recovery of cardiac myofibril assembly in adprhl1 morpholino injected embryos by transgenic synthesis of recombinant 40 kDa Adprhl1 proteins.*
Experiments that combine *adprhl1* MO knockdown with two distinct transgenes engineered to achieve *adprhl1* over-expression. This is the extended version of Figure 2. It additionally shows the morphology of the experimental heart ventricles and the extent of Adprhl1 protein production within. The extra panels locate the position within each ventricle wall of the high magnification images (that are also presented in Fig 2) that reveal myofibril patterns found in sample cardiomyocytes.

**1A-E:** A stage 41 tadpole and its dissected heart ventricle that was injected with the RNA-splice interfering MO, Adprhl1-e2i2MO, into dorsal (D-2/4) blastomeres. Additionally, it carried binary transgenes to over-express recombinant Adprhl1 protein, consisting of *Tg[myl7:Gal4]* driver and the *Tg[UAS:human$^{1\text{-}52}$-Xenopus$^{53\text{-}354}$ adprhl1]* responder. Left lateral view of head and trunk (A), while the dissected heart was placed with the anterior surface uppermost (B). The white square (B) denotes the position of a detail image of the ventricle (C) and the white square (C) in turn marks the position of further magnified images (D, E). Scale bars = 100 μm (B), = 10 μm (C) and = 5 μm (D, E). Fluorescence images (B-D) show anti-Adprhl1 immunocytochemistry (green), anti-myosin (red) and DAPI-stained nuclei (blue, D). The final panel (E) displays a merge of myosin and phalloidin actin stain, with the phalloidin coloured green to evaluate signal overlap. **2A-E:** A sibling tadpole that received the same Adprhl1-e2i2MO injection but carried only the UAS-responder transgene and hence did not produce excess recombinant Adprhl1. **3A-E:** A double transgenic sibling that synthesized recombinant human-*Xenopus* hybrid Adprhl1 but was not injected with the MO. **4A-E:** From a second experiment, a stage 42 tadpole that was injected with Adprhl1-e2i2MO and carried the *Tg[myl7:Gal4]* driver but a different *Tg[UAS:Xenopus adprhl1(silent 1-282bp)]* responder transgene. This incorporates silent nucleotide changes (synonymous substitutions) to the cDNA sequence in order to partially evade endogenous translational regulation. The heart had a strong MO defect on the anterior ventricle surface, but lower MO concentration and incomplete phenotype towards its right side (4B). **5A-E:** A double transgenic, silent mutation, sibling tadpole that synthesized recombinant *Xenopus* Adprhl1 but was not injected with the MO. **6A-E:** A non-injected sibling control harbouring only the silent mutation responder transgene that did not produce excess recombinant Adprhl1. Paired white arrowheads indicate Z-disc sarcomere positions, orange arrowheads denote non-striated filaments. V, ventricle; OT, outflow tract.

**Supplementary S2.** *Over-expression of recombinant 40 kDa Adprhl1 does not yield extra 23 kDa Adprhl1.*
Western blots of transgenic tadpole hearts that carried stable lines of the *Tg[myl7:Gal4]* driver and one of a series of *Tg[UAS:adprhl1]* responders designed to over-express variants of 40 kDa Adprhl1 protein. Stage 43-44 heart extracts were probed with Adprhl1 antibody, with Actin detection used to normalize the samples. Hum corresponds to a *Tg[UAS:human ADPRHL1]* responder that induces large-scale synthesis of human-species ADPRHL1 protein in tadpole hearts. This transgene cDNA sequence is sufficiently different from *Xenopus* to evade the endogenous translational control mechanism that normally limits the production of Adprhl1. Hyb corresponds to the *Tg[UAS:hum$^{1\text{-}52}$-Xen$^{53\text{-}354}$ adprhl1]* responder that over-synthesizes a human-*Xenopus* hybrid form of Adprhl1 that also escapes translational control. Xen denotes a *Tg[UAS:Xenopus adprhl1]* transgene. Using unmodified *Xenopus adprhl1* cDNA, transgene mRNA transcription is activated but no additional recombinant protein accumulates (Smith et al., 2016). WT equates to control (wild type) hearts.

On a separate gel with a higher signal exposure, Xen(silent) denotes the *Tg[UAS:Xenopus adprhl1(silent 1-282bp)]* transgene containing silent nucleotide changes to the 5'-*Xenopus* cDNA. This transgene partially evaded endogenous control and recombinant Adprhl1 accumulated in a fraction of the cardiomyocytes up to stage 42. However, there is a technical barrier to performing western blot analysis of transgenic hearts at these early stages. The time required to identify double-positive embryos by the transgenes' marker eye fluorescence and subsequent sample preparation in the numbers necessary to obtain a signal is prohibitive. The silent mutation transgene gave a transient protein induction (Smith et al., 2016) and no additional Adprhl1 was detected in this sample prepared from stage 44 hearts.

In hearts that synthesized excess 40 kDa hybrid Adprhl1, there was no commensurate increase in the abundance of the 23 kDa protein detected by the Adprhl1 antibody. It suggests the 23 kDa Adprhl1 species is not a processed fragment of the full length protein. Of course, this conclusion depends on the hybrid and natural *Xenopus* forms of Adprhl1 behaving equivalently. There are just 21 amino acid differences between the two variants.

**Supplementary S3.** *Excess Adprhl1 production does not trigger cell proliferation, nor cause cell death.*
Tadpole heart ventricles with transgenic over-expression of 40 kDa Adprhl1 protein combined with markers of cell division (A-D) and cell death (E-R). The cardiac *Tg[myl7:Gal4]* driver and *Tg[UAS:Xenopus adprhl1(silent 1-282bp)]* responder transgenes were utilized. **A-D:** Two hearts showing anti-Adprhl1 (green), mitosis marker anti-phospho-Histone H3 (red) and phalloidin actin stain (magenta). White squares (A, C) denote the position of detail images (B, D). Cells undergoing mitosis were readily detected within all regions of embryonic stage 40-41 hearts. There was no correlation between excess Adprhl1 production and cell division. Mitotic cells usually had no Adprhl1 signal (B) but occasionally did contain Adprhl1 protein (D). Note, a second P-H3-positive cell (A) near to the featured cell lay deeper within the myocardial wall so was not detected by the thin optical section of the high magnification image (B). Scale bars = 100 µm (A, C), = 10 µm (B, D).

**E-I:** Heart showing anti-Adprhl1 (green, E, F, H), ApopTag® TUNEL reaction stain (red, G, I), phalloidin (magenta, E, F, H) and DAPI (blue, F, H, I). The white square on the heart (Inset, E) locates images within the ventricle (F, G). Similarly, the squares (F, G) mark the position of detail images (H, I). The ApopTag® stain detects fragmented DNA of dying cells, caused by either apoptosis or necrotic destruction. The images focus on a cardiomyocyte on the ventricle anterior surface with excessive accumulation of Adprhl1 and a round appearance. Nevertheless, no ApopTag® signal was observed for this cell (G, I) nor indeed any Adprhl1-positive cells screened across 11 transgenic hearts. Scale bars = 100 µm (E, N), = 10 µm (F, G, J, K, O, P), = 5 µm (H, I, L, M, Q, R). **J-M:** Programmed cell death within the heart is a rare event during ventricle chamber outgrowth stages. Comparable images of a ventricle from a sibling transgenic tadpole (L has DAPI only). To prove the TUNEL reaction worked *in situ*, towards the apex, a solitary apoptotic cell was detected in an area without excessive Adprhl1 production (K, M). A typical condensed nucleus was observed with a fragmented DAPI stain (L) and intact ApopTag® signal (K, M). **N-R:** A further positive control for the ApopTag® detection method utilized a different responder transgene to induce necrotic cell death of the cardiomyocytes. The *Tg[UAS:M2(H37A)]* comprises part of a system for controllable genetic cell-ablation, producing the toxic viral ion channel M2(H37A) (Smith et al., 2007). The resulting small malformed heart was mounted with left side uppermost, with outflow tract to the left and ventricle to the right (inset, N). The images focus on the dying ventricle (O, P), in which numerous clusters of dots were present, positive for both ApopTag® fragmented DNA and DAPI (P-R). The scattered signals indicated cardiomyocytes had ruptured and that their cellular contents and degrading DNA had dispersed within the myocardium. V, ventricle; OT, outflow tract; A, apoptotic cell remnant; N, necrotic cell death fragments.

**Supplementary S4.** *Adprhl1 morpholinos - Position, sequence and activity in embryos.*
**A:** Diagram showing the hybridization position of MOs mapped to the first three exons of the S- and L-homeologous loci for *X. laevis adprhl1*. Morpholinos targeting potential translation initiation sites in *adprhl1* mRNA are shown by red arrows, with the corresponding methionine from the Adprhl1 protein sequence printed above. In addition to the 5'-most ATG predicted as the start for 40 kDa translation, there are five internal ATG sequences within the same reading frame. Morpholinos that target RNA-splicing (Smith et al., 2016) are coloured violet. **B:** Table containing the MO sequences and their complementarity to the S- and L-homeologs of *adprhl1*. Asterisks (*) denote sequence variability within *X. laevis* (see Supplementary Methods). Deliberately mismatched bases within control MOs are coloured blue. **C:** List showing the potential for hybridization to other members of the ADP-ribosylhydrolase gene family. Only MOs-S6 and L6 that target methionine-162 are noteworthy as they could cross-react with the four *adprh* loci that are present in *Xenopus* (*adprh.S, adprh.L, LOC495095, adprh-like.2.1.S*). **D:** The translation inhibition MOs produce varied effects on embryo development. Parts-of-whole charts showing the frequency of tadpole phenotypes assessed at stage 44 after 32 and 16 ng MO injection at the one-cell stage (percentage values listed for 32 ng injection). Heart defects were observed for the MOs that interfere with *adprhl1* RNA-splicing and for translation inhibiting MOs-L2 and S6. However, all three MOs designed to the most 5'-ATG (Met1), S1a, S1b and L1, caused unforeseen severe tail defects (after delayed blastopore closure at gastrulation). Morpholinos-3, 4 and 5 had no effect on embryo development. For MO-4 and 5, experiments are presented where a mixture of MOs targeting both S- and L-alleles was injected. Images showing representative embryos for active MOs are shown in Supplementary S5.

For the translation inhibiting MOs-L2 and S6 that did yield heart defects, both exhibited flaws that limit the MOs usefulness. Adprhl1-ATGMO2(L2) produced inert hearts with small ventricles and was designed to an internal ATG within exon 1 whose translation product would begin at Met26 (Supplementary S5H-K). However, this methionine is only found in the L-homeolog, with the MO being a poor match to the corresponding S-allele sequence. Thus S-allele function should be unaffected. Adprhl1-ATGMO6(S6) targeted an exon 3 sequence and potential translation from Met162 that was briefly considered as the start for the 23 kDa Adprhl1 protein. MO-S6 injection caused tadpole hearts that could contract but had failure of ventricle outgrowth (Supplementary S5L, M). Unfortunately, only the MO designed to the S-homeolog produced a heart phenotype whereas a preparation of -ATGMO6(L6) corresponding to the closely related L-allele sequence yielded distinct early developmental defects that precluded analysis of the heart (D).

**Supplementary S5.** *Adprhl1 morpholinos - Contrast between RNA-splicing versus translation inhibition.*
**A:** The diagram showing hybridization positions of MOs mapped to the first three exons of the S- and L-homeologous loci for *X. laevis adprhl1*. **B, C:** Reproduced for comparison, panels from Fig 1 featuring Adprhl1-e2i2MO RNA-splice interfering MO. Expression of *actc1* (heart and skeletal muscle, B) and *adprhl1* (C) mRNAs in stage 40 tadpoles after injection of 32 ng -e2i2MO. Impaired heart chamber growth and a loss of *adprhl1* mRNA signal is observed. Left-lateral view of tadpole and detail ventral view of heart region presented. **D, E:** Normal ventricle size and *adprhl1* signal in non-injected sibling tadpoles. **F-O:** Morpholinos designed to inhibit Adprhl1 protein translation produce varied effects on embryo development. Targeting distinct (but same reading frame) ATG-translation initiation sequences can yield malformations, but each exhibits a flaw that limits the MOs usefulness (see Supplementary S4 legend for details). **F, G:** Three overlapping MOs designed

to the 5’-most AUG of *adprhl1* mRNA each cause a tail growth defect, overshadowing any phenotype they might cause in the heart. Stage 39 tadpoles resulting from Adprhl1-ATGMO1(S1b) injection are shown. **H-K:** Adprhl1-ATGMO2(L2) injection results in inert hearts with small ventricles. Two severities of phenotype at stage 39 are shown, mildly affected tadpoles with small ventricles (H, J) and also details from tadpoles with a complete loss of ventricle growth (I, K). **L, M:** Adprhl1-ATGMO6(S6) gives some hearts that contract but have loss of chamber growth. **N, O:** Normal ventricle size and *adprhl1* signal in non-injected stage 39 sibling tadpoles. Red arrows denote aberrant morphology. H, heart; T, tail.

**Supplementary S6.** *Activity of distinct Cas9 RNAs and protein for tyrosinase gene knockout in X. laevis embryos.*
**A:** Comparing the coding sequence of four distinct Cas9 RNAs and the primary sequence of a commercial Cas9 protein preparation. The deduced Cas9 amino acid sequence originating from *Streptococcus pyogenes* is identical for all examples. Nevertheless, three shades of blue were used to depict Cas9 because the nucleotide sequences utilized differ due to distinct codon usage. Coding sequences were further modified by inclusion of epitope and purification tags (3xFLAG-green, V5-yellow, 6xHis-grey), nuclear localization signals (SV40-red, nucleoplasmin-magenta), self-cleaving 2A peptide (brown) and orange fluorescent protein reporter (orange). As an example of their size, the hSpCas9 protein is 1423 amino acids long, composed of N-terminal methionine, 22 aa 3xFLAG, 17 aa SV40-NLS, 1367 aa Cas9 (without N-term Met) and 16 aa nucleoplasmin-NLS sequences. **B:** Table showing the frequency of albino tadpoles obtained after disruption of the *tyrosinase* gene by injection of each Cas9 reagent together with *tyr* gRNAs into one-cell stage embryos (Materials and Methods). Parts-of-whole charts assigned the resulting stage 42 tadpoles to five albino phenotype classes that described the extent of pigmentation-loss and thus completeness of the gene knockout. **C:** Tadpoles representing the range of pigmentation-loss phenotypes observed after *tyrosinase* knockout. Left-lateral views, anterior half of tadpoles presented. Definitions of the five albino phenotype classes were comparable to those used by Guo *et al* (2014). **D:** Charts from a single experiment using Cas9 protein showing how the efficiency of *tyrosinase* knockout reduced as the time point of injection (minutes post-fertilization) increased. Beyond 90 minutes, embryos had reached the two-cell stage and thus injection of the same total mass of reagents was divided between both blastomeres. Based on this timed series, an upper limit of 60 minutes post-fertilization was set for all Cas9 injections at the one-cell stage.

**Supplementary S7.** *Adprhl1 gRNAs - Full list of gRNA experiments and embryo phenotype frequencies.*
Parts-of-whole charts showing the frequency of stage 44 tadpole phenotypes that occurred after injection of *adprhl1* gRNAs along with Cas9 protein into one-cell stage embryos. Red rectangles surround graphs for gRNAs whose activities were also examined by DNA sequencing. Green rectangles denote control gRNAs and also a graph presenting the cumulative total for all non-injected sibling tadpoles from the experiments. The highest frequencies of heart defects were detected using the gAdprhl1-e3-1 and in particular the -e6-1 gRNA. The lower seven panels show the consequence of combining gRNAs that hybridize to two genomic regions of *adprhl1* into a single injection. None of these combinatorial gRNA experiments increased heart defect frequencies beyond that obtained by gAdprhl1-e6-1 alone. Nonetheless, sequencing data was analysed for experiments where exons 3 and 4 were targeted at neighbouring positions to determine if resulting lesions contained deletions between the DSB sites (see Supplementary S11).

**Supplementary S8.** *Targeting adprhl1 exon 3 gives 100% mutation efficiency - S-homeolog DNA sequences.*

Sanger DNA sequences of *adprhl1* S-homeologous locus exon 3 after mutation by the gAdprhl1-e3-1(S+L) gRNA plus Cas9. Mutated sequences from the L-locus gave the same profile (data not shown). The hybridization position of the gRNA is depicted by the red arrow placed above the expected sequence (top 2 rows, exon and genomic). Alignment of 215 cloned (S-) isolates of amplified DNA obtained from 23 tadpoles, with every sequence carrying a lesion at the gRNA binding site. Mutant nucleotide sequences are coloured red. Missense mutations are listed first, followed by deletions (red hyphens, ordered by ascending size) and then sequences containing insertions (red arrowheads). Columns to the right give the number of instances of each sequence, alongside a genotype score that records the consequence of the given mutation to the Adprhl1 primary amino acid sequence. The key to interpret the genotype score is also included.

**Supplementary S9.** Exon 3 c*lassification of mutated adprhl1 sequences - Missense mutations are likely to retain function.*
**A:** All DNA sequences grouped by gRNA injection. Each DNA sequence of *adprhl1* exon 3 obtained after gRNA plus Cas9-mediated mutation was assigned a genotype score (a number code), based on the size of amino acid sequence modification it encoded. Parts-of-whole charts record the frequency of these genotype scores tallied for all embryos that received the -e3-1(S+L) gRNA (left chart), an embryo injected with a combination of neighbouring -e3-1(S+L) and -i2-2L gRNAs (centre chart), and a non-injected control embryo (right chart). The total number of S- and L-locus sequences analysed is listed below each chart. The key to interpret the genotype score is also included (far right). The inclusion of the -i2-2L gRNA induced larger deletions at the L-locus that would lead to exon 3 skipping, as shown by the increased proportion of score 02 sequences. **B:** Sequences and mutation details of individual embryos. Separate charts for 11 of the embryos, including 5 tadpoles that developed heart defects, 5 that developed normally and the one non-injected sibling. Columns list the number of sequences and the number of distinct sequences (in square brackets) for each embryo, plus the presence of larger deletions (and their size) that skip exon 3 and score 02. More importantly, for those embryos harbouring subtle mutations that scored 04 or 05, the precise amino acid change is also given. A standard nomenclature for protein sequence variations is used to describe the changes, with a simplified summary (highlighted) underneath stating whether the mutation caused a missense, deletion or aa insertion modification. Highlight colour matches the genotype score of the sequence. The rectangle border colour denotes a missense (teal), net deletion (blue) or net insertion (violet). At this position within exon 3, wherever missense mutations were found (eg p.(Lys145Ile)), the tadpole had preserved a normal heart morphology. **C:** Structure model of *X. laevis* Adprhl1 protein backbone, oriented with the ancestral active site cleft to the left and facing away (Arg271-Arg272 coloured red) (Smith et al., 2016). Exon 3 encodes a whole number of codons, contributing 41 amino acids arranged as two antiparallel α-helices. The extended loop mutated by the -e3-1(S+L) gRNA connects the two helices together and resides on the opposite face to the active site (coloured yellow and magenta).

Within exon 3, the consequence of in-frame, small amino acid deletions or insertions could not be defined with certainty. Nonetheless, triplet amino acid deletions were found in the normal cohort and may have protected the heart in two tadpoles (see tadpole#20170329014), which suggested these alleles were functional. Conversely, small insertions were biased towards animals with defective hearts, including examples where just a single amino acid was inserted between residues Lys145 and Pro146. Just one mutation gave apparently conflicting results. A 21 bp deletion resulting in a loss of 7 amino acids (p.(Met141_Gly147del)) was present in the heart defect group but also as the sole score 04-05 mutation detected in a normal tadpole (see tadpoles#20170406007, #20170406003). The model of Adprhl1 structure provided some context for the observed sequence modifications. It is plausible that a three amino acid deletion could be accommodated without loss of structural integrity (C).

**Supplementary S10.** *Targeting adprhl1 exon 4 with gAdprhl1-e4-1 gives 82% mutation efficiency - S- and L-homeolog DNA sequences.*
DNA sequences of *adprhl1* exon 4 after mutation by the gAdprhl1-e4-1 gRNA plus Cas9, with both S-homeologous locus (upper panel) and L-locus (lower panel) presented. The hybridization position of -e4-1 gRNA is shown by the red arrow and neighbouring gRNAs by grey arrows, placed above the expected sequence (top 2 rows, exon and genomic). Alignment of 93 DNA clones (S-63, L-30) of amplified DNA obtained from 5 tadpoles. 76 sequences were mutated but 17 remained unaltered, demonstrating this gRNA was less effective than those that targeted exons 3 and 6. As before, mutant nucleotide sequences are coloured red. Wild-type (WT) sequences are listed first, followed by missense mutations, deletions (red hyphens) and insertions (red arrowheads). The number of instances of each sequence is to the right, alongside its genotype score.

**Supplementary S11.** *Targeting adprhl1 exon 4 with adjacent gRNA pairs gives 88% mutation efficiency - S-homeolog DNA sequences.*
DNA sequences of *adprhl1* S-locus exon 4 after mutation by two adjacent gRNAs, gAdprhl1-e4-1 and -e4-2(S+L), plus Cas9. Sequences obtained from the L-locus gave the same profile (data not shown). Comparing data for both S- and L-loci from 4 tadpoles, of 95 DNA clones (S-68, L-27), 89 sequences were mutated but 6 S-sequences remained unaltered. Thus for exon 4 at least, combining pairs of gRNAs did not eradicate the low persistence of wild-type sequences. Some mutant sequences carried two separate lesions while others contained a single larger deletion between the two gRNA binding sites. As before, mutant nucleotide sequences are coloured red. Wild-type (WT) sequences are listed first, followed by missense mutations, deletions (red hyphens) and insertions (red arrowheads). The number of instances of each sequence is to the right, alongside its genotype score.

**Supplementary S12.** Exon 4 c*lassification of mutated adprhl1 sequences - Incomplete mutation rarely causes heart defects.*
**A:** All DNA sequences grouped by gRNA injection. Parts-of-whole charts record the frequency of sequence genotype scores tallied for all embryos that received Cas9 and gRNAs targeting *adprhl1* exon 4. The gAdprhl1-e4-1 gRNA was injected individually (far-left chart) and in combination with the neighbouring -e4-2(S+L) (centre-left chart) or -e4-3(S+L) gRNAs (centre-right chart), while sequences from non-injected control embryos were also compared (far-right chart). The total number of S- and L-locus sequences analysed is listed below each chart. The key to interpret the genotype score is also included. Mutation at exon 4 was extensive but incomplete and some wild-type sequences persisted for all combinations of gRNAs tested. **B:** Sequences and mutation details of individual embryos. Separate charts for 8 typical embryos with normal heart formation, including two representing each gRNA mixture plus sibling controls. Columns list the number of sequences and the number of distinct sequences (in square brackets) for each embryo, plus the presence of larger deletions (and their size) that skip exon 4 and genotype score 02. For smaller in-frame lesions, the precise amino acid change is also given. For these attempts to disrupt exon 4, most tadpoles retained a small proportion of wild-type sequence alleles. Of two presented here with 100% mutation, the reason their hearts were unaffected could not be determined conclusively. Nonetheless, one featured a subtle missense substitution while the second contained a deletion of 3 amino acids that would shorten an α-helix. They were also both notable for harbouring alleles likely to skip exon 4 from the mature mRNA. **C:** Structure model of *X. laevis* Adprhl1 protein backbone, oriented with the ancestral active site cleft to the left and facing away (Arg271-Arg272 coloured red). Exon 4 encodes a whole number of codons, contributing 47 amino acids. The start and end points of exon 4-derived amino acid sequence lie in close proximity within the model. Mutations

induced by the three exon 4 gRNA positions are dispersed (Gly172 green, Gln191 blue, His 213 magenta), but non reside on the same face as the active site.

**Supplementary S13.** *Targeting adprhl1 exon 6 causes near complete mutation and in-frame repair bias - L-homeolog DNA sequences.*
Cloned DNA sequences of *adprhl1* L-locus exon 6 after mutation by the gAdprhl1-e6-1 gRNA plus Cas9. Mutated sequences from the S-locus exon 6 are presented in Fig 4. Alignment of 249 (L-) DNA clones obtained from 16 tadpoles, with 248 carrying a lesion and just one instance of a wild-type (WT) sequence persisting. Mutant nucleotide sequences are coloured red. Deletion mutations are listed first (red hyphens) and then insertions (red arrowheads). Frequently occurring sequences containing in-frame 3, 9 or 12 bp deletions are highlighted. The number of instances of each sequence is to the right, alongside its genotype score.

Note, a handful of exon 6 clones had a mixed origin, containing S-sequence at one end and L-sequence at the other end of the PCR fragment. They must have been produced by two distinct PCR annealing and partial extension reactions occurring for a single DNA strand, exacerbated by the short extension times recommended by modern proof-reading DNA polymerases. They were assigned to S- or L- datasets according to the homeolog identity at the 5’- of the PCR. The first and nineteenth rows of this figure feature two such clones, classed as L-sequences but with 6 mismatches that have an S-origin.

**Supplementary S14.** *Xenopus Adprhl1 protein detection after exon 6 mutation.*
Western blot detection of Adprhl1 protein in stage 42-44 tadpole hearts after mutation of exon 6 using Cas9 plus the gAdprhl1-e6-1 gRNA. Pooled heart extracts of the -e6-1 sample came from animals with disrupted cardiogenesis. Despite the prevalent in-frame deletions, these hearts still produced both 40 and 23 kDa Adprhl1 similar to control hearts. Actin detection was used to normalize the samples.

**Supplementary S15.** *Range of ventricle phenotype severities observed after mutation of adprhl1 exon 6.*
After *adprhl1* mutation, tadpole heart phenotype was assessed at stage 44 and animals with aberrant morphologies were divided into two severity classes based on whether the ventricle was able to contract or not. The two examples here represent extremes from the range of hearts considered abnormal, from the most severely affected inert ventricle (A-H) to the mildest malformation observed in the beating ventricle class (I-P). It should be noted that most of the abnormal hearts were of an intermediate severity, were assigned to the beating group and resembled the earlier stage 40-42 examples shown in the principal figure (Fig 7), particularly with regard to the mosaicism found amongst the ventricular cardiomyocytes.

**A:** Strong phenotype. Cardiac oedema of a stage 44 tadpole after injection of the gAdprhl1-e6-1 gRNA plus Cas9. Right-lateral view of tadpole and left detail of small inert heart presented. Aside from the heart, there are no other discernible defects. Axial structures are straight and gut looping has commenced. **B, C:** Fluorescence images of the dissected heart placed with anterior-left surface uppermost showing phalloidin actin filament stain (green) scanned at the level of the ventricle myocardial wall (B) and a slice through the lumen (C) located 10 µm deeper. Ventricle growth has completely failed and no trabeculae ridges have formed at the inner surface of the chamber. **D-G:** Abnormal cardiomyocytes (D) with merged signals for phalloidin actin (green), anti-myosin (red) and DAPI nuclei (blue) from the region of ventricle wall framed by the white square (B). The white square (D) in turn marks the further magnified images (E-G) that show separate myosin and actin signals in addition to the channel merge. **H:** Muscle filaments inside a single cardiomyocyte identified by the open arrowhead (E-H). The cardiomyocytes vary with regard to the composition of their myofibril structure. Many retain a round shape and contain clusters of short, thin muscle filaments. These have primitive striated patterns to the myosin filaments but the actin strands are poorly defined with little periodicity (open arrowhead, E-H). Where cardiomyocytes have assembled longer myofibrils, the sarcomere spacing appears abnormal (filled arrowhead, E-G, also see below). Scale bars = 100 µm (B), = 10 µm (D) and = 5 µm (E). **I:** Mildest malformed phenotype after *adprhl1* mutation. Left-lateral view of stage 44 tadpole and ventral view of its small heart. **J, K:** The dissected heart ventricle with anterior surface uppermost scanned at the level of the myocardial wall (J) and a slice through the lumen (K) 8 µm deeper. The ventricle is small with a thin myocardial wall, but trabeculae ridges have formed correctly inside. **L-P:** Cardiomyocytes of the ventricle (L-O) and a myofibril detail within a single cell (P). A few round cells are present with prominent red myosin stain (open arrowhead, L) but most have produced myofibrils (filled arrowhead, M-P). At high magnification, the periodicity of the actin filaments in particular appears equally spaced rather than the characteristic striated pattern, with no brighter actin stripe indicative of a Z-disc (P). **Q-X:** Comparable images from a sibling non-injected control stage 44 tadpole. There is no oedema (Q), while the ventricle has the normal packing of elongated cardiomyocytes in the chamber wall (R, T) and parallel (base to apex) alignment of deep lying trabeculae (S). Within the myocardial wall, myofibrils extend predominantly in a perpendicular (to chamber) direction and their sarcomere repeats are mature with a clear Z-disc actin stripe (filled arrowhead, U-X). Oed, oedema; V, ventricle; OT, outflow tract; A, atria.

**Supplementary S1.**

Limited recovery of cardiac myofibril assembly in adprhl1 morpholino injected embryos by transgenic synthesis of recombinant 40 KDa Adprhl1 proteins

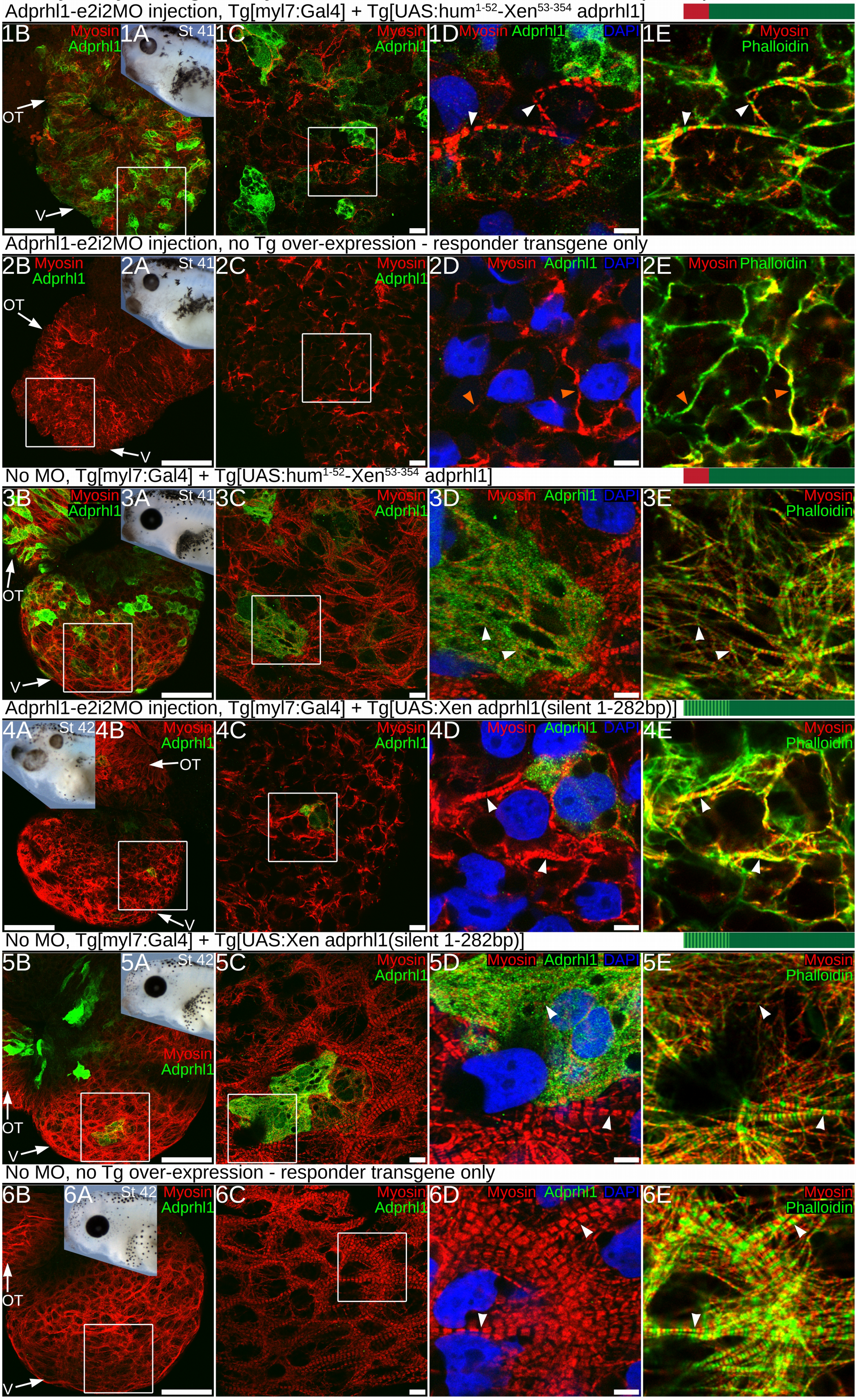

#### Supplementary S2.

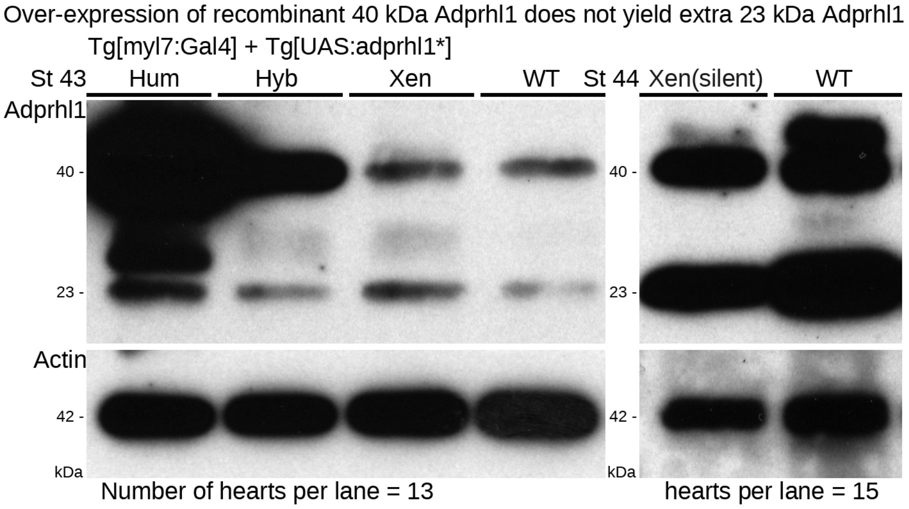

**Supplementary S3.**

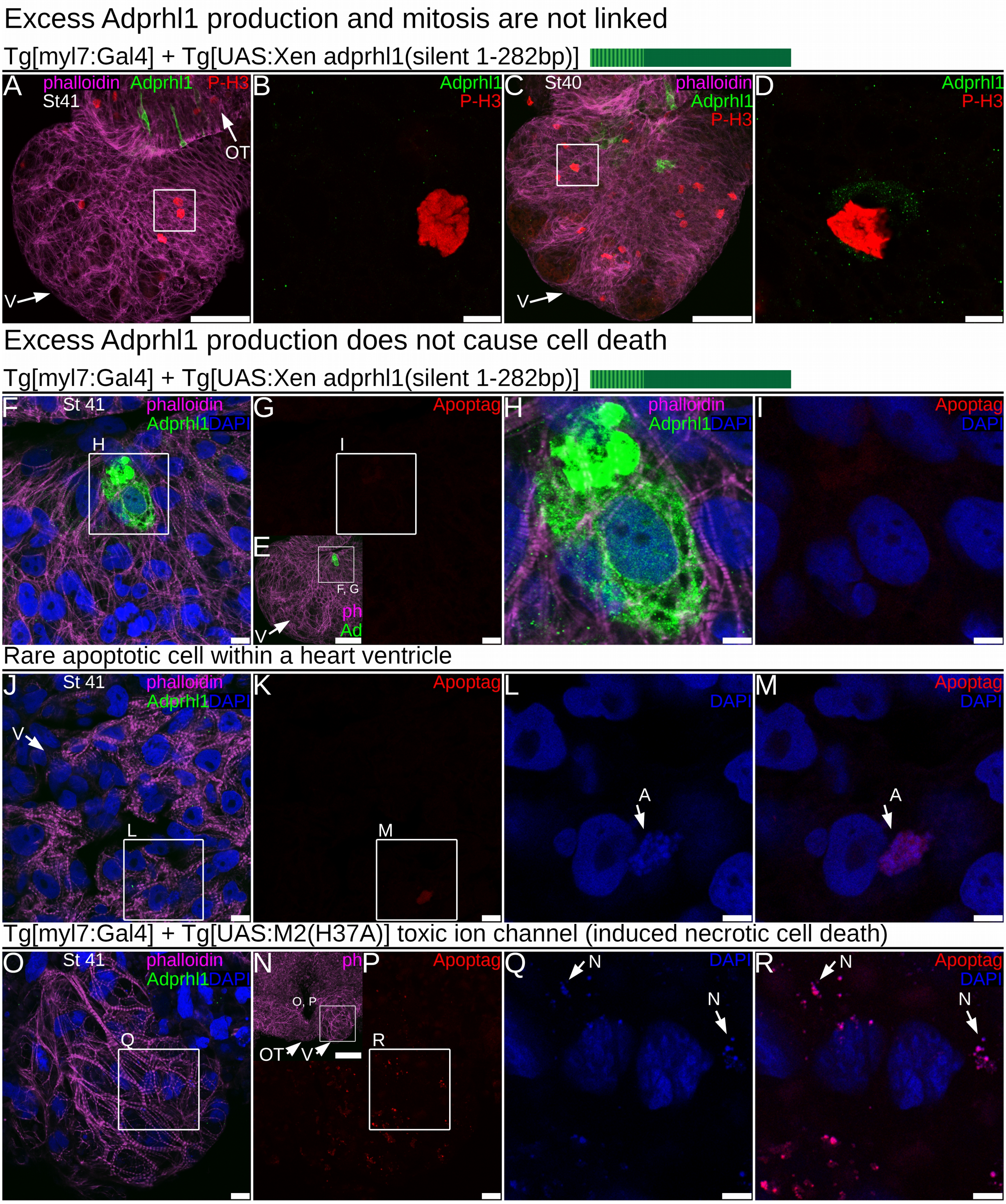

**Supplementary S4.**

#### *Adprhl1* morpholinos - Position, sequence and activity in embryos

A *X. laevis adprhl1* S-homeolog

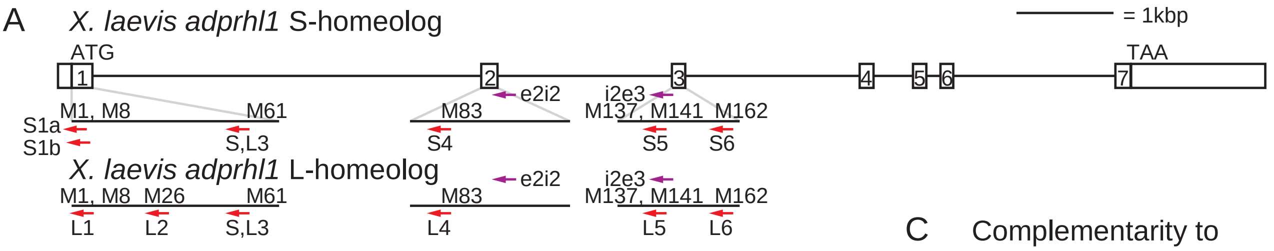


B Morpholino sequences

C Complementarity to ADP-ribosylhydrolase gene family members

| ATG | Morpholino | Sequence 5'-3' | Complementarity (n/25) S- | L-homeolog | adprh.S (n/25) | adprh.L | LOC-495095 | adprh-like.2.1.S | adprhl2.S | adprhl2.L |
|---|---|---|---|---|---|---|---|---|---|---|
| S1a | Adprhl1-ATGMO1(S1a) | CCTTAAACTTCTCCATAGCAAGGGC | 25 | 23 | | | | | | |
| S1b | Adprhl1-ATGMO1(S1b) | TGCAGCCTTAAACTTCTCCATAGCA | 25 | 22 | | | | | | |
| L1 | Adprhl1-ATGMO1(L1) | CATCGCAGCCTTAAACTTCTCCATG | 23 | 25 | | | | | | |
| L2 | Adprhl1-ATGMO2(L2) | GGATCTTTACTCCTGAAACACACAT | 19 | 25 | | | | | | |
| S,L3 | Adprhl1-ATGMO3(S,L3) | ATTAGAGTATTGTTACTCACTGGCC | 25 | 25 | 17 | 17 | 17 | x | 9 | 9 |
| S4 | Adprhl1-ATGMO4(S4) | GTTTTACCATGTCACGGTACAGGTC | 25 | 23 | | | | | | |
| L4 | Adprhl1-ATGMO4(L4) | GTTTCACCATGTCACGATACAGGTC | 23 | 25 | | | | | | |
| S5 | Adprhl1-ATGMO5(S5) | AGTATCTCATCCCAATACACATGGC | 25 | 24 | 19 | 19 | 19 | x | x | x |
| L5 | Adprhl1-ATGMO5(L5) | AGTATTTCATCCCAATACACATGGC | 24 | 25 | 18 | 18 | 18 | x | x | x |
| S6 | Adprhl1-ATGMO6(S6) | GATGGTTATGAGTCATTCTCCCACT | 25 | 24 | 23 | 21 | 20 | 20 | 12 | 13 |
| L6 | Adprhl1-ATGMO6(L6) | GATGGTTATGCGTCATTCTCCCACT | 24 | 25 | 22 | 20 | 20 | 20 | 12 | 13 |
| S6Mis | Adprhl1-ATGMO6Mis | GATAGTTATAAGTAATTCTACCAAT | 20 | 19 | 19 | 17 | 15 | 15 | 11 | 11 |
| **RNA-splice** | | | | | | | | | | |
| e2i2 | Adprhl1-e2i2MO | AGGCTCAGCATCTTACAAACCTTTT | 23 | 25* | | | | | | |
| e2i2Mis | Adprhl1-e2i2MOMis | AAGCTAAGCATATTAAAAACATTTT | 18 | 20 | | | | | | |
| i2e3 | Adprhl1-i2e3MO | ACCTAAGAAACAACTAGAGTCACTG | 23 | 25* | | | | | | |
| i2e3Mis | Adprhl1-i2e3MOMis | AACTAAAAAACAAATAGAATCAATG | 18 | 20 | | | | | | |

D Effect of morpholinos on embryo development

(injection at 1-cell stage)

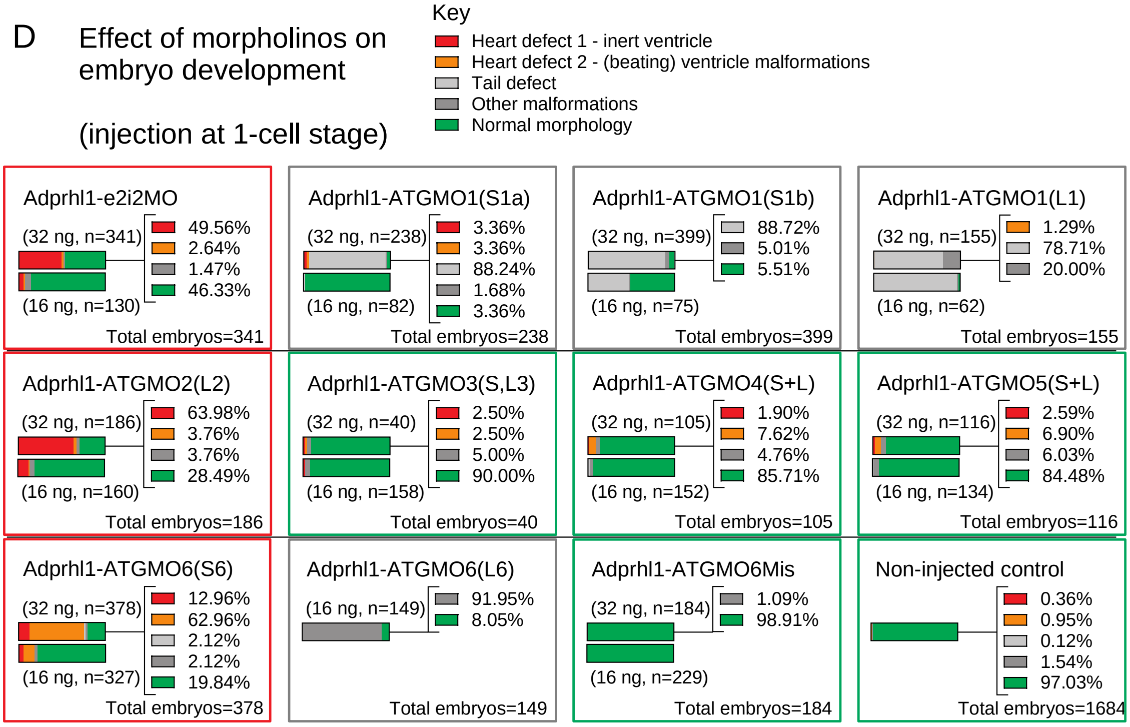

**Supplementary S5.**

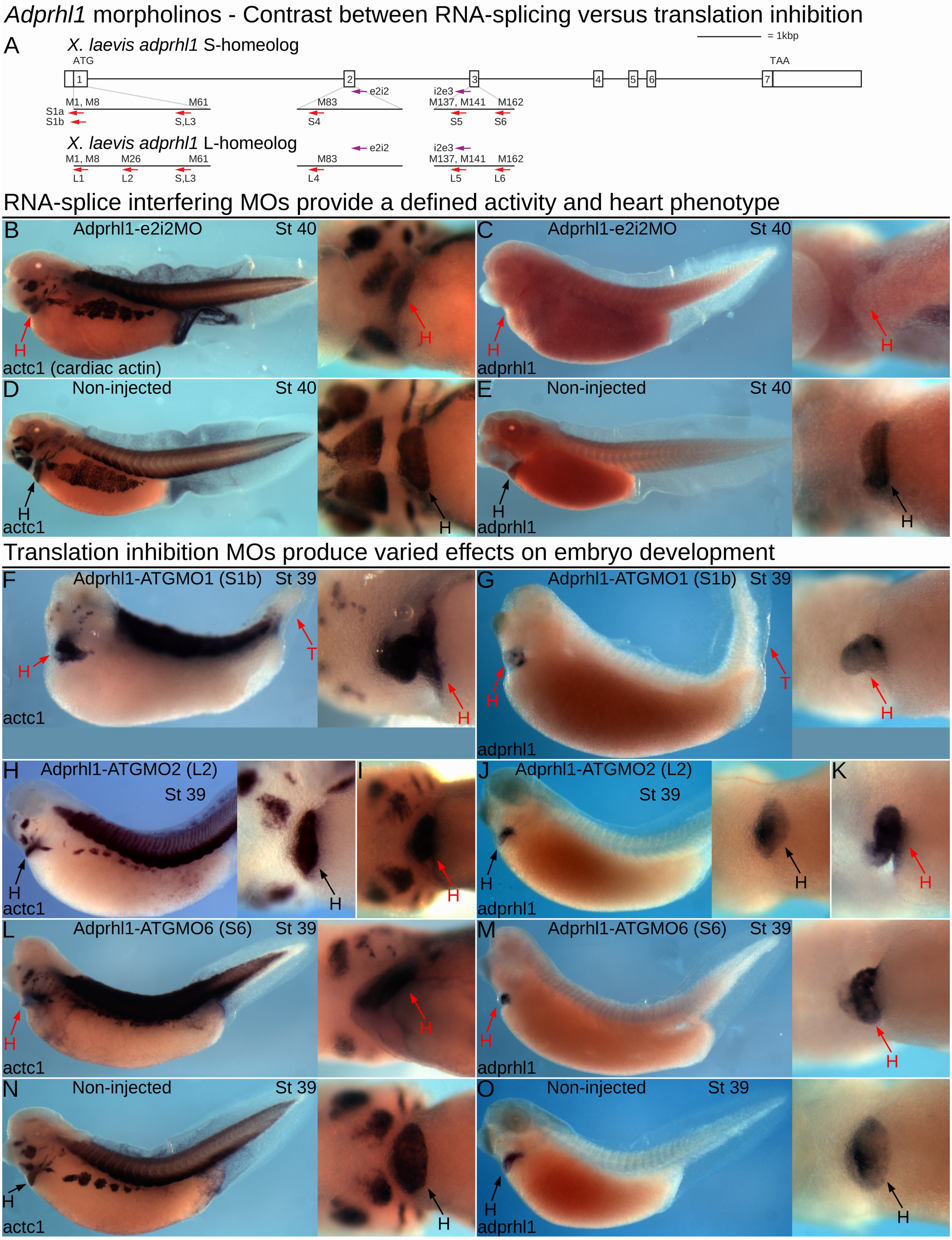

#### Supplementary S6.

##### Activity of distinct Cas9 RNAs and protein for *tyrosinase* gene knockout in *X. laevis* embryos

A

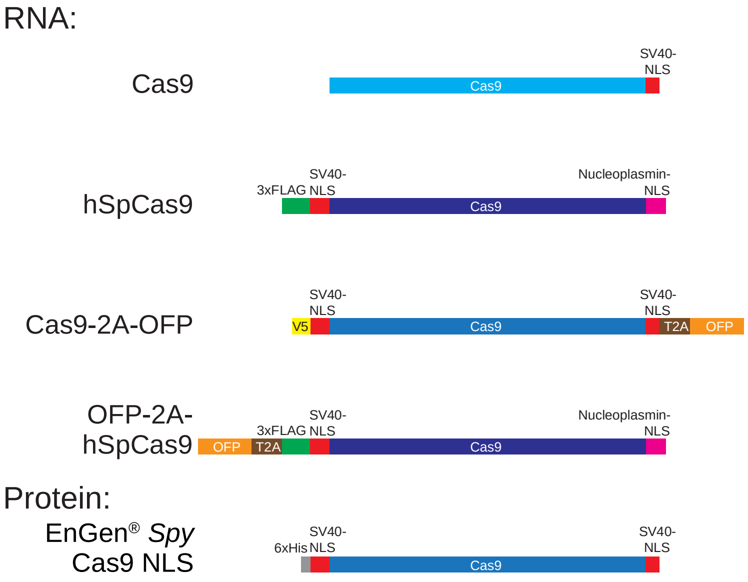


Cas9 (Addgene:47322) and Cas9-2A-OFP (GeneArt:A21174) share 77% Cas9 nucleotide sequence conservation

B

| Source | Cas9 Nucleotide Sequence Conservation | *Tyrosinase* gene knockout efficiency |
|---|---|---|
| Addgene: 47322 | 80% | 0.80%, 5.60%, 69.60%, 24.00%; Total embryos=125 |
| Addgene: 51815 | 100% | 24.24%, 22.51%, 43.94%, 3.25%, 6.06%; Total embryos=462 |
| GeneArt: A21174 | 86% | 6.28%, 69.96%, 12.11%, 11.66%; Total embryos=223 |
| Produced in house | 100% | 0.99%, 6.95%, 71.52%, 14.57%, 5.96%; Total embryos=302 |
| NEB: M0646M | NA | 72.92%, 8.33%, 17.71%, 1.04%; Total embryos=192 |

Key to albino severity

- Complete albino
- Nearly albino
- Partial albino 1
- Partial albino 2
- Pigmented

C Example albino tadpoles

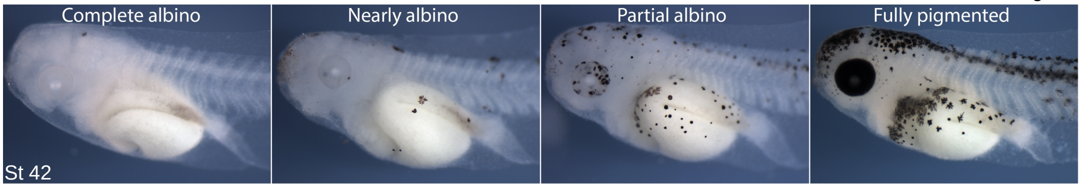


Class definitions are equivalent to Guo *et al* (2014) Development 141(3):707-14, figure 5O-R.

| | | |
|---|---|---|
| Complete albino: | 100% albino. | No pigmented cells present in stage 42 tadpole. |
| Nearly albino: | >90% albino. | No pigmented cells in the eyes (retinal pigment epithelium). Few pigmented cells (melanocytes) in the tail. |
| Partial albino 1: | ~51-90% albino | Some pigmented cells in the eyes and tail. |
| Partial albino 2: | ~5-50% albino | Mostly pigmented. Few bare patches visible within the eyes' pigment. |
| Fully pigmented: | 0% albino. | Fully pigmented. No bare patches visible within the eyes' pigment. |

The 'partial albino' definition encompasses a broad range of pigment-loss phenotypes and hence was divided into two groups.

D Timed series during a single injection experiment with Cas9 protein

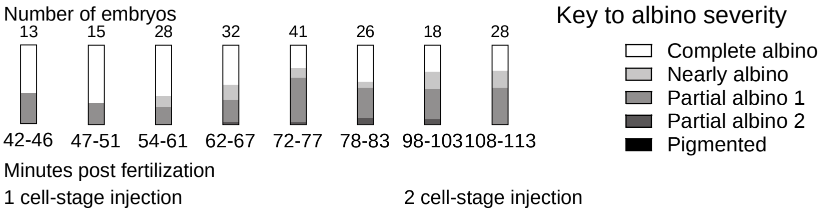

**Supplementary S7.**

#### *Adprhl1* gRNAs - Full list of gRNA experiments and embryo phenotype frequencies

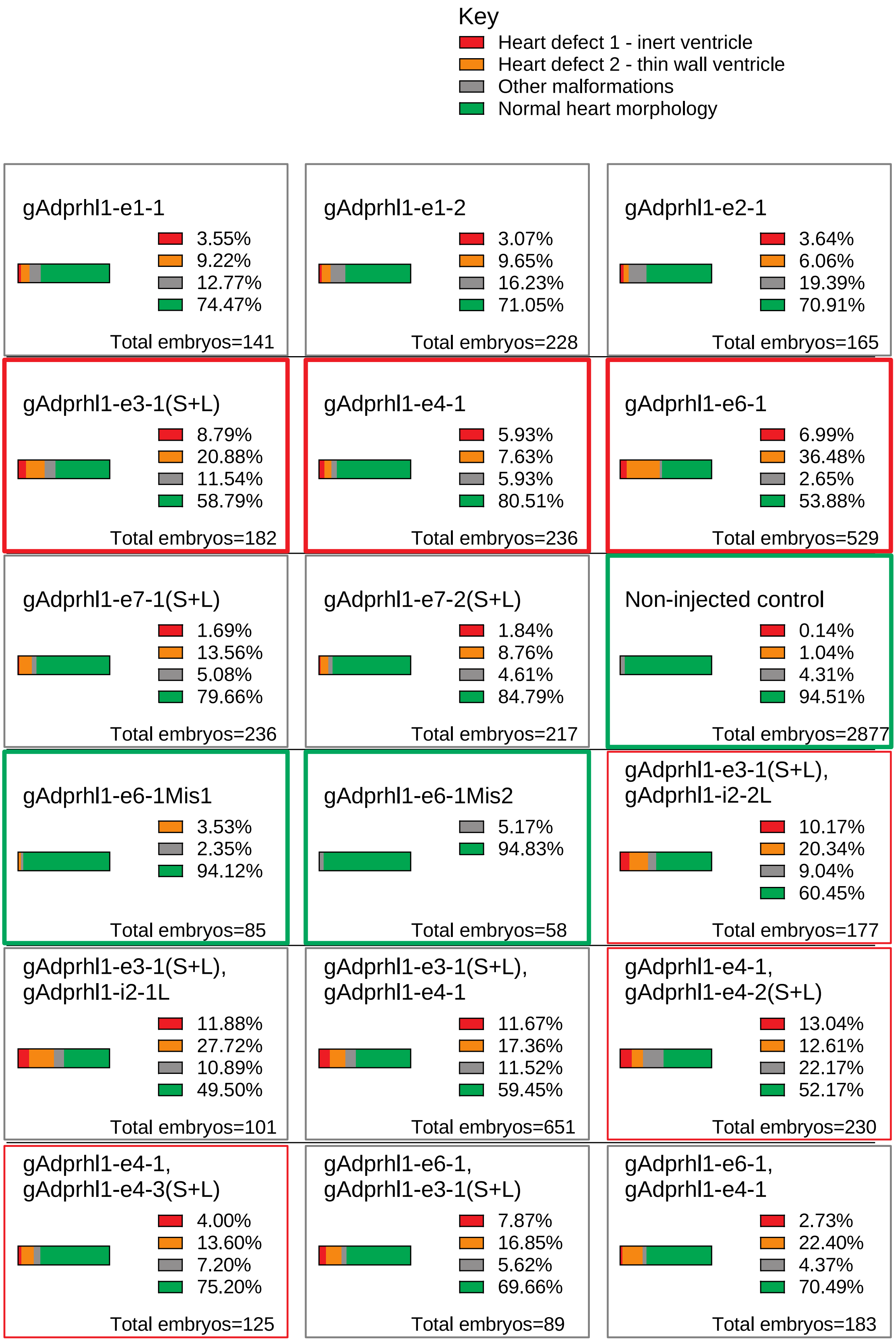

### Supplementary S8.

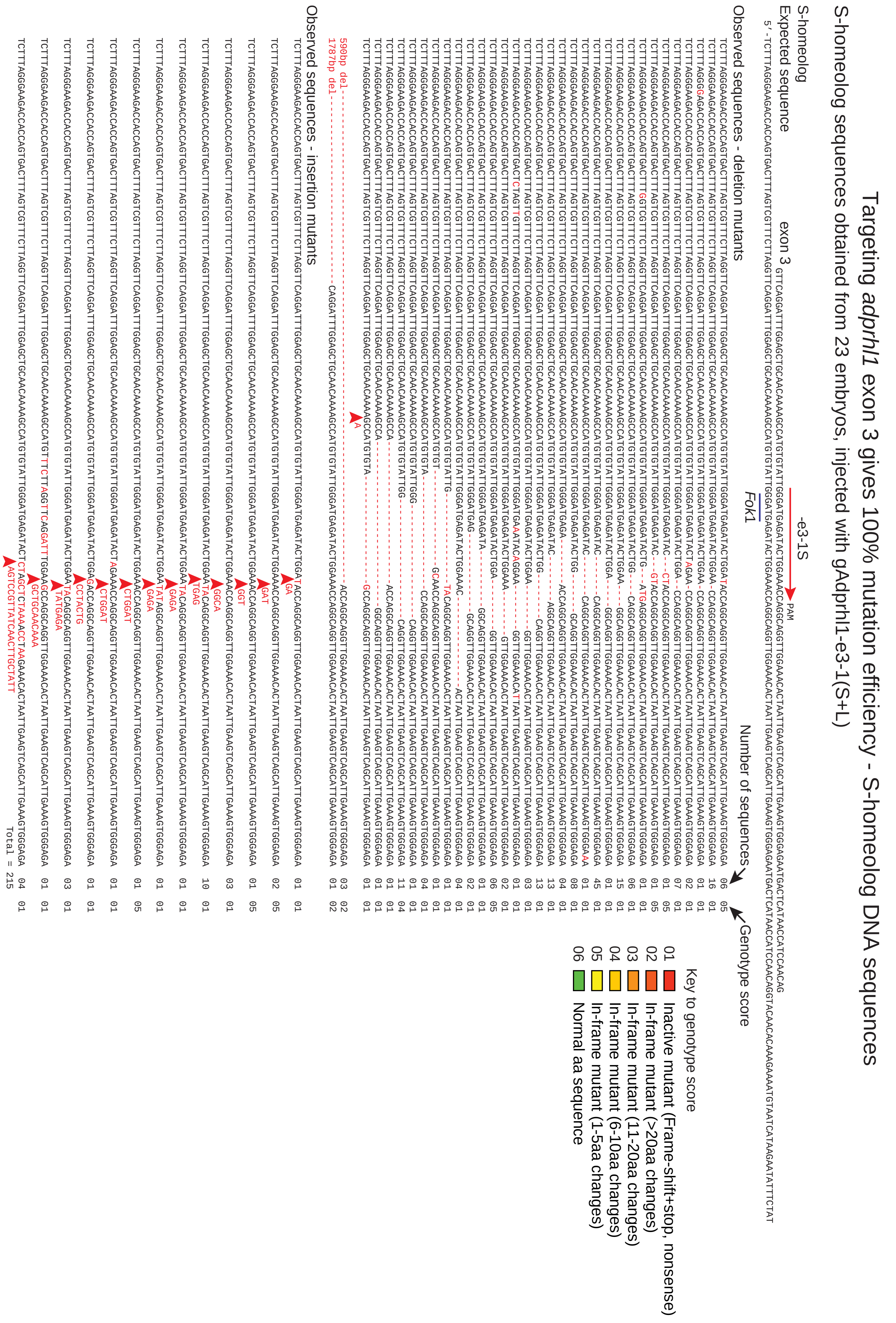

**Supplementary S9.**

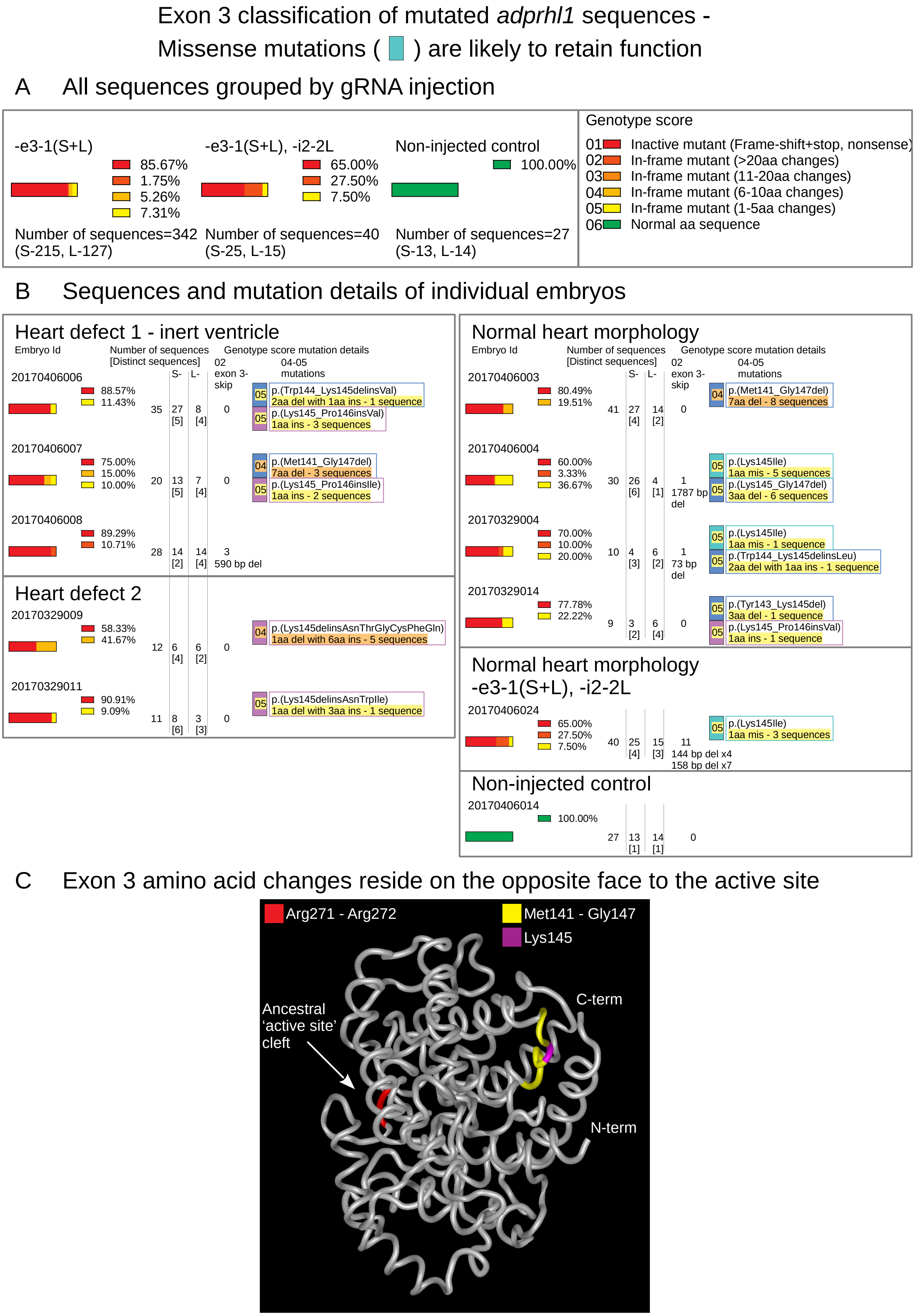

#### Supplementary S10.

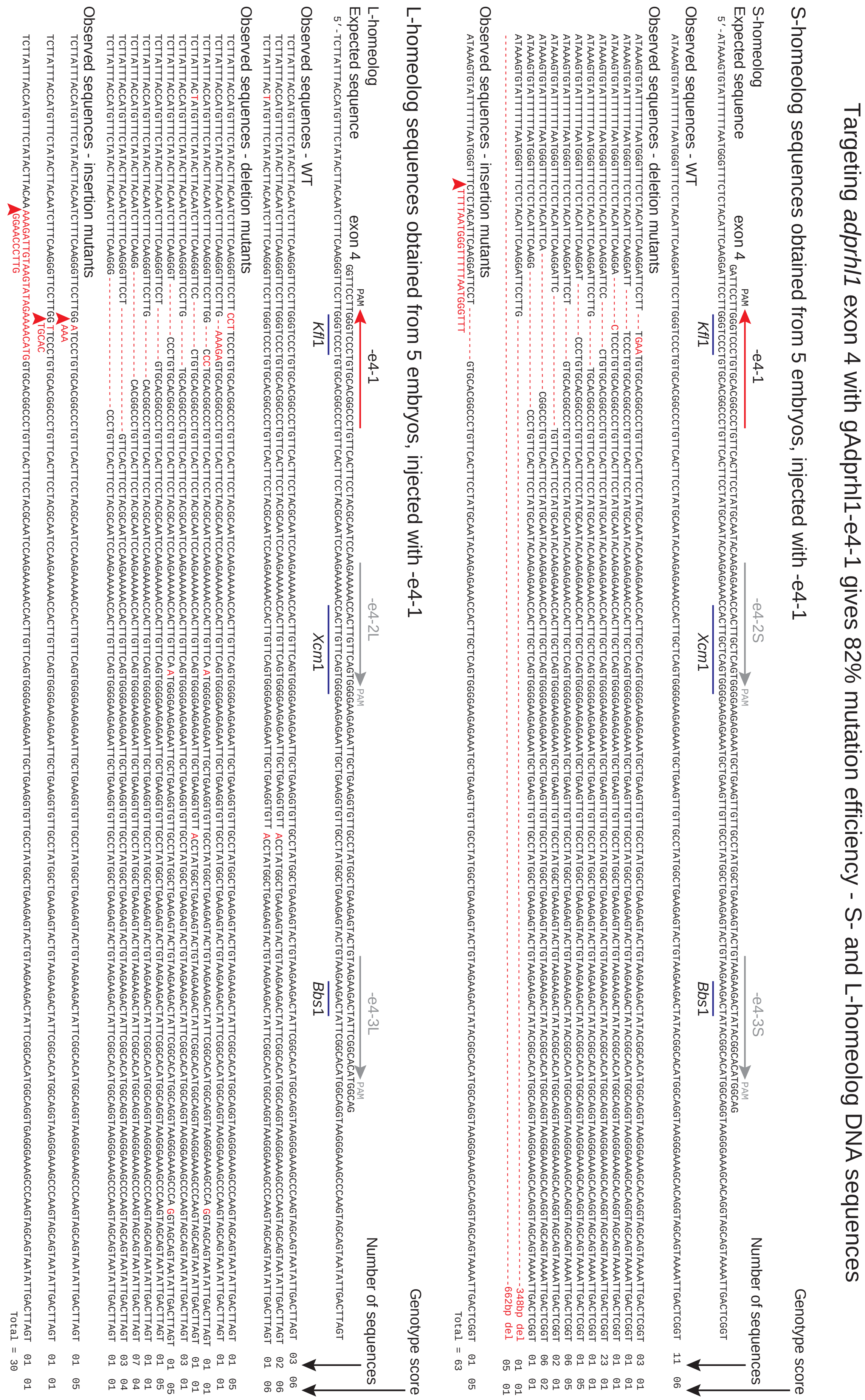

### Supplementary S11.

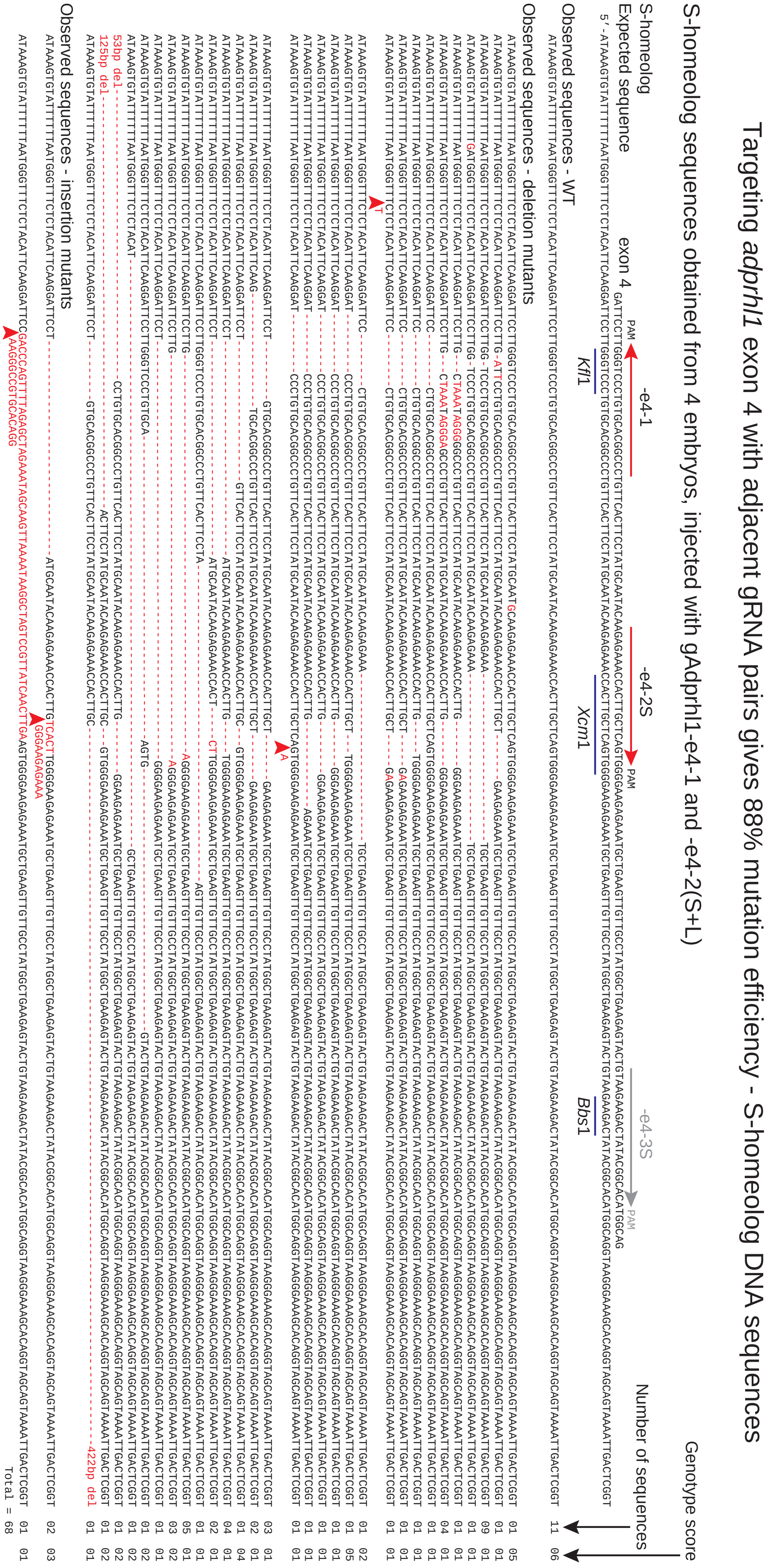

**Supplementary S12.**

### Exon 4 classification of mutated *adprhl1* sequences - Incomplete mutation rarely causes heart defects

Genotype score

- 01 Inactive mutant (Frame-shift+stop, nonsense)
- 02 In-frame mutant (>20aa changes)
- 03 In-frame mutant (11-20aa changes)
- 04 In-frame mutant (6-10aa changes)
- 05 In-frame mutant (1-5aa changes)
- 06 Normal aa sequence

#### A All sequences grouped by gRNA injection

**-e4-1**
51.61%
6.45%
10.75%
12.90%
18.28%
Number of sequences=93
(S-63, L-30)

**-e4-1, -e4-2(S+L)**
69.47%
10.53%
4.21%
2.11%
2.11%
11.58%
Number of sequences=95
(S-68, L-27)

**-e4-1, -e4-3(S+L)**
60.53%
13.16%
1.32%
5.26%
19.74%
Number of sequences=76
(S-58, L-18)

**Non-injected control**
1.32%
98.68%
Number of sequences=76
(S-52, L-24)

#### B Sequences and mutation details of individual embryos

##### gAdprhl1-e4-1
##### Normal heart morphology

| Embryo Id | | Number of sequences [Distinct sequences] | S- | L- | Genotype score mutation details: 02 exon 4-skip | In-frame mutations |
|---|---|---|---|---|---|---|
| 20170323015 | 43.33%<br>20.00%<br>23.33%<br>10.00%<br>3.33% | 30 | 21<br>[9] | 9<br>[3] | 6<br>23 bp del | 05 p.(Gly172Pro) 1aa mis - 1 sequence<br>05 p.(Gly172_Leu174del) 3aa del - 2 sequences<br>04 p.(Phe170_Cys175del) 6aa del - 7 sequences |
| 20170323016 | 52.17%<br>17.39%<br>30.43% | 23 | 18<br>[5] | 5<br>[2] | 0 | 05 p.(Gly172_Leu174del) 3aa del - 3 sequences<br>05 p.(Gly172delinsGluArg) 1aa del with 2aa ins - 1 sequence |

##### gAdprhl1-e4-1, -e4-3(S+L)
##### Normal heart morphology

| Embryo Id | | Number of sequences [Distinct sequences] | S- | L- | Genotype score mutation details: 02 exon 4-skip | In-frame mutations |
|---|---|---|---|---|---|---|
| 20170328075 | 71.88%<br>25.00%<br>3.13% | 32 | 20<br>[10] | 12<br>[6] | 8<br>17 bp del<br>23 bp del<br>23 bp del | 05 p.(Leu171_Gly172delinsGlnAsp) 2aa mis - 1 sequence |
| 20170328078 | 52.63%<br>10.53%<br>36.84% | 19 | 19<br>[10] | 0 | 0 | 05 p.(Gly172_Leu174del) 3aa del - 1 sequence<br>05 p.(Gly172_Leu174del, His213_-Met214delinsGln) 3aa del and 2aa del with 1aa ins - 1 sequence |

##### gAdprhl1-e4-1, -e4-2(S+L)
##### Normal heart morphology

| Embryo Id | | Number of sequences [Distinct sequences] | S- | L- | Genotype score mutation details: 02 exon 4-skip | In-frame mutations |
|---|---|---|---|---|---|---|
| 20170328044 | 79.31%<br>6.90%<br>13.79% | 29 | 21<br>[6] | 8<br>[4] | 1<br>256 bp del | 02 p.(Thr176_Glu205delinsLysTrp) 30aa del with 2aa ins - 1 sequence |
| 20170328048 | 63.33%<br>20.00%<br>6.67%<br>6.67%<br>3.33% | 30 | 22<br>[15] | 8<br>[6] | 2<br>87 bp del<br>125+ 3 bp del | 05 p.(Phe170_Gly172del, Gln191del) 3aa del and 1aa del - 1 sequence<br>04 p.(Gly172_Leu178del, Gln191del) 7aa del and 1aa del - 1 sequence<br>04 p.(Gly172_Leu174del, Leu189_-Gly193del) 3aa del and 5aa del - 1 sequence<br>03 p.(Gly172_Gln191delinsAlaArgProCys-SerLeuProThrGlnSerLysLysAsnHisLeu) 20aa del with 15aa ins - 2 sequences<br>02 p.(Gly172_Trp192delinsGlu) 21aa del with 1aa ins - 3 sequences<br>02 p.(Leu171_Gly193delinsProValHisGly-ProValHisPheLeuCysAsnThr) 23aa del with 12aa ins - 1 sequence |

##### Non-injected control
##### Normal heart morphology

| Embryo Id | | Number of sequences [Distinct sequences] | S- | L- | Genotype score mutation details: 02 exon 4-skip | In-frame mutations |
|---|---|---|---|---|---|---|
| 20170405021 | 100.00% | 17 | 11<br>[1] | 6<br>[1] | | |
| 20170405022 | 3.03%<br>96.97% | 33 | 23<br>[2] | 10<br>[1] | | 05 p.(Ala177Thr) 1aa mis - 1 sequence<br>(Natural sequence variation) |

#### C Exon 4 amino acid changes in α-helices not associated with the active site

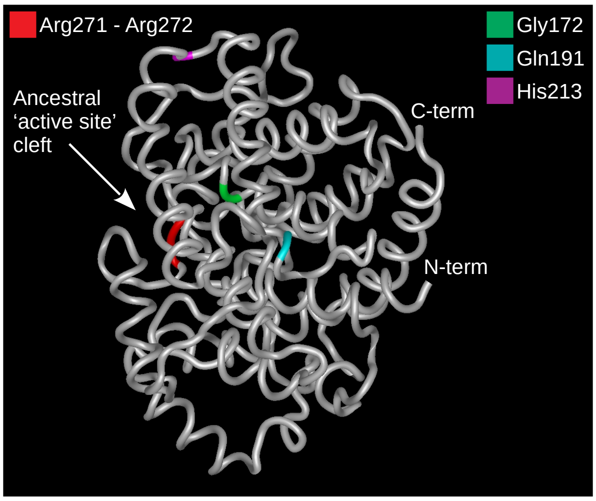

**Supplementary S13.**

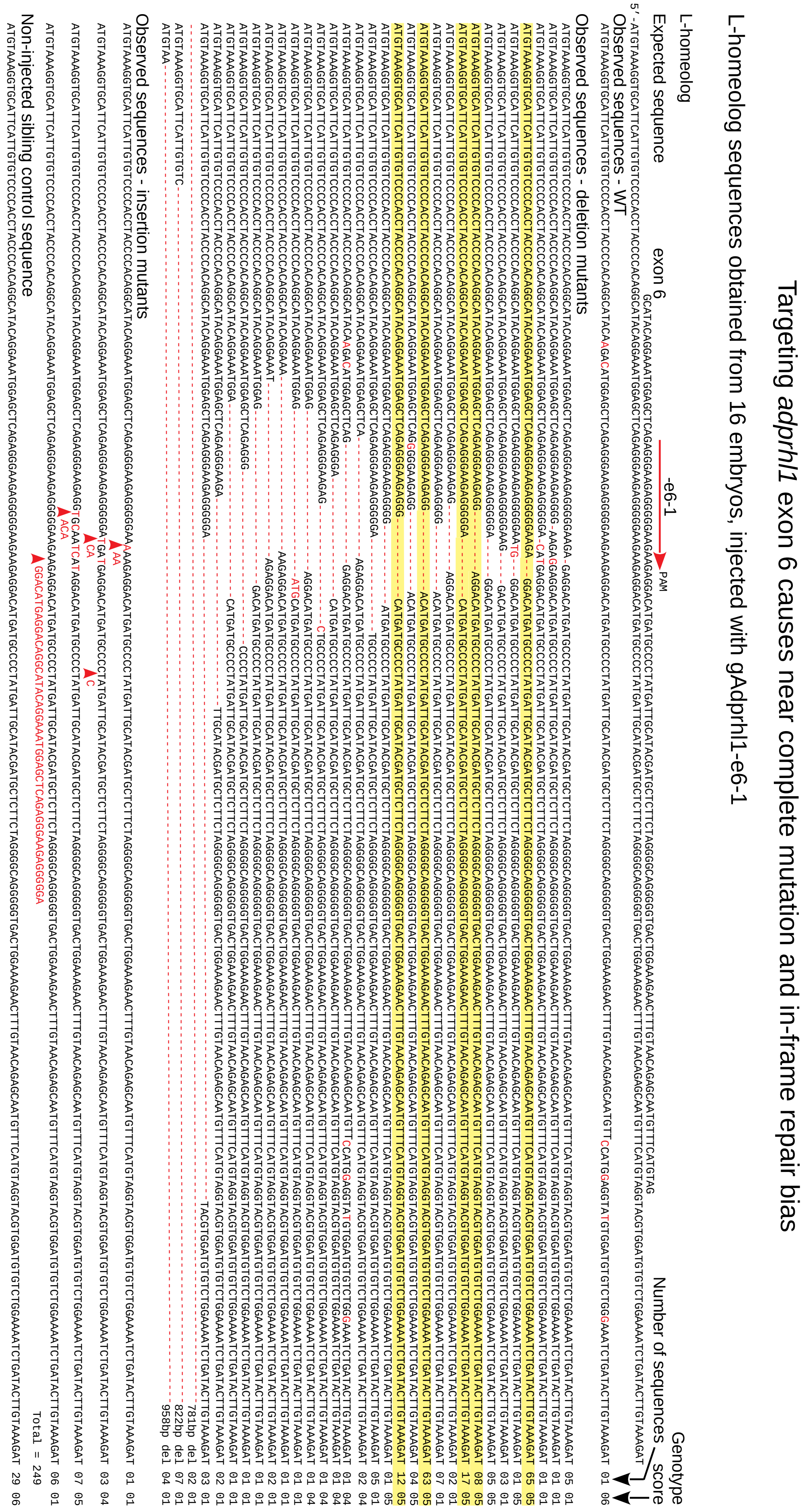

**Supplementary S14.**

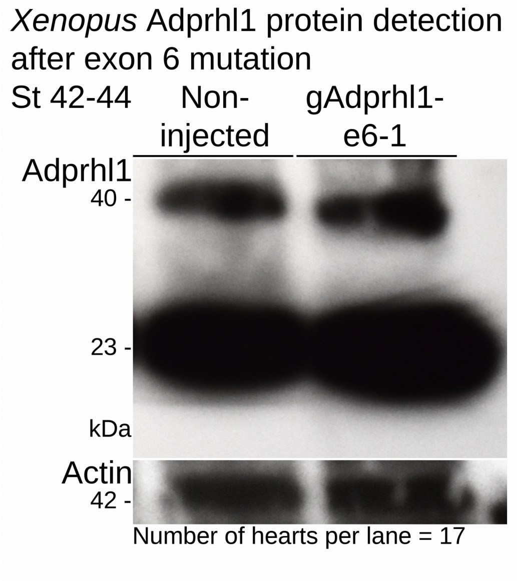

#### Supplementary S15.

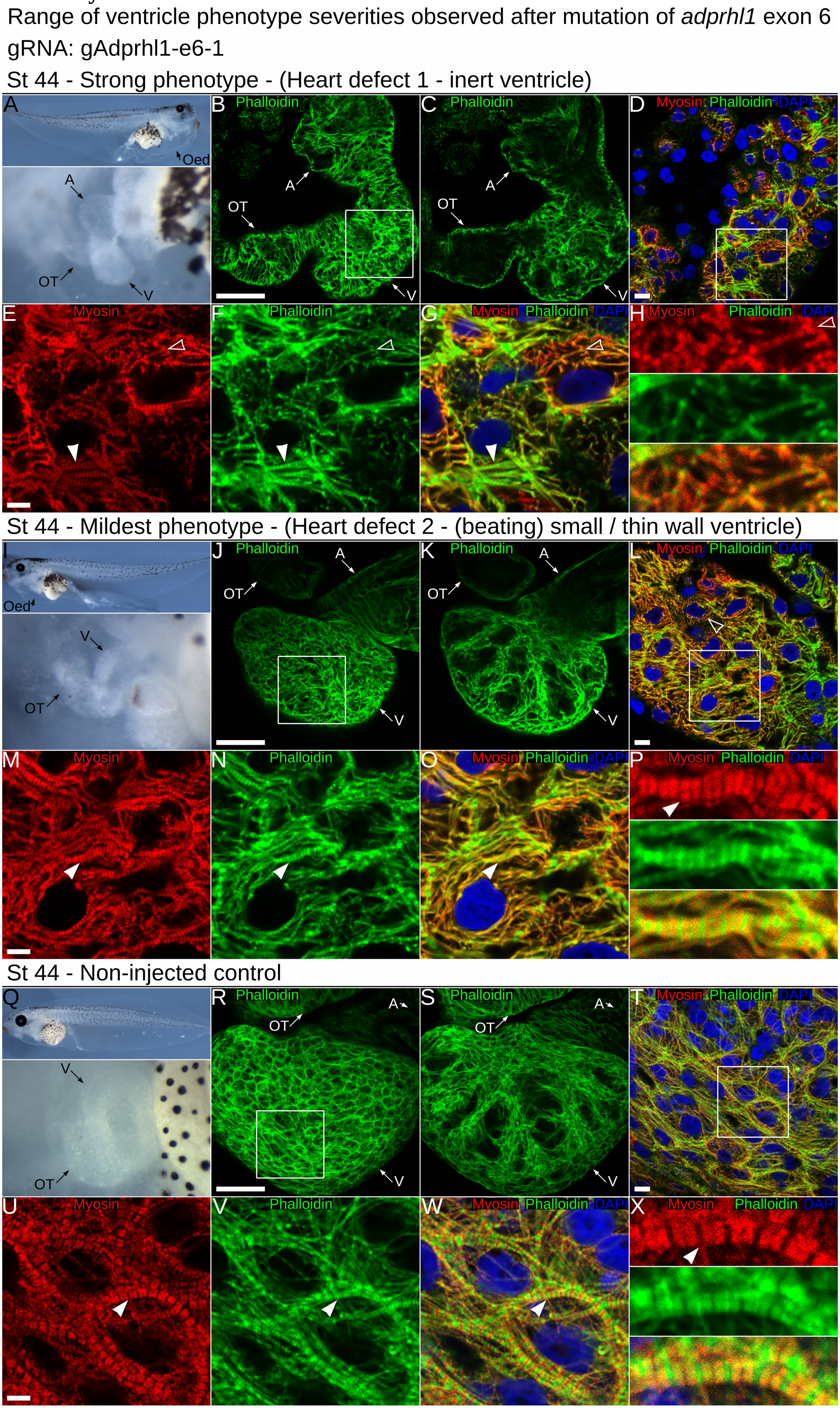

### Supplementary Methods.

*Background information on CRISPR/Cas9 optimization in Xenopus embryos.*
In aquatic vertebrate species like *Xenopus*, several approaches to targeted gene mutation are possible. The quickest is to supply Cas9 and gRNA reagents to newly fertilized embryos in sufficient quantity to cleave all DNA copies of the selected gene and instantly reveal mutant phenotypes as the injected embryos develop. A more traditional method would be to induce DSBs in a portion of the alleles, get germline transmission of a mutation and breed subsequent generations to obtain homozygous mutant animals. Gene mutations have been produced using both approaches in *X. tropicalis*, while the longer maturation time of *X. laevis* frogs and larger pseudo-tetraploid genome mean only the rapid, transient gene knockout experiments are practical. Nonetheless, the large and robust *X. laevis* embryo offers the experimental advantage of a longer time window for Cas9 to act prior to DNA synthesis occurring at the one-cell stage and has only four copies of a given gene that would require disrupting. Knockout phenotypes of well characterised genes have been obtained at higher frequency, and in greater number, using G0-generation *X. laevis* rather than *X. tropicalis* (Wang et al., 2015).

Initial studies in *Xenopus* supplied the endonuclease in the form of a Cas9 RNA. The RNAs sourced from different labs had distinct nucleotide sequences, some with codons optimised for use in mammalian species and the addition of one or two nuclear localization signals. There were obvious differences in efficacy of the Cas9 RNAs in *Xenopus*, with those utilizing RNAs with low *in situ* activity forced to compensate by increasing the mass of RNA injected (to unusual >2 ng quantities) and consequently experiencing problems with embryo toxicity (Bhattacharya et al., 2015). Injection of Cas9 protein has taken over from RNA (Naert et al., 2016). It allows a pre-incubation step to associate gRNA with the Cas9 that can facilitate instant activity once injected. With protein injection rather than Cas9 RNA, there is also lower risk of DNA damage from prolonged nuclease action beyond the first few cell divisions.

*Design of synthetic guide-RNAs.*
Selection of the *X. laevis adprhl1* gRNA sequences that were used in partnership with *S. pyogenes* Cas9 followed several design criteria. The entire exonic coding sequences of both S- and L-homeologous alleles were assessed. Where sequence differences existed between S- and L-alleles, separate gRNAs were prepared for each homeolog and then mixed prior to use. Only four sequences matched a $GGN_{18}$-(NGG) consensus that would enable *in vitro* transcription of gRNA from a plasmid template. Some additional gRNAs were selected (representing each *adprhl1* exon) that contained longer 21-22 nucleotides of gene-specific sequence. Ultimately, adopting PCR-based transcription templates gave greater freedom of gRNA sequence choice. It was possible to select $GN_{19}$-(NGG) sequences for gRNAs that target specific regions of interest within the *adprhl1* locus, notably those that encode the ancestral active site of the protein.

Current computational tools used to predict off-target activity of gRNAs have difficulty advancing beyond sequence alignment complementarity towards identifying those potential off-targets that are likely to be cleaved *in vivo* (Reviewed Wilson et al., 2018). Here, all gRNA sequences were assessed for hybridization at other positions using the JGI *X. laevis* genome v9.1 and Cas-Offinder (Bae et al., 2014), which considered the potential for both sequence mismatches and bulges. Results for potential off-targeting by the most active gAdprhl1-e6-1 gRNA to other gene coding sequences with up to two changes tolerated are listed below. None of the additional genes would be anticipated to contribute to a heart phenotype if mutated and more importantly, none of the potential interactions align with both of the homeologous alleles present in *X. laevis*.

| Gene | Exon | Expression | Mismatch | DNA bulge | RNA bulge. |
|---|---|---|---|---|---|

| | | | | | |
|---|---|---|---|---|---|
| nefm.L | 3 | Hindbrain, spinal cord | 0 | 0 | 2 bases near 5’-. |
| stac3.S | 2 | Skeletal muscle only | 1 (within seed) | 0 | 1 base near 5’-. |
| spryd3.S | 10 | brain | 1 (within seed) | 0 | 1 base near 5’-. |
| Xelaev18028958m.g | | Not determined (ND) | 1 (within seed) | 0 | 1 base near 5’-. |
| Xelaev18038829m.g | | ND | 1 (within seed) | 0 | 1 base near 5’-. |
| Xelaev18046191m.g | | ND | 1 (within seed) | 0 | 1 base near 5’-. |
| LOC100490289-like.L | | ND | 1 (within seed) | 0 | 1 base near 5’-. |
| LOC100497528.S | | ND | 1 (within seed) | 0 | 1 base near 5’-. |

*Transcription of guide-RNAs and Cas9-encoding RNAs.*

Two different methods were employed for synthesis of gRNAs. Where the gene-specific element of a gRNA began with a 5’-GG sequence, a plasmid template intermediate was produced. The pUC57-Simple-gRNA backbone plasmid (Addgene:51306, Yonglong Chen) (Guo et al., 2014) facilitated cloning of annealed complementary oligonucleotides with 4 base 5’-overhangs. Oligo-1: 5’-TA<u>G</u>G$N_{18}$ (transcription start site underlined). Oligo-2: 5’-AAAC$N^{rev\ comp}_{18}$. Plasmid DNA was linearized with *DraI* and used as template in a MEGAshortscript™ T7 transcription (Ambion). Transcribed gRNA was purified by ethanol precipitation and resuspended in 50 µl water.

Where the gRNA contained only a single 5’-G, a template DNA was prepared directly by PCR (Adapted from Naert et al., 2016). Long overlapping complementary oligonucleotides supplied all the necessary components: T7 RNA polymerase promoter, gene-specific and sgRNA sequences.
Oligo-1: 5’-GAAATTAATACGACTCACTATA<u>G</u>$N_{19}$GTTTTAGAGCTAGAAATAGC.
Oligo-2: 5’-AAAAGCACCGACTCGGTGCCACTTTTTCAAGTTGATAACGGACTAGCCTTATTTTAACTTGCTATTTCTAGCTCTAAAAC (this oligo universal to all templates).
The two oligonucleotides were annealed and a PCR-extension reaction performed using Platinum SuperFi DNA polymerase (Invitrogen), 50 µl reaction, 0.2 µM oligos, 25 cycles of 98 °C 10 sec, 55 °C 10 sec, 72 °C 10 sec. Template DNA was eluted through a QIAquick PCR purification column (QIAGEN) and added to a MEGAshortscript™ T7 transcription (Ambion). The transcription incubation time was increased to 5-7 hours to compensate for the suboptimal T7 promoter sequence.

Four different Cas9-encoding RNAs were tested for activity in *tyrosinase* gene knockout experiments, although Cas9 protein ultimately proved most effective in this assay (EnGen® Spy Cas9 NLS, NEB: M0646M). Two Cas9 RNA variants were sourced from Addgene plasmids, pCS2-Cas9 (Addgene:47322, Alex Schier) (Gagnon et al., 2014) and pCS2+hSpCas9 (Addgene:51815, Masato Kinoshita) (Ansai and Kinoshita, 2014). The Cas9-2A-OFP coding sequence (GeneArt:A21174) was purified using *NcoI-PmeI* sites and cloned into a modified pCS2. An OFP-2A-hSpCas9 sequence was prepared by cloning a synthesized OFP-2A- Strings DNA fragment (GeneArt) into pCS2+hSpCas9 using *BamHI-NcoI* sites. All plasmids were linearized with *NotI* and transcribed using mMessage mMachine™ SP6 reactions (Ambion). RNA was purified by LiCl precipitation and resuspended in 50 µl water.

*Tyrosinase gRNA sequences.*

Two gRNAs were previously designed against both *X. laevis tyrosinase* homeologous alleles (Wang et al., 2015). Their gene-specific sequences correspond to the *tyr* DNA listed below.

*tyra*-T: `5′-GGGTCGATGATAGAGAGGAC` (direction: →, PAM: `TGG`).
*tyrb*-T: `5′-GGCCCGTAGCAGAGCTGGTG` (direction: ←, PAM: `AGG`).

*Adprhl1 gRNA sequences.*

The *adprhl1* sequences used to design the principal gRNAs used in the study are listed below. Mismatched bases of control gRNAs are coloured red.

gAdprhl1-e3-1S: 5′-GGGATGAGATACTGGAAACC (direction: →, PAM: AGG).
gAdprhl1-e3-1L: 5′-GGGATGAAATACTGGAAACC (direction: →, PAM: AGG).
gAdprhl1-e4-1: 5′-GGGCCGTGCACAGGGACCCA (direction: ←, PAM: AGG).
gAdprhl1-e6-1: 5′-GAGGGAAGAGGGGGAAGAAG (direction: →, PAM: AGG).
gAdprhl1-e6-1Mis1: 5′-GAGGGAAGAGGGGGAACAAG (direction: →, PAM: AGG).
gAdprhl1-e6-1Mis2: 5′-GAGGGAAGAGGGGGAACTAG (direction: →, PAM: AGG).

Additional *adprhl1* gRNAs were also tested for activity, some with longer 21-22 nucleotides of gene-specific sequence. Mismatched 5'-bases added to enable efficient T7-transcription are coloured blue.

gAdprhl1-e1-1: 5′-GGCAAGAAGAGCTAAAACAACT (direction: →, PAM: TGG).
gAdprhl1-e1-2: 5′-GGTGAGACTAGTGATTCCGCTG (direction: ←, PAM: TGG).
gAdprhl1-e2-1: 5′-GGCACACCCCATTCAATGAAAA (direction: →, PAM: AGG).
gAdprhl1-i2-1L: 5′-GGAAAGCACTTAGTGACCAG (direction: ←, PAM: TGG).
gAdprhl1-i2-2L: 5′-GGGAATGTAATAAAAATTAG (direction: ←, PAM: TGG).
gAdprhl1-e4-2S: 5′-GGAGAAACCACTTGCTCAGTG (direction: →, PAM: GGG).
gAdprhl1-e4-2L: 5′-GGAAAAACCACTTGTTCAGTG (direction: →, PAM: GGG).
gAdprhl1-e4-3S: 5′-GGAAGAAGACTATACGGCACA (direction: →, PAM: TGG).
gAdprhl1-e4-3L: 5′-GGAAGAAGACTATTCGGCACA (direction: →, PAM: TGG).
gAdprhl1-e5-1S: 5′-GGTTTTATTTTGAAGCCAAG (direction: →, PAM: TGG).
gAdprhl1-e5-1L: 5′-GGTTTTATTTTGAAACCAAG (direction: →, PAM: TGG).
gAdprhl1-e7-1S: 5′-GCAGGAGAAGGTGGTGCCAC (direction: →, PAM: TGG).
gAdprhl1-e7-1L: 5′-GCAGGAGAAGGTGGTGCTAC (direction: →, PAM: TGG).
gAdprhl1-e7-2S: 5′-GGTGTCTTTATGGATTGCTCTA (direction: →, PAM: TGG).
gAdprhl1-e7-2L: 5′-GGTGTCTGTATGGATTGCTCTA (direction: →, PAM: TGG).

Note -e5-1 (S and L) fits the optimal gRNA consensus but gave consistently poor synthesis yields due to an adverse T-repeat sequence and could not be assessed by injection into embryos.

*Cas9 and gRNA injection into Xenopus laevis embryos.*

Preliminary experiments tested the activity of different Cas9 RNAs by co-injecting them with the two gRNAs that target *tyrosinase*. A mixture containing a Cas9 RNA (125 pg/nl) and the two gRNAs (both at 125 pg/nl) was prepared and 4 nl injected into one-cell stage embryos, giving a final 500 pg mass of each reagent. Injections were directed towards the animal pole of the embryo (uppermost third). Injection of embryos continued from 35 until 60 minutes post-fertilization. They were incubated at room temperature (22 °C) until 90 mpf, then transferred to 17 °C. Culture media used for injection and first 24 hours incubation was 0.1xNAM, 0.5 % Ficoll®-400, 20 µg/ml gentamycin. Thereafter, 0.1xNAM was used.

Greater efficiency of *tyrosinase* and *adprhl1* gene knockout was achieved using a commercial Cas9 protein preparation, EnGen® Spy Cas9 NLS (NEB: M0646M). In this case, gRNAs were preloaded onto Cas9 protein using a mixture assembled in the following order: 1 µl 1.3 M KCl (302 mM final), 2 µl gRNA (<1 µg of a single gRNA, so <233 pg/nl final) and 1.3 µl EnGen Cas9 (26 pmol). The mixture was incubated at 37 °C for 10 minutes, immediately prior to injection. Injection of 4 nl mixture and subsequent embryo culture was as before. The method was adapted from Burger *et al* (2016). For *tyrosinase* knockout, the phenotype classes used to define the extent of pigmentation-loss are described in Supplementary Figure S6 and follow those of Guo *et al* (2014).

For *adprhl1* knockout, embryos that gastrulated normally were allowed to develop to tadpole stage 44. Their external morphology was recorded each day and the appearance of their heart closely monitored. Tadpoles were assigned to one of four distinct phenotype classes, which differed slightly from those used to assess morpholino injection (see MO section, Supplementary Methods):
Heart defect 1 - inert ventricle. As per MO study.
Heart defect 2 - thin wall ventricle. In tadpoles showing a cardiac oedema that produced a beating heart, the ventricle was frequently thin-walled and became increasingly dilated by stage 44. This was especially true for embryos that received the exon 6 -e6-1 gRNA.
Other malformations. Any non-cardiac developmental defect visible externally by stage 44, however subtle.
Normal (heart) morphology. Perfect development through to stage 44.
There was no need to define a separate phenotype class for tail malformations as they did not occur in these CRISPR/Cas9 experiments (in contrast to Adprhl1-ATGMO1 morpholino injections).

*Sanger sequence analysis of mosaic adprhl1 exon 3, 4 and 6 mutations in G0-generation X. laevis.*
DNA was extracted from individual tadpoles. Frozen tissue was mixed with 200 µl 50 mM Tris pH8.8, 1 mM EDTA, 0.5% Tween20 containing freshly added 600 µg/ml proteinase K and incubated at 55 °C for 20 hours. PCR amplification of *adprhl1* genomic sequences used Platinum SuperFi DNA polymerase (Invitrogen) and 1 µl tadpole extract per 30 µl reaction. Plasmid clone isolates of amplicon DNA were prepared using a Zero Blunt™ TOPO™ PCR cloning kit (K287520, Invitrogen) and their inserts were sequenced. Sanger sequencing allowed study of 2 kbp amplicons so that the presence of larger deletions and insertions could be detected.

PCR primers for *adprhl1* genotyping.

| Exon 3 genotyping. | (position, direction). |
|---|---|
| p2483: 5′-TGCAAAGAGGGTTCTTTAGGGAAG | (intron 2, →). |
| p2460: 5′-GTCATTCTCCCACTTTCAATGCTGAC | (exon 3, ←). |
| p2555: 5′-TTGAAACCAGATAACTACCTG | (exon 2, →). |
| p2556: 5′-TCTCCCACTTTCAATGCTGAC | (exon 3, ←). |
| p2566: 5′-AGCCTCCCCGTATTCTCTAAG | (intron 2, →). |
| p2569: 5′-GAAATATTTATAGATTTTCATAAGGTGG | (intron 3, ←). |

PCR fragment sizes.
p2483+p2460, S-homeolog=161 bp, L=161 bp.
p2555+p2556, S=1944 bp , L=1326 bp.
p2566+p2569, S=426 bp.

| Exon 4 genotyping. | (position, direction). |
|---|---|
| p2515 5′-TCTTCTGCCATGAGACAAGGT | (intron 3, →). |
| p2517 5′-TCTTCCAGGTAAAACTGCCAC | (exon 5, ←). |
| p2541 5′-GAGCAATACCCAGAGTTTCTT | (intron 3, →). |
| p2542 5′-GAGCAATACCCAGAGGTGCTT | (intron 3, →). |

PCR fragment sizes.
p2515+p2517, S=876 bp.
p2541+p2517, S=893 bp.
p2542+p2517, L=944 bp.

| Exon 6 genotyping. | (position, direction). |
|---|---|
| p2560 5′-AGTTTTACCTGGAAGAAAGAG | (exon 5, →). |
| p2648 5′-GTGAGCTTATCTTTACAAGTATC | (intron 6, ←). |

p2649 `5′-TAAAGGGACACTTTCTTATCCAG` (intron 6, ←).
p2514 `5′-TCATCTCAAGCTGCTGGTATA` (exon 7, ←).
p2657 `5′-GGCTGGTCATGTGGCATGGTCA` (intron 6, ←).
PCR fragment sizes.
p2560+p2648, S=413 bp, L=404 bp.
p2560+p2649, S=604 bp, (L=592 bp).
p2560+p2514, S=2133 bp, L=1774 bp.
p2560+p2657, L=638 bp.

Each mutated sequence was assigned a genotype score (a number code) according to the size of the amino acid lesion that it encoded. This classification of mosaic mutations found within an individual tadpole was then compared against its cardiac morphology. Some assumptions were made for sequence deletions that disrupted exon splice junctions. Where a splice acceptor site was lost, it was assumed the exon was skipped. Where a splice donor sequence was removed, it was assumed the following intron was inappropriately retained.
Genotype score codes.
01: Inactive mutant. Frame-shift-stop or nonsense mutation.
02: In-frame mutant causing more than 20 amino acid changes.
03: In-frame mutant causing between 11-20 amino acid changes.
04: In-frame mutant causing 6-10 amino acid changes.
05: In-frame mutant causing 1-5 amino acid changes.
06: Normal amino acid sequence.

*Amplicon-EZ (NGS) sequence analysis of mosaic adprhl1 exon 6 mutations.*
The Amplicon-EZ next generation sequencing service (Genewiz) was used as a cost-effective method to obtain deeper coverage of the mosaic exon 6 mutations from individual tadpoles, providing 50,000 reads per sample. Genomic PCRs obtained with primers p2560+p2648 were sequenced directly using Illumina® technology. The Galaxy web platform (www.usegalaxy.org) was used to assemble sequence reads and combine duplicates (coalesce identical aligned reads) (Afgan et al., 2018). Reads were aligned to small 100 bp reference sequences of the *adprhl1* S- and L-alleles, covering 50 bp either side of the -e6-1 gRNA PAM. SeqMan Pro (DNASTAR) was also used to sort the sequences according to the variants found at the gRNA site and count the number of wild-type reads that occurred for each tadpole.

*Western blot detection of Adprhl1 protein.*
Adprhl1 protein was detected using a rabbit antibody raised against an ADPRHL1 peptide (mouse $^{248}$DNYDAEERDKTYKKWSSE$^{265}$, encoded by exons 5-6) (Smith et al., 2016). The antibody is active against the *Xenopus*, mouse and human species orthologs. For protein extraction from embryonic *Xenopus* hearts, typically 100 hearts were dissected, pooled, snap frozen, homogenized in 120 µl RIPA buffer and boiled with an equal volume of 2x reducing protein sample buffer. Individual adult mouse hearts were homogenized in 400 µl RIPA buffer, the resulting slurry mixed with 400 µl sample buffer, then aliquots diluted a further three-fold with RIPA/sample buffer before SDS-PAGE.

*Immunocytochemistry of Xenopus hearts.*
Immunocytochemistry was performed on whole tadpoles and subsequently the hearts were dissected, mounted in 12 µl CyGEL Sustain (biostatus) and viewed using Zeiss LSM5-Pascal or LSM710 confocal microscopes. Confocal images of whole hearts captured 2 µm deep optical sections. Both the ventricle myocardial wall and also deeper trabecular layers were assessed by scanning different depths. Images of cardiomyocytes and myofibrils were 1 µm optical sections, at a

depth 1-2 µm below the outer (apical) myocardial surface. Antibodies used were Adprhl1, Myosin A4.1025 (DSHB) and phospho-Histone H3(Ser10) 3H10 (Sigma), along with fluorescent dye-conjugated secondary antibodies (Jackson ImmunoResearch). Atto 633-conjugated phalloidin (Sigma) stained actin filaments. Dying cells were visualized with the ApopTag® Red In Situ Apoptosis Detection kit (Sigma).

*Adprhl1 morpholino sequences.*
Morpholino oligonucelotides (Gene-Tools) were previously designed to interfere with *X. laevis adprhl1* RNA-splicing (Smith et al., 2016) while new MOs aimed to inhibit the initiation of protein translation. For translation inhibition, MOs were synthesized that matched both S- and L-homeologous alleles and their activities were assessed both individually and mixed together. Furthermore, distinct MOs were designed to hybridize to every ATG-sequence within the first three exons that could initiate translation in the correct reading frame, in an attempt to identify the N-terminus of the 23 kDa Adprhl1 protein species.

For the RNA-splice interfering MOs, the advent of the improved JGI *X. laevis* genome v9.1 has revealed some sequence variability. Nonetheless, the two MOs remain valid for targeting both S- and L-alleles (Supplementary S4. Note, previous RT-PCR analysis would have successfully detected both homeologous alleles (Smith et al., 2016)). For Adprhl1-e2i2MO, approximately half of the L-alleles sequenced are a perfect match to the MO while half contain two base mismatches (GenBank accessons GU188989, GU188990 have matching sequence, JGI genome v9.1 J-strain allele has mismatches at MO bases 6 and 15). For Adprhl1-i2e3MO, half the L-alleles sequenced match the MO and half contain a single mismatch (J-strain allele has the matching sequence, GU188989 has a mismatch at MO base 25).

Sequences.
Adprhl1-ATGMO1(S1a): 5′-CCTTAAACTTCTCCATAGCAAGGGC-3′.
Adprhl1-ATGMO1(S1b): 5′-TGCAGCCTTAAACTTCTCCATAGCA-3′.
Adprhl1-ATGMO1(L1): 5′-CATCGCAGCCTTAAACTTCTCCATG-3′.
Adprhl1-ATGMO2(L2): 5′-GGATCTTTACTCCTGAAACACACAT-3′.
Adprhl1-ATGMO3(S,L3): 5′-ATTAGAGTATTGTTACTCACTGGCC-3′.
Adprhl1-ATGMO4(S4): 5′-GTTTTACCATGTCACGGTACAGGTC-3′.
Adprhl1-ATGMO4(L4): 5′-GTTTCACCATGTCACGATACAGGTC-3′.
Adprhl1-ATGMO5(S5): 5′-AGTATCTCATCCCAATACACATGGC-3′.
Adprhl1-ATGMO5(L5): 5′-AGTATTTCATCCCAATACACATGGC-3′.
Adprhl1-ATGMO6(S6): 5′-GATGGTTATGAGTCATTCTCCCACT-3′.
Adprhl1-ATGMO6(L6): 5′-GATGGTTATGCGTCATTCTCCCACT-3′.
Adprhl1-ATGMO6Mis: 5′-GATAGTTATAAGTAATTCTACCAAT-3′.
Adprhl1-e2i2MO: 5′-AGGCTCAGCATCTTACAAACCTTTT-3′.
Adprhl1-e2i2MOMis: 5′-AAGCTAAGCATATTAAAAACATTTT-3′.
Adprhl1-i2e3MO: 5′-ACCTAAGAAACAACTAGAGTCACTG-3′.
Adprhl1-i2e3MOMis: 5′-AACTAAAAAACAAATAGAATCAATG-3′.

Aside from the described sequence similarity between ATGMO6 and the ADP-ribosylhydrolase gene family, BLAST searches did not indicate any potential MO hybridization to other *Xenopus* mRNAs or genomic regions that could cause off-target activity. Nonetheless, all potential interactions are listed below.

| MO | Aligns with | Complementarity | Strand (MO seq). |
|---|---|---|---|
| L2 | *Xelaev18011653m*, exon 2 | 20/25 | sense (+) strand. |

| | | | |
|---|---|---|---|
| L2 | *ago2.L*, intron 1 | 20 | + strand. |
| S6 | *LOC108699836*, intron 14 | 20 | - strand. |
| e2i2MOMis | *irp10.L*, exon 5 | 21 | - strand 3'-UTR. |
| i2e3MOMis | *gda.L*, intron 13 | 20 | - strand. |

*Morpholino injection into Xenopus laevis embryos.*
When the Adprhl1-e2i2MO morpholino was combined with stable transgenic *adprhl1* over-expression lines, embryos were injected into both dorsal blastomeres at the four-cell stage (Smith et al., 2016). In all other experiments, MOs were injected at the one-cell stage in order to provide a better comparison with the CRISPR/Cas9 *adprhl1* gene knockout. The RNA-splice interfering MOs gave a slightly reduced effectiveness with one-cell stage injection because it results in a lower relative concentration within heart forming tissue. Two masses of MO were assessed, 32 ng and 16 ng. All embryos that gastrulated normally were allowed to develop to tadpole stage 44. Their external morphology was recorded each day.

Tadpoles were assigned to one of five distinct phenotype classes:
Heart defect 1 - inert ventricle. First visible sign of aberrant morphology at stage 40-41, with developing cardiac oedema. Heart remains small, string-like. The cardiac pacemaker initiates pulsing contractions but the small ventricle region remains inert.
Heart defect 2 - (beating) ventricle malformations. A broader class of cardiac phenotypes. First sign of aberrant morphology again at stage 40-41, with developing cardiac oedema. The forming ventricle does propagate a heart beat but is malformed. The most common malformation is a small ventricle. Other cardiac malformations observed include a thin-walled ventricle, becoming increasingly dilated by stage 44, or a ventricle with incorrect left-right and antero-posterior orientation.
Tail defect. First sign of aberrant morphology at stage 32, showing a twist in the trunk region and a loss of tail outgrowth. Tail defect becoming more extreme by stage 44. Heart malformation frequently observed but difficult to ascertain whether a primary defect or a consequence of failed development elsewhere.
Other malformations. This class mostly found with MO-L6. Normal gastrulation but subsequently develop severe malformations by stage 24.
Normal morphology. Perfect development through to stage 44.

*Binary transgene system for cardiac Adprhl1 expression.*
The design of transgenes for cardiac over-expression of Adprhl1 proteins has been described previously (Smith et al., 2016). The full name of the principal transgenes used here are: *Tg[myl7:Gal4, γCrys:eCFP]*, *Tg[UAS:human$^{1-52}$-Xenopus$^{53-354}$ adprhl1, γCrys:DsRed1]* and *Tg[UAS:Xenopus adprhl1(silent 1-282bp), γCrys:DsRed1]*. Additional responder transgenic lines featured as controls for western blot detection of recombinant proteins. The *X. laevis adprhl1* cDNA was from IMAGE:4409193 and corresponded to the S-homeologous allele sequence. The 5’-cDNA of human *ADPRHL1* was from IMAGE:5299214. The *X. laevis adprh* S-allele cDNA used for an *in situ* hybridization probe was from IMAGE:6953995.

*Binary transgene system for controllable genetic cell ablation.*
The *Tg[UAS:M2(H37A), γCrys:DsRed1]* line utilized the toxic mutant version (H37A) DNA of the *influenza* virus ion channel protein M2 (Smith et al., 2007). The plasmid clone was produced exactly as for *adprhl1* responder transgenes.
